# Toss GERALT into chloroplast to make it green: an age-dependent regulator of chloroplast biogenesis and chlorophyll biosynthesis

**DOI:** 10.1101/2024.08.16.608208

**Authors:** Agnieszka Katarzyna Banaś, Joanna Grzyb, Piotr Zgłobicki, Andrzej Pacak, Beata Myśliwa-Kurdziel, Katarzyna Leja, Małgorzata Kozieradzka-Kiszkurno, Kinga Kłodawska, Robert Konieczny, Maria Pilarska, Ewa Niewiadomska, Aneta Bażant, Łukasz Szewc, Ewa B. Kowalska, Wojciech Strzałka

## Abstract

Proteins belonging into cryptochrome/photolyase family act as either blue light photoreceptors or as enzymes responsible for repair of pyrimidine dimers produced under UV. Surprisingly, the members of plant-specific photolyase/blue-light receptor 2 (PPL/PHR2) subclade of this family, are almost completely uncharacterized. Here, we focused on the physiological role of protein encoded by Arabidopsis *At2g47590* gene. Our results demonstrate for the first time its crucial role in cotyledon greening and maintenance of plant fitness. Based on the phenotype of the mutant in *At2g47590* gene i.e. albino cotyledons and pale green true leaves we named this gene *GERALT* (*<u>GER</u>MINATION <u>AL</u>BINO <u>T</u>RANSIENT).* Using different approach including analysis of plant phenotypes, chloroplast sizes and architecture, transcriptomes, photosynthetic pigments, maximum PSII quantum yield (F_V_/F_M_) we show that the proper plant functioning is the effect of co-operation of GERALT-dependent and -independent pathways with the role of the former diminishing with plant age. Lower levels of transcripts dependent on plastid encoded polymerase and higher levels of these dependent on nuclear encoded polymerase, smaller chloroplasts with large grana stacks and very weakly developed stromal thylakoids, lower levels of photosynthetic pigments with higher chlorophyll a/b ratio, are among characteristic features of *geralt* plants. We believe that these results encourage scientific community to study PPL/PHR2 proteins which seems to play a special role in plants.

## 1. Introduction

Light is one of the most important factors controlling plant life. On the one hand, it is a driving force of photosynthesis. On the other hand, it serves to integrate plant functioning according to the environmental conditions. Plant life cycle, starting from germination, through the growth of seedlings and adult plants to the flowering and senescence, is modulated by light quality and quantity. In addition to retrograde chloroplast to nucleus signals originating from photosynthesis, light is sensed by specialized photoreceptors. UVB is perceived by UVR8 using the tryptophan triad for photoreception (Rizzini et al., 2011). The specified wavelengths absorbed by other photoreceptors depend on the chromophore bound. Phototropins, cryptochromes (CRYs) and F-box containing flavin binding proteins (e.g., ZEITLUPE (ZTL), FLAVIN BINDING, KELCH REPEAT, F-BOX1 (FKF1) and LOV KELCH PROTEIN2 (LKP2)) are flavoproteins acting as UVA/blue light photoreceptors. Red/far red is sensed by bilin-binding phytochromes (Paik and Huq, 2019).

Cryptochromes form huge protein complexes enabling integration of blue light and other internal and external signals. They not only regulate the expression of genes involved in germination, photomorphogenesis and flowering but are also crucial elements of a plant circadian clock (Wang and Lin, 2020). In the dark, CRYs exist as monomers and undergo oligomerization under blue light. Blue light regulates also interaction of CRY complexes with other proteins. Cryptochromes together with photolyases belong to a cryptochrome/photolyase family (CPF). Photolyases carry two non-covalently bound chromophores. In addition to the flavin adenine dinucleotide (FAD) also found in cryptochromes which acts as a catalytic chromophore, they bind an antenna chromophore: either pterin (methenyltetrahydrofolate (MTHF)) or deazaflavin (8-hydroxy-7,8-didemethyl-5-deazariboflavin (8-HDF)) (Sancar 2003). Photolyases directly bind to UV-induced DNA lesions i.e. cyclobutane pyrimidine dimers (CPDs), or (6–4) pyrimidine–pyrimidone photoproducts (6–4 PPs). This binding is independent on light, however they need UVA/blue light energy to split a bond between neighboring pyrimidines in a process called photoreactivation. Although it is a matter of debate whether the common ancestor of CPF was an iron–sulfur cluster containing 6-4 P- or a CPD-specific photolyase there is no doubt that such protein appeared early during life evolution (Lucas-Lledó and Lynch, 2009; Xu et al., 2021). Throughout the history of CPF proteins there have been many gain-and-lose events that have resulted in the emergence of cryptochromes, proteins that have lost the ability of DNA repair but are able to sense blue light. This scenario is confirmed by the presence of proteins that combine both functionalities. Some *Drosophila, Arabidopsis, Synechocystis*, human-type cryptochromes (CRY-DASHs) that repair, at least *in vitro*, CPDs only in a single stranded DNA, function as blue light photoreceptors and transcription regulators (Kiontke et al., 2020).

Based on phylogenetic analysis CPF proteins in higher plants can be classified into five subfamilies: i) 6-4 photolyases, ii) class II CPD photolyases, iii) plant cryptochromes, iv) *Drosophila, Arabidopsis, Synechocystis*, human (DASH)-type cryptochromes (CRY-DASHs) and v) plant photolyases/photolyase/blue-light receptor 2 (PPL/PHR2) (Mei and Dvornyk, 2015; Deppisch et al., 2022). Seven genes encoding CPF proteins are found in Arabidopsis genome. AtCRY1 and AtCRY2 are canonical cryptochromes. AtCRY1 functions in cytoplasm and nucleus, AtCRY2 seems to be solely a nuclear protein (Wang and Lin 2020). In turn, AtCRY3 is classified as a member of a CRY-DASH subclade. This protein is localized in chloroplasts and mitochondria and, at least *in vitro*, can bind and repair CPDs in single stranded DNA (Kleine et al., 2003; Pokorny et al., 2008). The role of AtCRY3 *in vivo* has not yet been elucidated. Photoreactivation in Arabidopsis is performed by two photolyases: CPD-specific AtPHR1/AtUVR2 (PHOTOREACTIVATING ENZYME 1, AtUVR2 -UV RESISTANCE 2) and 6-4 PP specific AtUVR3 (UV RESISTANCE 3)(Jiang et al. 1997). There are no data available for the two remaining Arabidopsis CPF proteins, At4g25290 and At2g47590/AtPHR2 (PHOTOLYASE/BLUE-LIGHT RECEPTOR 2).

In this work we focused on the second one, being the member of PPL/PHR2 subfamily of CPF proteins. This plant specific subfamily is characterized by unique features such as a truncated FAD binding domain (Mei and Dvornyk, 2015; Deppisch et al., 2022). The physiological role of PPL/PHR2 proteins, already present in *Chlorophyta* is almost completely unknown. The papers by Mei and Dvornyk (Mei and Dvornyk, 2015) and Deppish et al. (Deppisch et al., 2022) report that PHR2/PPL proteins have CPD photolyase activity based on the work of Petersen et al. 1999 (Petersen et al., 1999) concerning the *Chlamydomonas reinhardtii* PHR2 protein with annotation in the gene bank AAD39433.1/AF129458_1. In the phylogenetic trees, however, this protein localizes within the CPDII subfamily of CPF proteins (Mei and Dvornyk 2015; Deppisch et al. 2022). In contrast, the PPL/PHR2 subfamily includes the still uncharacterized *Chlamydomonas reinhardtii* protein 12g534550v5/XP_042918714.1. To gain knowledge about this mysterious group of proteins, we characterized the Arabidopsis mutant in *At2g47590/AtPHR2* gene. We show that such plants have albino cotyledons and pale green true leaves. Adult plants are dwarf and their growth is delayed as even over 5 month-old mutants still produce inflorescences and seeds. Since *PHR2* symbol is also used for a plant gene involved in the response to phosphate starvation (which stands for *PHOSPHATE STARVATION RESPONSE2*) we propose to rename *At2g47590/AtPHR2* with *<u>GER</u>MINATION <u>AL</u>BINO <u>T</u>RANSIENT* (*GERALT*), a name that describes the mutant phenotype.

In our research, we characterized the efect of a mutation in the *GERALT* gene on the seedling development. Two life stages were chosen for more detailed analysis, 5 day-old and 12 day-old seedlings having only albino cotyledons or with the first true leaves, when the mutants start to turn green, respectively. The transcriptomes, chloroplast ultrastructures, photosynthetic efficiencies, levels of photosynthetic pigments in these plants and WT seedlings of the same age were compared. Whereas the amounts of selected plastid encoded RNA polymerase (PEP)-dependent chloroplast transcripts are lower in the *geralt* mutant the steady-state levels of selected nuclear encoded RNA polymerase (NEP)-dependent transcripts are similar or higher than in wild type (WT) plants. With the visible greening that progresses with the age of the mutant plants, an increase in photosynthetic quantum yield is observed, but even in adult plants this parameter is clearly lower than in WT ones. At the both life stages size and shape of *geralt* chloroplasts and the architecture of their thylakoids are disturbed. GFP-tagged GERALT protein is localized in the stromal fraction of chloroplasts and in stromules.

All obtained results suggest that GERALT acts neither as a photolyase nor blue light photoreceptor, cryptochrome, but as a modulator of chloroplast functions. Its role diminishes with plant age.

## 2. Results

### 2.1. Overview of the GERALT gene and the T-DNA mutant line used

*GERALT* (*At2g47590*) gene consists of four exons and three introns (Supplemental Fig. S1A). The encoded protein with molecular weight of 48.9 kDa is made up of 447 amino acids. According to InterPro 97.0 database (Paysan-Lafosse et al., 2023) two main domains, i.e. Crypto_Photolyase_N_sf (IPR036155) and Crypto/Photolyase_FAD-like (IPR036134) are predicted in GERALT protein (Fig. 1A). However, FAD binding domain is notably shorter as compared with other members of CPF (Fig. 1A, (Mei and Dvornyk, 2015; Deppisch et al., 2022)) what may be later reflected in its function.

**Fig. 1.**
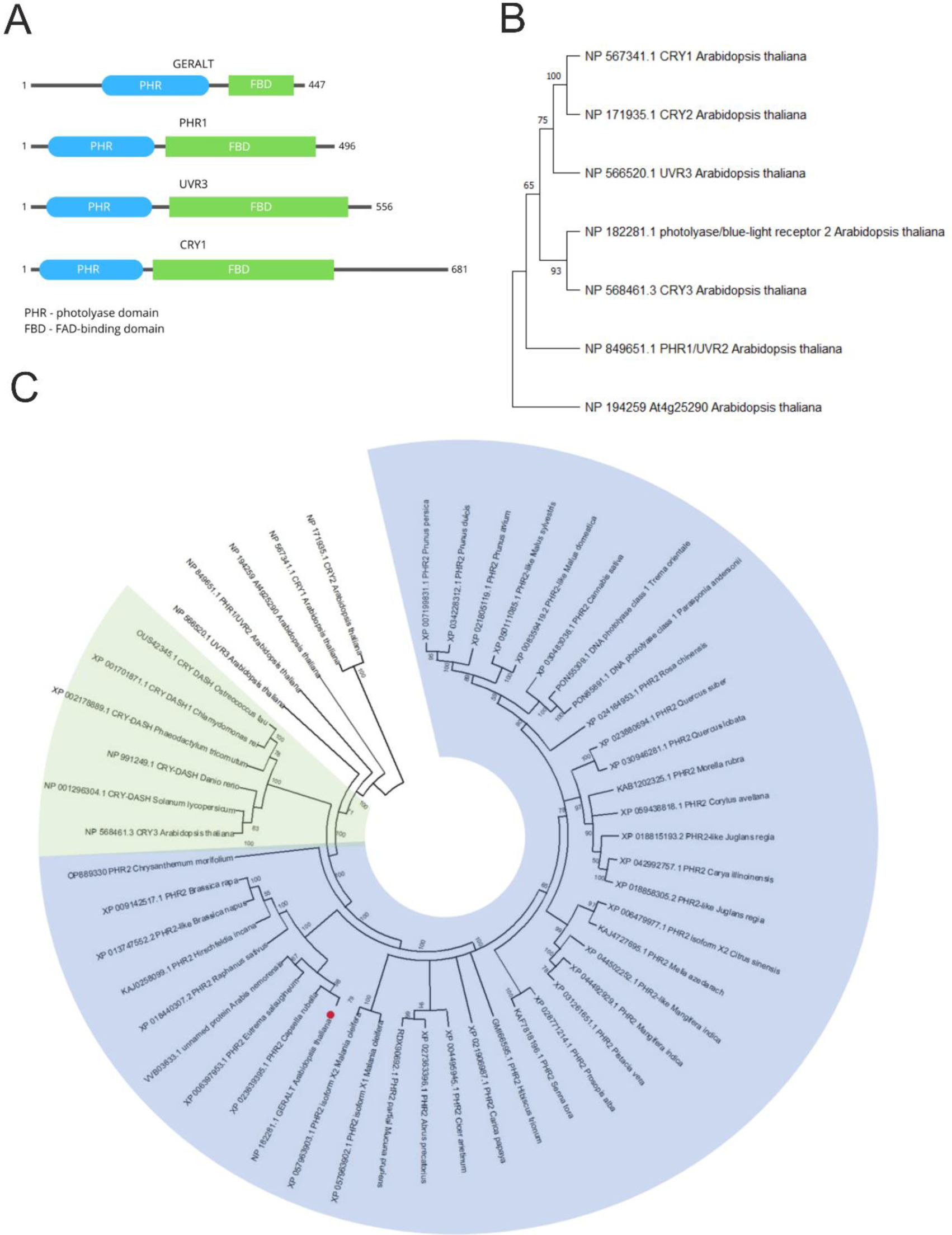
GERALT domain structure and position of this protein in cryptochrome-photolyase phylogenetic tree. (A) The domain structure of GERALT protein. Based on InterPro 97.0 database two main domains, i.e. N-terminal Crypto_Photolyase_N_sf (IPR036155) and C-terminal Crypto/Photolyase_FAD-like (IPR036134) are predicted. (B) Phylogenetic tree of Arabidopsis CPF proteins. (C) Phylogenetic tree of selected CPF proteins. The sequence of all proteins used to construct the tree is given in Supplementary Data S1. CRY-DASH proteins are on a green background, PL/PHR2 subclade is on a blue background. GERALT protein is marked with a red dot.

The homozygosity of *geralt* (SALK_116579) plants was verified by PCR (Supplemental Fig. S1B). The full lenght *GERALT* transcript is not found in *geralt* plants indicating that these plants are knock-out mutants (Supplemental Fig. S1C).

### 2.2. Phylogenetic analysis

Phylogenetic analysis reveals that GERALT, together with other plant PPL/PHR2 proteins, form a separate subclade of CPF proteins (Fig. 1C). This subclade and CRY-DASH are sister groups. All other Arabidopsis CPF proteins (if both cryptochromes are considered together) belong into separate branches (Fig. 1B).

### 2.3. Phenotype of the mutant and transgenic lines

The time of the main root emergence (2 days after seed sowing) as well as the frequency of seed germination (nearly 100%) were similar in *geralt* and WT but seedlings of *geralt* can survive only *in vitro* due to their need of sugar supplementation. When grown with 1% sucrose *geralt* develops a main root, hypocotyl and cotyledons, but all seedlings are smaller and show much greater variability in size, when compared to WT (Fig. 2A-D). Their albino cotyledons are mostly fully expanded (Fig. 2B), but in opposite to WT most of the cotyledons as well as the upper part of hypocotyl are pinkish due to the elevated anthocyanin level (Fig. 2B, C). Few *geralt* seedlings (16%) have small undeveloped cotyledons which often remain coated by the remnants of the seed coat (Fig. 2D). After 12 days in culture, WT seedlings have 2-3 pairs of leaves and well-developed root system with 5-8 lateral roots (Fig. 2E). In contrast, mutant seedlings exhibit great variability in size but are always smaller than the WT (Fig. 2F-I). The root system of *geralt* is poorly developed with 1-3 short and slender lateral roots (ca. 80% of seedlings), or lacks lateral roots altogether (ca. 20% of seedlings). The cotyledons are either light green (ca. 75% of seedlings) or completely white (ca. 25% of seedlings), both with signs of vitrification. Some mutant seedlings (22%), even those with albinotic cotyledons, produce 1-2 pairs of green leaves (Fig. 2G).

**Fig. 2.**
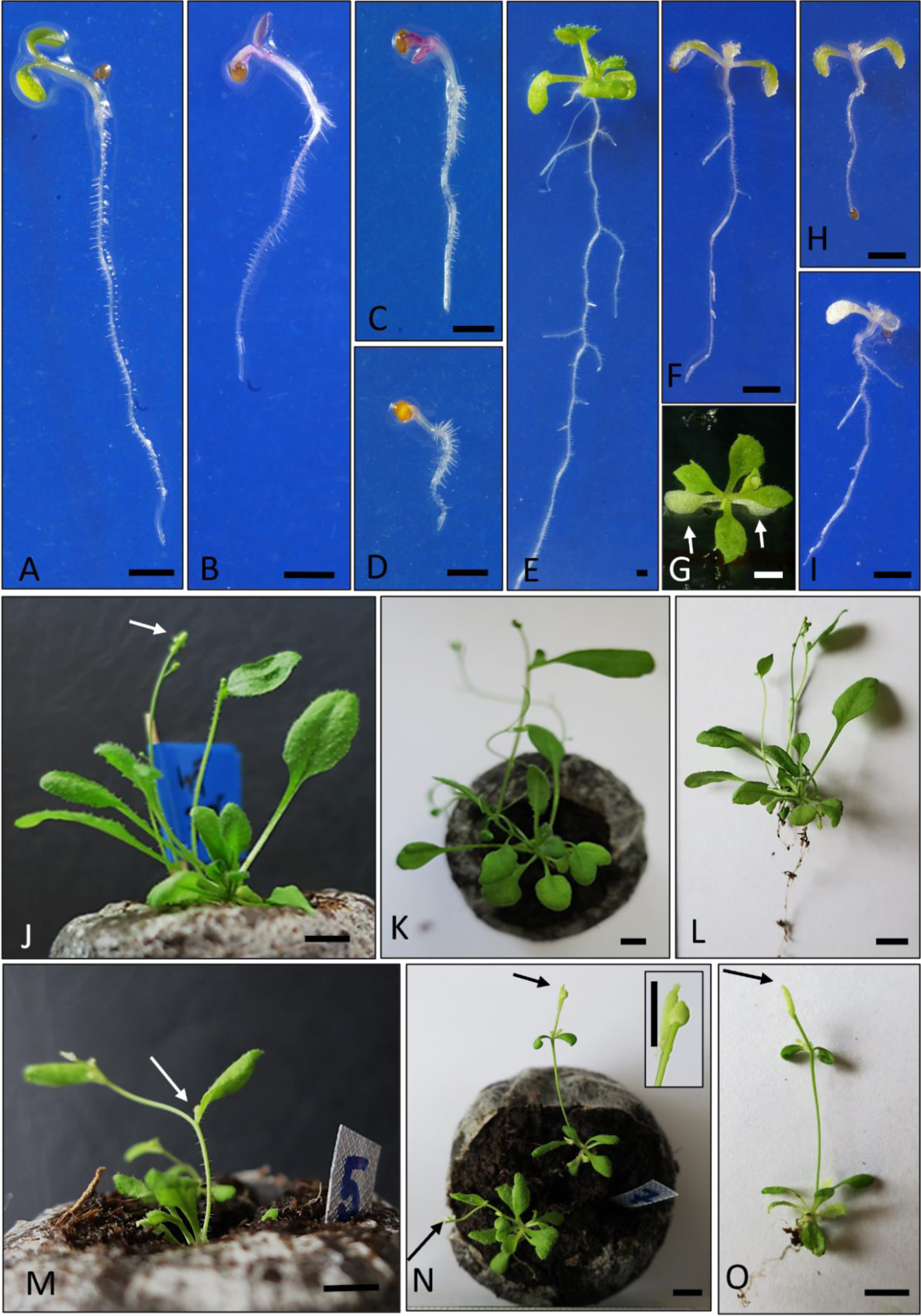
Phenotypes of geralt and WT seedlings and plants. Images of seedlings at 5 (A-D) and 12 (E-I) day and plants at 42 day after sowing (J-O). (A, E, J-L) WT plants. (B-D, F-I and M-O) geralt plants. Arrows indicate albino cotyledons (G) and inflorescence or silique (J, M, N, O). The inset in (N) magnifies the top of the shoot for a more detailed view. Bar represents: 1 mm (A-I) and 0.5 mm (J-O).

Healthy, green *geralt* seedlings can survive after transfer to soil. At 42 days after sowing, all mutants have a dwarf phenotype and are smaller than WT plants (Fig. 2J-O). Most of them produce only a small rosette (ca 85% of plants), but some (ca. 15%) have a single inflorescence stem of different length with 1-2 leaves. The leaves of the rosette and those growing on the stem are wrinkled and serrated when compared to WT ones (Fig. 2K, N). At this stage of development, all mutants with inflorescence stem produce small inflorescences; single siliques can also be observed (Fig. 2N inset). The retarded growth of the mutants is observed, even 5-6 month old plants are still alive and produce new inflorescences and seeds (Supplemental Fig. S2C). The seedlings of the second T-DNA line tested (SALK_149834) with the insertion in the *GERALT* gene have similar phenotype to *geralt* mutant (compare Supplemental Fig. S2A and B).

The full length coding sequence (CDS) of *GERALT* transcript is not detected in the mutant, thus one may link the observed phenotype with the lack of GERALT protein. To confirm this mutant plants were complemented with the full length CDS of the *GERALT* gene under 35S promoter. These lines are marked as *geralt*_OE. On the 5^th^ day, all complemented lines of *geralt* exhibited a WT phenotype (compare Fig. 2A with Supplemental Fig. S3A-D). Over the next 7 days, seedlings of lines *geralt*-OE_2F and *geralt*-OE_9A developed similarly to WT ones, having 2-3 pairs of green leaves and a well-developed root system (Fig. 2E, Supplemental Fig. S3E, G). In contrast, seedlings of lines *geralt*-OE_3C and WT-OE_3E (i.e. WT plants transformed with *geralt* CDS under 35S promoter) elongated and only those of 3E line sporadically (8%) produced single, slender, and short lateral roots (Supplemental Fig. S3F, H). Some seedlings from lines *geralt*-OE_3C (12%) and *geralt*-OE_3E (18%) only produced single leaves (Fig. S3F, H), while the rest failed to produce any leaves at all. Since the greatest phenotypic differences between WT and mutant plants are observed at the early stages of the plant life we focused mainly on early seedling development from 5 to 12 days.

### 2.4. Chloroplast ultrastructure

The albino cotyledons and pale green leaves point to the role of GERALT protein in regulation of chloroplast functioning. The chloroplast ultrastructure of seedlings and adult WT and *geralt* mutants were compared using transmission electron microscopy technique (TEM). Chloroplasts of 8 day-old WT seedlings are lens-shaped with the well-developed thylakoids and grana stacks (Fig. 3A). In 8 day-old seedlings of *geralt* mutants, two types of chloroplasts are observed: (i) having a centrally located network intertwining tubules surrounded by stroma, and few large vacuoles with fibrillar materials and electron-dense plastoglobules mostly arranged in groups (Fig. 3B), (ii) having large grana stacks and very weakly developed stromal thylakoids, grana stacks are usually swollen (Fig. 3C). Chloroplast of adult WT plants are disk-shaped, with well-developed thylakoids and grana stacks (Fig. 3D). In contrast, *geralt* chloroplasts are from nearly spherical to elongated in shape with larger and thicker grana stacks arranged irregularly in relation to the long axis of the chloroplasts (Fig. 3E).

**Fig. 3.**
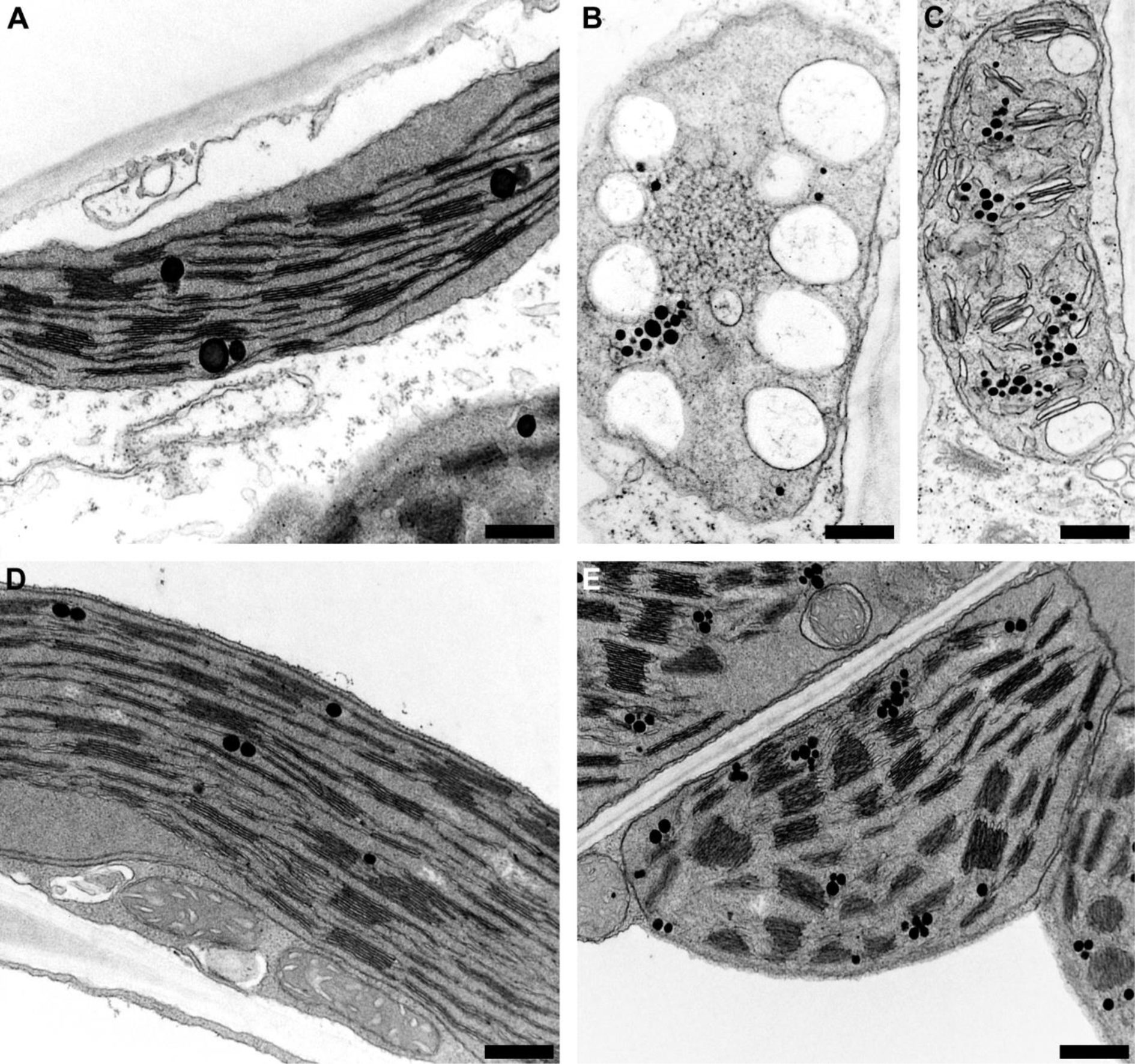
Ultrastructure of geralt chloroplasts. TEM images of chloroplasts from: i) 8 day-old seedlings grown in vitro (A, B, C) and ii) soil grown adult plants (D, E). (A, D) WT, (B, C, E) geralt plants. Scale bars: 0.4 µm (A-D) and 0.5 µm (E).

Confocal microscopy was used to study in depth the chloroplasts of the mutant seedlings. Knockout of *GERALT* gene results in significant reduction of chloroplast size (Fig. 4). The difference is especially obvious in mesophyll cells; in WT plants these cells are almost completely filled with chloroplasts while in mutant a lot of chloroplast-free space is visible. For overwiev, see CLSM images (Supplemental Fig. S4) and CLSM Z-stacks of WT and *geralt* cotyledons (Supplemental Video S1 and Supplemental Video S2). In addition, 3D overviews of individual WT and *geralt* chloroplasts are presented as Z-stack movies (Supplementary Video S3 and Supplementary Video S4). If analyzed quantitatively, the length of major axis of mutant both mesophyll and pavement chloroplasts is about half as long as of WT ones (Fig. 4E). Overexpression of GERALT-GFP under 35S promotor does not resulted in biologically significant difference of chloroplast size as compared with WT plants (Supplemental Fig. S5). Complementation of the mutant with *GERALT* CDS restored normal, WT-like chloroplast size (Supplemental Fig. S5). It is worth noting, that the difference in chloroplast size between WT and *geralt* mutant persists over plant growth. In 5 day-old WT seedlings chloroplasts are in overall smaller, than in 12 day-old ones. The same applies to WT overexpressor and *geralt* complemented lines. Chloroplasts of *geralt* mutant also slightly grow between 5^th^ and 12^th^ day after sowing, however do not even reach the size found in 5 day-old WT plants (Supplemental Fig. S5). The reduced chloroplast size seems to be associated with enormously high granal stacks and weakly developed stromal thylakoids (Fig. 4F) in mesophyll chloroplasts, and almost of the same size in pavement chloroplasts of the mutant (Fig. 4G). Observations of higher grana thickness of *geralt* chloroplasts are in agreement with TEM results (compare Fig. 4F and G with Fig. 3E).

**Fig. 4.**
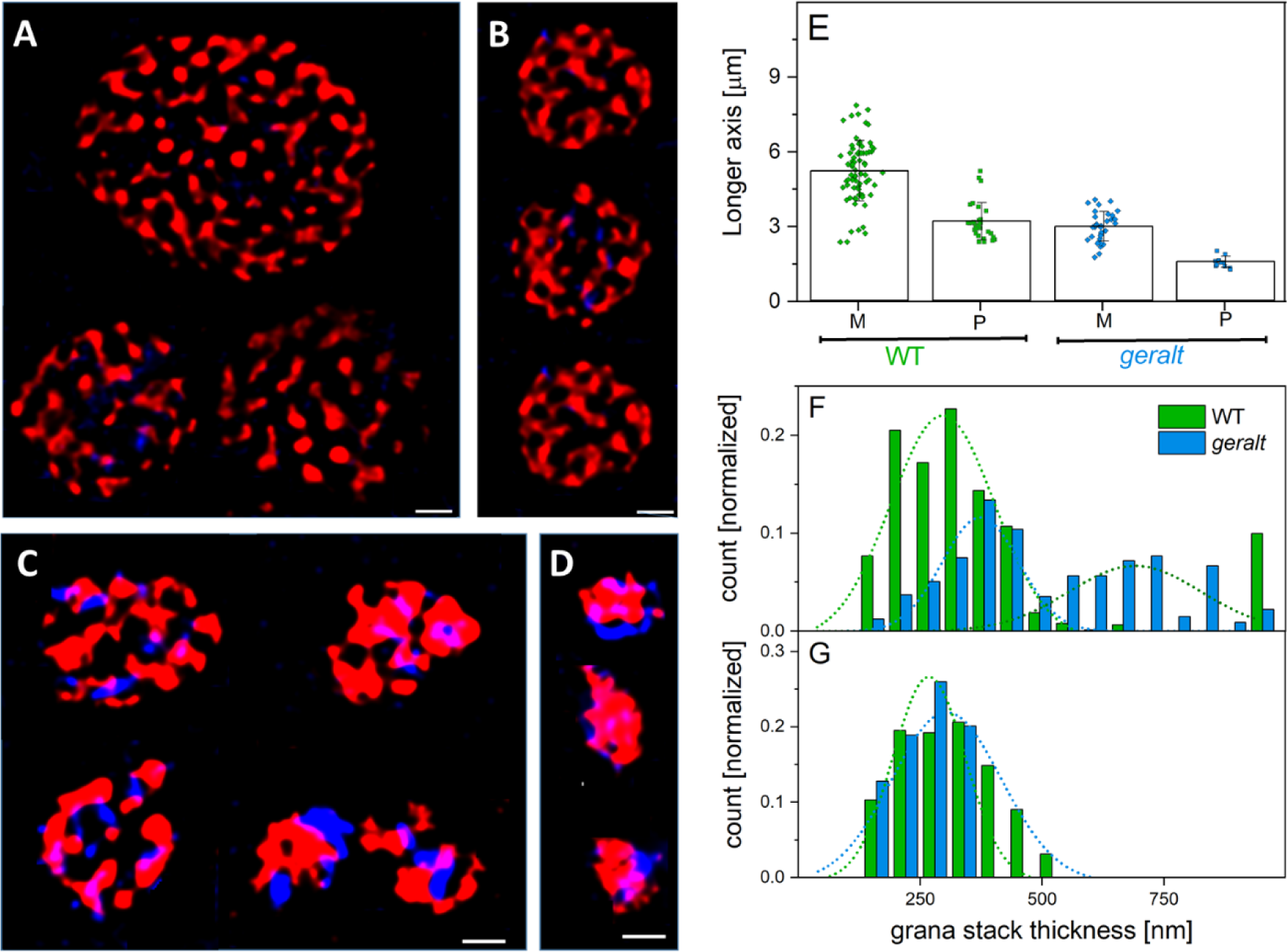
SIM2 images of geralt and WT chloroplasts. Representative SIM2 images of WT (A, B) and geralt (C, D) chloroplasts from mesophyll (A, C) and pavement (B, D) cells from cotyledons of 5 day-old seedlings. Images (which are cropped from a bigger picture) show an overlay of chlorophyll (red) and DAPI (blue) emission. Scale bar = 1 μm; note about 1.5 – fold magnification of images of geralt chloroplasts (i.e. A, B vs C, D). (E) The length of the major axis of mesophyll - M and pavement - P chloroplasts from WT and geralt cotyledons. Points represent individual data, while bars show mean and SD. Comparison of grana stack thickness of mesophyll (F) and pavement (G) WT and geralt chloroplasts. Z-stack images for WT and geralt mesophyll cell chloroplasts are available as Supplementary Video S1 and S2.

### 2.5. Levels of photosynthetic pigments

To characterize further chloroplast functioning in *geralt* plants the levels of photosynthetic pigments were analyzed (Fig. 5). As expected, the level of total chlorophyll (Chl) is much lower (about 25 times) in *geralt* mutant than in WT plants (Fig. 5A). What is more, the content of Chl is comparable in 5 and 12 day-old WT seedlings, also in the case of the mutant, this parameter hardly changed at both analyzed time points. Interestingly, such a big difference is not found in chlorophyllide (Chlide) amount; at 12 day-old seedlings there is only 5 times less total protochlorophyllide (Pchlide) in *geralt* than in WT. Between 5^th^ and 12^th^ day of growth an accumulation trend (about 2 times increase) in Chlide amount in WT but not in *geralt* plants is observed. It should be noted that no Chlide b is detected in *geralt* seedlings, probably its amount is below the sensitivity level of the method used. Interestingly, the Chl a/b ratio is about 2.1 for 5 day-old WT plants and drops to 1.6 at the 12^th^ day of culture. In 5 day-old mutant seedlings this ratio is lower (1.2) and drops to about 1 in 12 day-old plants.

**Fig. 5.**
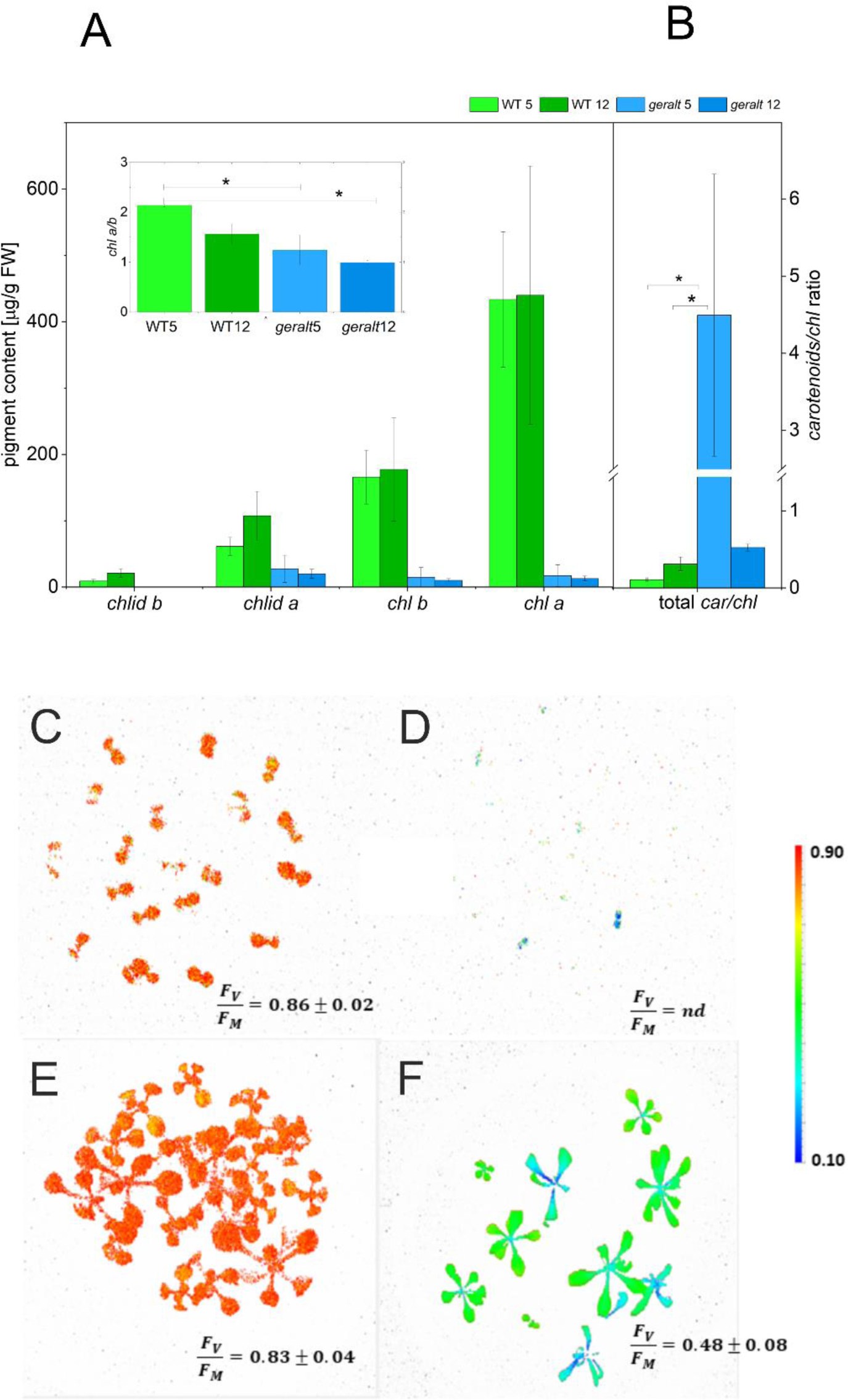
Composition of photosynthetic pigments and photosynthetic activity in 5 day-old and 12 day-old seedlings of WT and *geralt* mutant. (A) Chlorophyllide (chlide) and chlorophyll (chl) content (mg/g fresh weight) and (B) total carotenoid (car) per chlorophyll ratio in 5 and 12 day-old seedlings. Values are means ± SE calculated from at least three independent biological repetitions. Asterisks indicate statistically significant differences between samples (*p-value < 0.01, calculated with one way Anova with a post-hoc TukeyHSD). The maximum PSII quantum yield (F_V_/F_M_) of (C,F) WT and (D,F) *geralt* seedlings measured for (C, D) 5-day-old and (E, F) 12-day-old seedlings, that were pre-adapted to darkness for 10 min prior to the fluorescence measurements. Mean F_V_/F_M_ values with standard deviations are respectively shown in the figures (calculated for 3 independent biological repetitions).

Total content of carotenoids, similarly as content of Chl, does not change significantly between 5 day-old and 12 day-old WT plants (Fig. 5B). The same is true for *geralt* mutant, although there is about 5 time more carotenoids in WT plants. This results in a huge variation in carotenoids/Chl ratio between WT and mutant plants. In WT this ratio is about 0.14 at both time points, while in 5 day-old *geralt* seedlings it reaches almost 4.5 and drop to about 0.6 at day 12. This indicates an excess of carotenoids in relation to possible sites of their binding in photosynthetic pigment-protein complexes in *geralt* chloroplasts.

### 2.6. Photosynthetic efficiency

To characterize the functioning of the photosynthetic apparatus, the maximal PSII quantum yield (F_V_/F_M_) of the mutant was measured. For 12 day-old *geralt* seedlings F_V_/F_M_ stays at much lower level than for respective WT seedlings (Fig. 5C to F), and it could even not be measured for 5-day-old *geralt* seedlings (Fig. 5D). This confirms the functional impairment of PSII in *geralt* seedlings. Interestingly, F_V_/F_M_ in 5-day-old WT (Fig. 5C) and *geralt*-OE_3C and 9A (Supplementary Fig. S6) exceeded 0.85 which is typical for healthy plants (Bjőrkman and Demmig, 1987).

### 2.7. Gene expression

To determine how the *geralt* mutation affects the global transcriptome and whether it reflects the changes in phenotype with age the Next-Generation Sequencing (NGS) was used. Whereas the steady-state levels of 416 of 27,655 genes (p-value Bonferroni correction ≤ 0.05) differ by more than 2-fold (decrease or increase) in 5 day-old (*geralt* vs WT) seedlings, only 125 such transcripts are found in 12 day-old seedlings (Supplemental Excel File S1, Fig. 6A). Only 17 genes are differentially expressed in both plant age groups studied (Supplemental Table S1). The gene which expression is the most severly inhibited in the mutant (920 and 461 times lower at 5 day-old and 12 day-old seedlings, respectively) is *At4g13575* encoding protein with an unknown function.

**Fig. 6.**
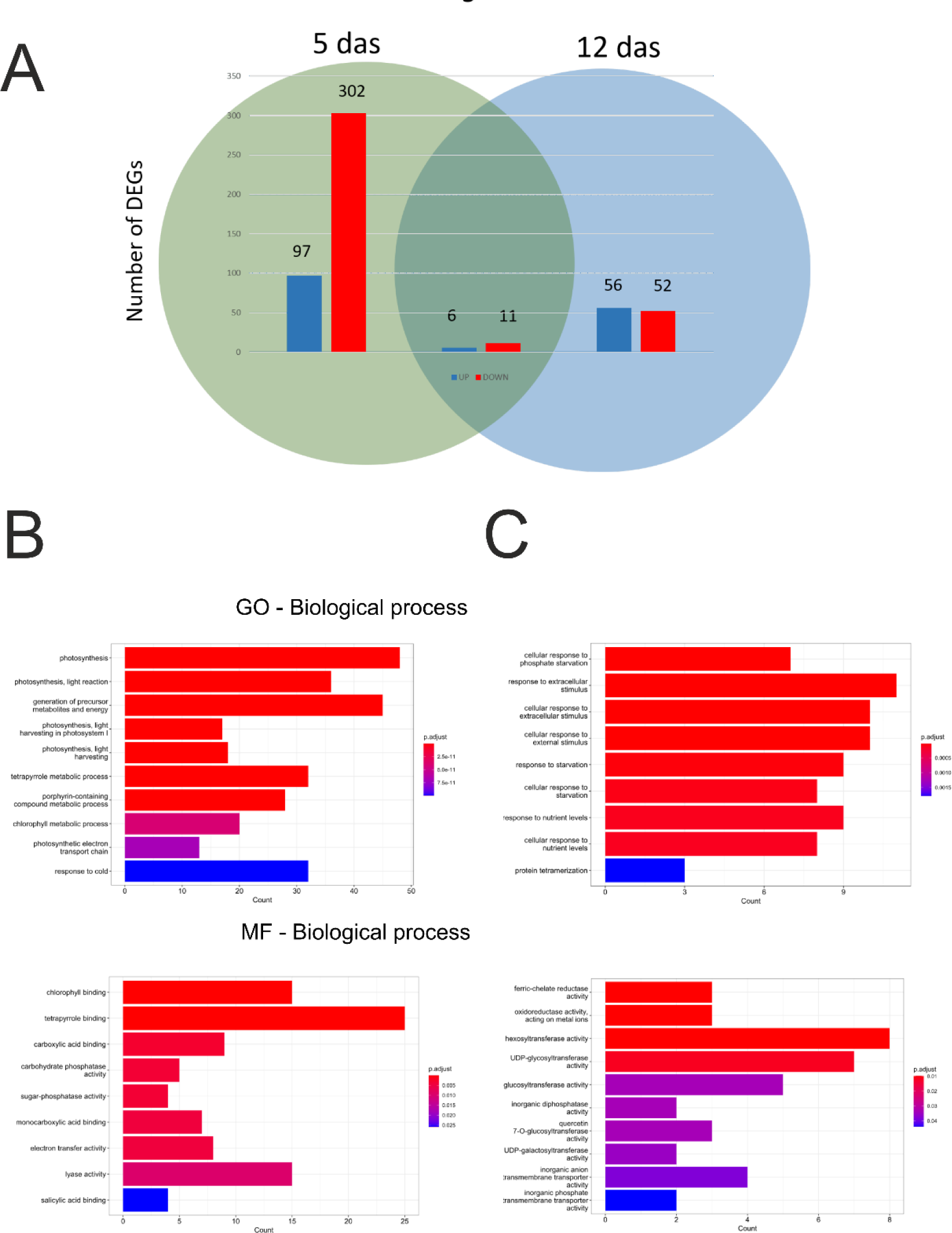
Differentially expressed genes (DEGs) (p-value Bonferroni correction ≤ 0.05) which differ over 2-fold between *geralt* and WT seedlings. (A) Venn diagram of DEGs between geralt and WT in 5-day old and 12 day-old seedlings. Numbers of up- and down regulated genes are shown as blue and red bars respectively. Gene ontology represented as molecular function and biological processes in (B) 5 day-old and (C) 12 day-old seedlings. (B) GO Biological Process, ten genes categories, Molecular function (MF), the list of nine activities present in proteins encoded by genes with significantly enrichment in GO molecular function categories out of 417 genes (Bonferroni corrected p-value). (C) GO Biological Process, ten genes categories, Molecular function (MF), the list of nine activities present in proteins encoded by genes with significantly enrichment in GO molecular function categories out of 125 genes (Bonferroni corrected p-value).

As described above, with the age of the plants, the differences in phenotypes, in the levels of photosynthetic pigments and PSII photosynthetic efficiencies between mutants and WT decrease. They are accompanied by a decrease in the number of DEGs related to photosynthesis. (Fig. 6B vs Fig. 6C – molecular function, Supplemental Excel File S1, Supplemental Table S2). Among such genes are also these encoded in chloroplasts with a characteristic drop in PEP-dependent and increase in NEP-dependent transcript levels.

To make sure that *geralt* mutation indeed influences expression of chloroplast genes real-time PCR analysis was performed. In this experiment 7 day-old seedlings dark-adapted overnight and illuminated for 3 h with 50 µmol·m^-2^·s^-1^ of blue or red light or left in the darkness were used. Different light conditions were used to investigate whether GERALT may act as a blue light photoreceptor to regulate gene expression. In this case, red light with the same intensity as blue light served as a control for the blue-light specific response. The amounts of PEP-dependent chloroplast transcripts (*psbD, psbB, psaB, rbcL*) tested with real-time PCR are lower in the mutant (Fig. 7A). At the same time the steady state levels of NEP-dependent transcripts (*rpoA, accD, rpoC1*) are higher than in WT plants (Fig. 7B). The levels of transcript encoded by both NEP and PEP are similar (*atpB*) or higher (*clpP*) in *geralt* than in WT under all light condition used (Fig. 7C). None of the chloroplast gene tested are regulated by the applied light conditions.

**Fig. 7.**
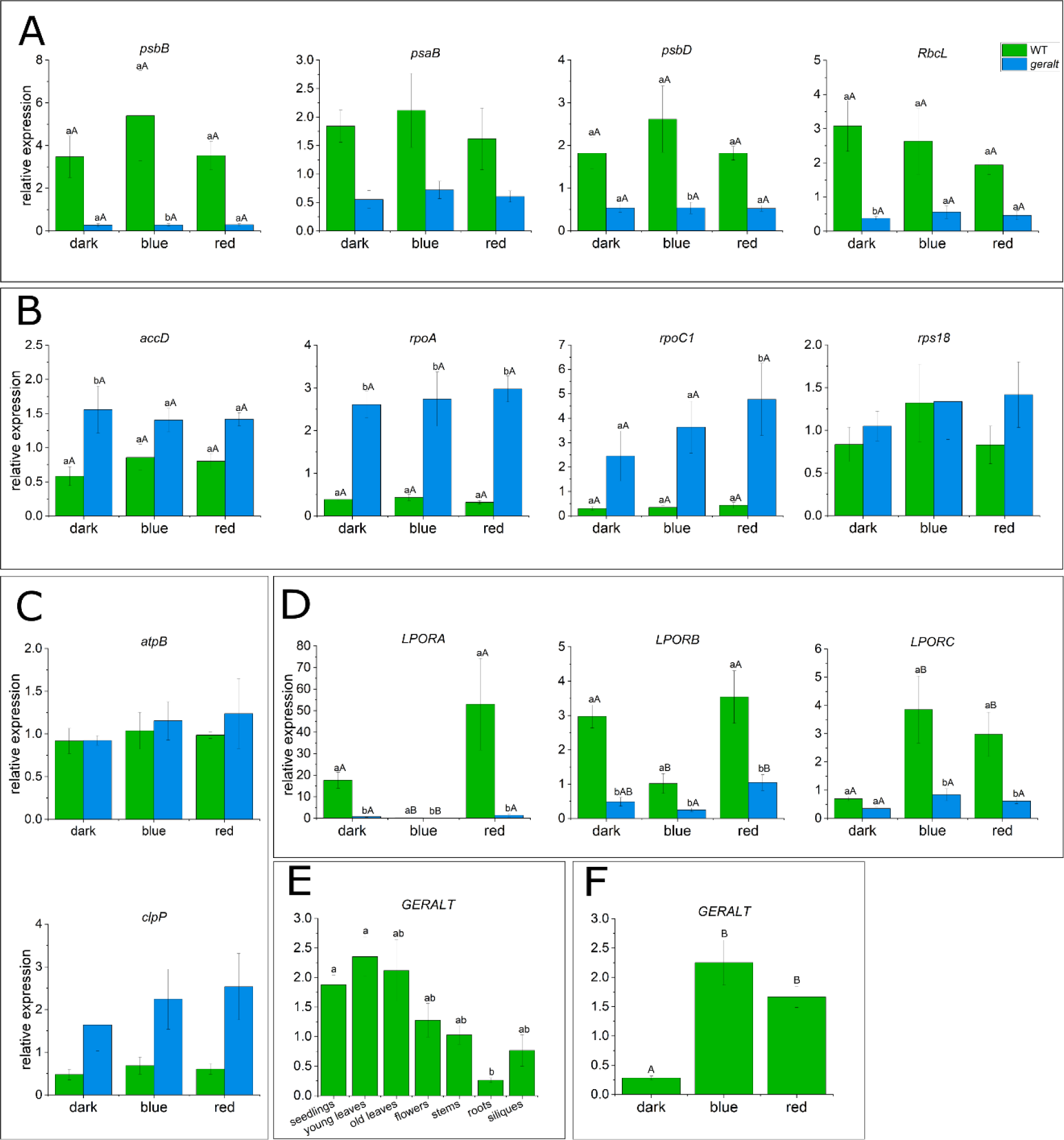
The expression of GERALT in WT plants and the expression of chloroplast and nuclear genes in *geralt* mutant. The expression of chloroplast (A) PEP-dependent, (B) NEP-dependent, (C) PEP+NEP-dependent genes and (D) LPORA, LPORB and LPORC genes in 7 day-old in vitro grown WT and geralt seedlings. 6 day-old in vitro grown WT seedlings were dark adapted overnight and, at the next day, they were illuminated for 3 h with 50 µmol m−2s−1 of blue (B - 470 nm) or red (R - 660 nm) light or stayed in dark. The illumination started about approximately 2 h after turn on the light in the culture chamber. (E) The steady-state levels of GERALT mRNA in 7 day-old in vitro grown WT seedlings and different organs of adult soil grown plants. The leaves with visible yellowing symptoms are described as old leaves. The seedlings and plant organs were harvested approximately 4 h after turn on the light in the culture chamber. (F) The impact of 3h illumination with 50 µmol m−2s−1 of blue (B - 470 nm) or red (R-660 nm) light on GERALT mRNA levels. 6 day-old in vitro grown WT seedlings were dark adapted overnight and, at the next day, they were illuminated with light or stayed in dark. Each bar corresponds to an average of at least 3 biological replicates. Error bars represent SE. Different lowercase letters (a and b) indicate significant differences (p-value ≤ 0.05) between (A-D,F) WT and geralt for a given illumination conditions or (E) different organs, different capital letters (A and B) indicate significant differences (p-value ≤ 0.05) between light conditions for a given genotype.

Because of perturbed greening in the mutant the steady-state levels of protochlorophyllide oxidoreductase (*LPOR*) genes were examined. The levels of *LPORA* and *LPORB* are lower in *geralt* seedlings under all applied light conditions with a pattern of *LPORA* light-dependent regulation similar to that observed in WT plants (Fig. 7D). While the blue-light visibly reduces *LPORB* transcript levels in WT plants it is almost uneffective in *geralt* mutant. The dark levels of *LPORC* mRNA in WT and *geralt* are similar with the lack of light up-regulation of this gene in the mutant (Fig. 7D).

### 2.8. GO analysis

Genes selected based on significant changes in expression levels (Bonferroni-corrected p-value) between *geralt* mutant and WT plants at two time points were analyzed. Gene Ontology analysis (GO Biological Process) of 417 genes identified in 5-day-old plants (Bonferroni-corrected p-value) showed that most of the significantly different genes belong to categories such as photosynthesis, mainly light reactions, and production of precursor metabolites and energy (Figure 6B, upper graph). The functions the encoded proteins perform (GO: Molecular Functions) include binding tetrapyrroles and Chls as well as participating in sugar metabolism (Fig. 6B, lower graph). GO analysis for 4,702 genes whose expression was significantly different between the mutant and WT seedlings (based on p-value) is shown in Supplementary Fig. S7A and C.

Analysis (GO Biological Process) of 125 differentially expressed genes (Bonferroni-corrected p-value) identified in 12-day-old *geralt* seedlings showed that most of them are involved in the response to extracellular stimuli and phosphate starvation, or the overall response to nutrient levels (Figure 6C upper graph). These genes encode primarily proteins with transferase activity (hexosyltransferase, UDP-glycosyltransferase, glucosyltransferase) and transmembrane transporter activity (Fig. 6C bottom graph). GO analysis for 2,663 genes whose expression changed significantly (based on p-value) is shown in Supplementary Fig. S7B and D.

### 2.9. Expression of the GERALT gene

The described above differences between WT and *geralt* seedlings suggest the role of the tested gene in regulation of chloroplast functioning. Indeed, the mRNA of the *GERALT* is present mainly in green parts of plants with the lowest level of this transcript in roots (Fig. 7E). The steady-state levels of *GERALT* mRNA increase clearly under blue and red light (Fig. 7F).

### 2.10. Subcellular and suborganellar localization of GERALT protein

The *in silico* analysis using several bioinformatic tools collected in the SUBA5 database (https://suba.live/; (Hooper et al., 2017) predicts the localization of GERALT in various compartments, including plastids, mitochondria, cytosol, nuclei. None of the software tested (TargetP-2.0 https://services.healthtech.dtu.dk/services/TargetP-2.0/; LOCALIZER https://localizer.csiro.au/; PredSL http://aias.biol.uoa.gr/PredSL/index.html; TPPred3.0 https://tppred3.biocomp.unibo.it/ welcome/default/index) predicts the presense of the transit peptide responsible for transporting this protein into plastids. To conclusively resolve the subcellular localization of the tested protein transgenic Arabidopsis plants (WT-OE_3E) expressing GERALT-GFP fusion protein under the control of 35S promoter were used.

Green fluorescence, clearly identified as GFP by λ-stack based emission spectra with GFP-specific emission maximum ∼ 510 nm, in comparison to a weak green chloroplast autofluorescence with emission maximum ∼ 550 nm, is found exclusively in WT-OE_3E chloroplasts (Supplemental Fig. S8). The pavement cell chloroplasts of this plants are characterized by the highest intensity of GFP signal, especially in a meristematic-like tissue of young (3 day-old) cotyledons (Supplemental Fig. S8 E-H). The GERALT-GFP signal is also identified in mesophyll chloroplasts (Supplemental Fig. S8 I-L), however it was less pronounced, we found it in some (but not all) tested leaf fragments and not in all plants tested. Interestingly, GERALT-GFP protein diffuses freely to stromulae formed between pavement cell chloroplasts and nuclei (Supplemental Fig. S8 M-R).

Detailed characterization of localization of GERALT-GFP within chloroplast was possible using SIM^2^ technique (Fig. 8A). Clear GFP signal is found within chloroplast interior (stroma) without colocalising with Chl emission. There was also no clear colocalisation of GFP and DAPI emission, suggesting no direct interaction of GERALT-GFP with chloroplast nucleoids (at least under the conditions tested). A 3D overview of described chloroplasts can bee seen in Z-stack based movie available in Supplementary materials (Supplemental Movie S5). It is noticeable at this video that in Z-axis GFP occupies space extending over the chloroplast edges, determined by Chl fluorescence. This is most likely due to the large volume of stroma characteristic of pavement chloroplasts.

**Fig. 8.**
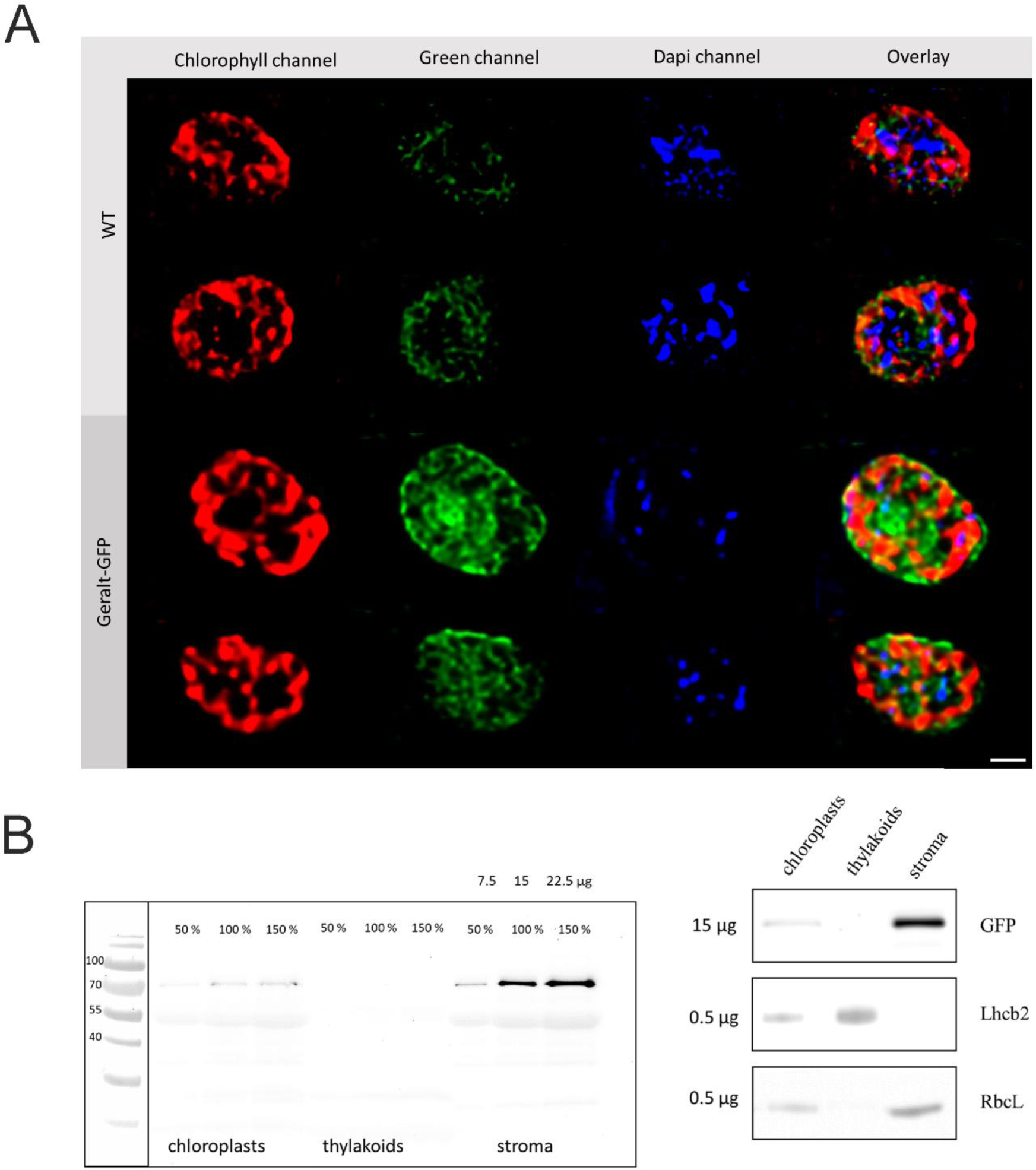
GERALT-GFP fusion protein is mainly localized in chloroplast stroma. (A) SIM2 images of pavement chloroplast of WT and WT-OE_3E overexpressing GERALT-GFP fusion protein. Chlorophyll channel - excitation 470 nm, emission 650-750 nm. Green channel - excitation 470 nm, emission 490-530 nm, DAPI channel - excitation 405 nm, emission 440-480 nm. Overlay – superimposition of all three channels. Images are cropped from bigger pictures. Scale bar = 1 μm. (B) Western blot analysis of GERALT-GFP subchloroplast localization. Proteins isolated from chloroplasts purified from WT-OE3E 5 week old soil grown plants were separated into stroma and thylakoid fractions, separated by SDS PAGE (12 % polyacrylamide gels) and electroblotted. Proteins were probed with anti-GFP, anti-RbcL or anti-Lhcb2 antibodies.

It is worth to remind here, that in SIM^2^ technique some artifacts might be produced, especially due to light scattering at the edges of organelles (Iwai et al., 2018). Therefore, the apperance of a green envelope in the images correlating with chloroplast edges should be interpreted with caution. It is not probably related to signal from GERALT-GFP fusion protein. The above mentioned light-scattering artifact and autofluorescence originating from flavin emission, might be also responsible for weak green fluorescence found in WT chloroplasts.

The localization of GERALT protein in the stromal fraction as predicted using Plastogram software ((http://biongram.biotech.uni.wroc.pl/PlastoGram/, (Sidorczuk et al., 2023)) and observed using confocal microscopy (Fig. 8A) was confirmed with the Western-blot technique (Fig. 8B).

## 3. Discussion

### 3.1. GERALT belongs to PPL/PHR2 subclade of CPF proteins

Phylogenetic analysis reveals that GERALT belongs to PPL/PHR2 subfamily of CPF proteins. (Fig. 1C, (Mei and Dvornyk, 2015; Deppisch et al., 2022)). Two unique: ppl2 and ppl3 motifs which separate PPLs/PHR2s from other subclades of CPF only have been identified (Deppisch et al., 2022). The analysis of all Arabidopsis genes encoding CPF proteins also shows the close relationship between GERALT and AtCRY3 belonging to PPL/PHR2 and CRY-DASH subclade, respectively (Fig. 1B). CRY-DASH proteins have appeared early during evolution, as a protein belonging to this subgroup has been found in at least one Archeon genus. It is proposed that PPL/PHR2 subclade apeared as a result of CRY-DASH gene duplication (Deppisch et al., 2022). As PPL/PHR2s are specific to *Viridiplantae* and *Rhodophyta* (red algae), this duplication probably occured in archaeplastid common ancestor of these groups (Mei and Dvornyk, 2015; Brawley et al., 2017). From then PPL/PHR2 and CRY-DASH proteins evolved independently. The analysis of amino acid sequences of GERALT and CRY-DASH proteins from Arabidopsis, *Chlamydomonas, Solanum, Danio* and *Osterococcus* indicates that the similarity between CRY-DASH proteins from evolutionarily distant organisms is higher than between GERALT and AtCRY3 (Supplemental Fig. S9).

### 3.2. Chloroplast biogenesis and chlorophyll biosynthesis are disturbed in geralt mutants

In this work we documented that GERALT protein is localized in chloroplasts (Fig. 8). We observed this protein in stroma and in stromules, i.e. stroma-filled tubular plastid extensions (Supplemental Fig. S8).

The *GERALT* mRNA is expressed mainly in organs containing chloroplasts (Fig. 7E). According to PlantCare (https://bioinformatics.psb.ugent.be/webtools/plantcare/ html/; (Lescot et al., 2002) and AGRIS databases (https://agris-knowledgebase.org/AtcisDB/; (Davuluri et al., 2003)) motifs recognized by transcription factors involved in plant response to light are present in the *GERALT* promoter (Supplemental Table S3). Light activation of *GERALT* promoter seems to be one of the factors leading to an increase in steady-state levels of the transcript which expression it controls (Fig. 7F). A logical consequence of this light regulation and localization of GERALT protein in chloroplasts is its involvement in light-dependent modulation of plastid maintenance including their biogenesis, chlorophyll biosynthesis and formation of photosynthetic apparatus. The characteristic features of *geralt* phenotype including albino cotyledons, serrated leaves and retarded growth (Fig. 2, Supplemental Fig. S2) may originate from disruption of these processes (Hricová et al., 2006; Liu et al., 2016; Rodríguez-Alcocer et al., 2023).

The lack of GERALT protein influences chloroplast morphology in Arabidopsis seedlings and mature plants (Fig. 3, Fig. 4). The chloroplast size and shape as well as the thylakoid architecture are disturbed with the more severe effects observed in the seedling cotyledons. The striking features of ultrastructure of *geralt* chloroplast are the high granal stacks and weakly developed stromal thylakoids. This picture is supported by a declined Chl a/b ratio in comparison to WT. This is in agreement with the data of Trotta et al. (Trotta et al., 2019) delivered by use of subsequent differential centrifugation steps, where isolated grana have a lower Chl a/b ratio in comparison to stromal thylakoids and to the total thylakoid membranes. The presence of grana depleted of the stromal thylakoids network negatively affects the functioning of photosynthetic electron transport in *geralt* seedlings, as indicated by a lower PSII efficiency (a detailed study on photosynthetic performance of the mutant is under way). It is important to note here, that during chloroplasts biogenesis the components of photosynthetic electron transport develops in sequence: first, grana are formed with PSII complexes, next, stromal thylakoids with PSI complexes (Liang et al., 2018). The thylakoid architecture in *geralt* mutant suggests that chloroplast biogenesis program stopped before the development of the stromal thylakoids. This is accompanied by a lower expression of genes encoding components of photosynthetic machinery and enzymes responsible for Chl biosynthesis resulting in drastically reduced levels of photosynthetic pigments (Fig. 5A and B, Fig. 6, Supplemental Fig S7, Supplemental Fig. S10A). Consequently, all photosynthetic parameters are lower in *geralt* plants (Fig. 5C and D vs Fig. 5E and F).

### 3.3 Transcriptome is affected in geralt mutant

The analysis of WT and *geralt* seedling transcriptomes with NGS revealed a set of DEGs (Fig. 6, Supplemental Table S1 and S2, Supplemental Excell File S1). Most of them are involved in photosynthesis, transport and metabolism of phosphate, iron and sugars. Beside the genes encoding proteins which different levels can be easily linked with the *geralt* phenotype, there are genes which are not expected to be differentially expressed in the mutant. Among the latter ones, the most common are transcripts encoding proteins involved in protein degradation, response to pathogens (mainly responsible for glucosinolate biosynthesis) and abiotic stresses (Fig. 6B and C, Supplemental Fig. S10). The identified DEGs are encoded both in nuclear and plastid genomes. At this stage, it is difficult to say which differences in transcriptomes depend directly on the lack of the GERALT protein and which are side effects of disturbed photosynthesis and subsequent impaired chloroplast to nucleus retrograde signaling.

The results of the analysis of the functions performed by proteins encoded by DEGs are supported by GO analyses. GO analyses clearly show differences between two time-points investigated: 5 and 12 days of culture. In 5 day-old plants most of the genes affected by the mutation in the *GERALT* gene belong to the categories such as photosynthesis, generation of precursor metabolities and energy and light reactions of photosynthesis categories. In the case of 12 day-old plants the identified genes encode proteins which are involved in response to extracellular stimuli, cellular response to phosphate starvation or general to the response to nutrient levels.

### 3.4 GERALT modulates the levels of PEP- and NEP-transcribed genes

The amounts of PEP-dependent chloroplast transcripts tested with real-time PCR are lower in the mutant, but the differences are not drastic. They may be the effect of the reduced amount of PEP polymerase itself as the slight decline of the nuclear transcripts encoding PEP subunits (i.e. *SIGA* and *PAP8*) are observed in *geralt* seedlings (Fig. 7A, Supplemental Table S2). At the same time the steady state levels of NEP-dependent transcripts are higher than in WT plants (Fig. 7B). It is clearly visible for *rpoA* and *rpoC1* genes encoding PEP subunits. The levels of transcripts encoded by both NEP and PEP are similar or higher in *geralt* than in WT seedlings under all light conditions used (Fig. 7C). Similar tendency, i.e. lower levels of PEP-dependent genes and higher levels of NEP-dependent ones is common for albino and pale green mutants and is typical for plants without functional PEP subunits (Hajdukiewicz and Allison, 1997; Pfalz and Pfannschmidt, 2013; Yoo et al., 2019; Tadini et al., 2020).

### 3.5. The light regulation of LPORB and LPORC is affected in geralt mutant

Three *LPOR* genes are found in Arabidopsis genome and the role of the proteins they encode is intensively studied mainly during seedling de-etiolation. It is well known that illumination of etiolated seedlings cause a rapid decrease of *LPORA* levels and an increase of *LPORC* levels, whereas data concerning *LPORB* are unclear (Gabruk and Mysliwa-Kurdziel, 2015). Interestingly, we demonstrated that the levels of *LPOR* transcripts in 7 day-old Arabidopsis WT seedlings are differentially impacted by blue and red light (Fig. 7D). *LPORA* level is down regulated by blue light and up-regulated by red light in light grown WT seedlings. Only blue light controls *LPORB* mRNA amount causing the decrease in its level. Only for *LPORC* both blue and red light act similarly as positive regulators (Fig. 7D). This is a very interesting result that requires further in-depth research, which, however, is beyond the scope of this study. Nevertheless, focusing on the comparison of the regulation of individual *LPOR* genes in the mutant relative to the WT, we can observe that the light regulation of *LPORB* and *LPORC* mRNA levels is affected in *geralt* seedlings (Fig. 7D). Interestingly, the difference concerns both regulation by of *PORB* and *PORC* levels by blue light and red light-dependent up-regulation of *PORC* observed in WT seedlings. It could indicate that GERALT serves as a blue and red light photoreceptor. Although cryptochromes are primarily blue light photoreceptors, their red light reception mechanism has been described in detail for the *Chlamydomonas* cryptochrome (Oldemeyer et al., 2020).

As mentioned in the Results section, GERALT has a putative FAD binding domain. This domain is shorter as compared with other CPF proteins and lacks some conserved residues (Fig. 1B, (Mei and Dvornyk, 2015; Deppisch et al., 2022)). This suggests that the function of GERALT-associated FAD differs from that described for the chromophore associated with cryptochromes. Therefore, the observed differences in the *LPOR* expression should be interpreted as the result of a lack of modulation of blue/red-light-specific pathways in plants lacking GERALT, rather than as evidence for the direct function of this protein as a photoreceptor.

### 3.6 The role of GERALT is decreasing with the plant age

The biggest differences between WT and *geralt* mutant phenotypes are visible at the cotyledon stage. As the plants grow, these differences slowly diminish, but even adult mutants are clearly different from WT plants. This points to a decreasing role of GERALT as the plant ages. With time the cotyledons slowly turn green, the true leaves of the mutant are greener than cotyledons, the older plants can survive in soil without sugar complementation. Changes in the leaf color easily detected by the naked eye, are accompanied by an increase in the photosynthetic efficiency (F_V_/F_M_) parameter (Fig. 5C andF). While Chl fluorescence is hardly detectable in 5 day-old *geralt* seedlings, F_V_/F_M_ reaches value of 0.5 in 19 day-old plants. Also, the number of DEGs, including genes encoding proteins associated with photosynthesis, decreases (Supplemental Table S2). For example the levels of *HEMA1* (*GLUTAMYL-tRNA REDUCTASE 1*) and *LPOR* genes encoding enzymes catalyzing the first and the penultimate regulatory steps of Chl biosynthesis, respectively, are more affected in younger *geralt* seedlings. Hence, the levels of photosynthetic pigments increase with seedling age, as visible for both Chl a and Chl b (Fig. 5A). Between 5^th^ and 12^th^ day of growth the levels of Chl a and Chl b in *geralt* seedlings rise 4 and 8 times, respectively. A similar effect of seedling age is observed for genes encoding components of PSI and PSII. The most severely down-regulated transcripts are *LHB1B1 (LIGHT-HARVESTING CHLOROPHYLL-PROTEIN COMPLEX II SUBUNIT B1)*, *LHCB2.3 (PHOTOSYSTEM II LIGHT HARVESTING COMPLEX PROTEIN 2.3*)*, RBCS-2B* (*RUBISCO SMALL SUBUNIT 2B*) with respectively 20.0; 33.0; 109 fold lower levels in 5 day-old *geralt* seedlings. The only differentially expressed photosynthesis-related gene undergoing differential expression in 12 day-old *geralt* plants is *RBCS-2B* with the level 21.3 fold lower than in the WT seedlings. Such a pattern suggests that light-dependent photosynthetic machinery is adjusted first during mutant development, while Rubisco (and light-independent machinery of photosynthesis) is still down-regulated in 12 day-old mutant plants.

Cotyledons are formed already during embryogenesis. Proplastids present in all cotyledon cells develop into chloroplasts immediately after illumination. The development of chloroplasts in true leaves is more complex (Charuvi et al., 2012; Pogson et al., 2015). They originate from proplastids coming from the shoot apical meristem and leaf primordia. The occurrence of mutants having green cotyledons and albino/variegated/pale green true leaves (including *var2*) or albino cotyledons but producing green true leaves (i.e. *cyo1/sco2, wco, spd1* and *ecd2*) indicates the existence of two independent developmental programs in these organs (Chen et al., 1999; Yamamoto et al., 2000; Shimada et al., 2007; Ruppel et al., 2011; Wang et al., 2021). The chloroplast ultrastructure, expression of photosynthesis-related genes and photosynthetic capacity is similar in Arabidopsis cotyledons and true leaves pointing that these developmental programs converge at the same endpoint (Shi et al., 2020).

All the features of the mutants described above strongly suggest that the role of GERALT changes during the life of the plant. Our working hypotheses is that GERALT, along with other factors, controls signaling pathways operating both in cotyledons and in true leaves (Fig. 9). In the proposed model GERALT-dependent and GERALT-independent pathways act together and complement each other to ensure the proper plant growth including chloroplast biogenesis, production of pigments and proteins building photosynthetic apparatus. The GERALT-dependent pathways are the most important at the cotyledon stage and their role decreases, though not to zero, as plants age. PEP functioning appears to be one of the processes controlled by both GERALT-dependent and -independent pathways.

**Fig. 9.**
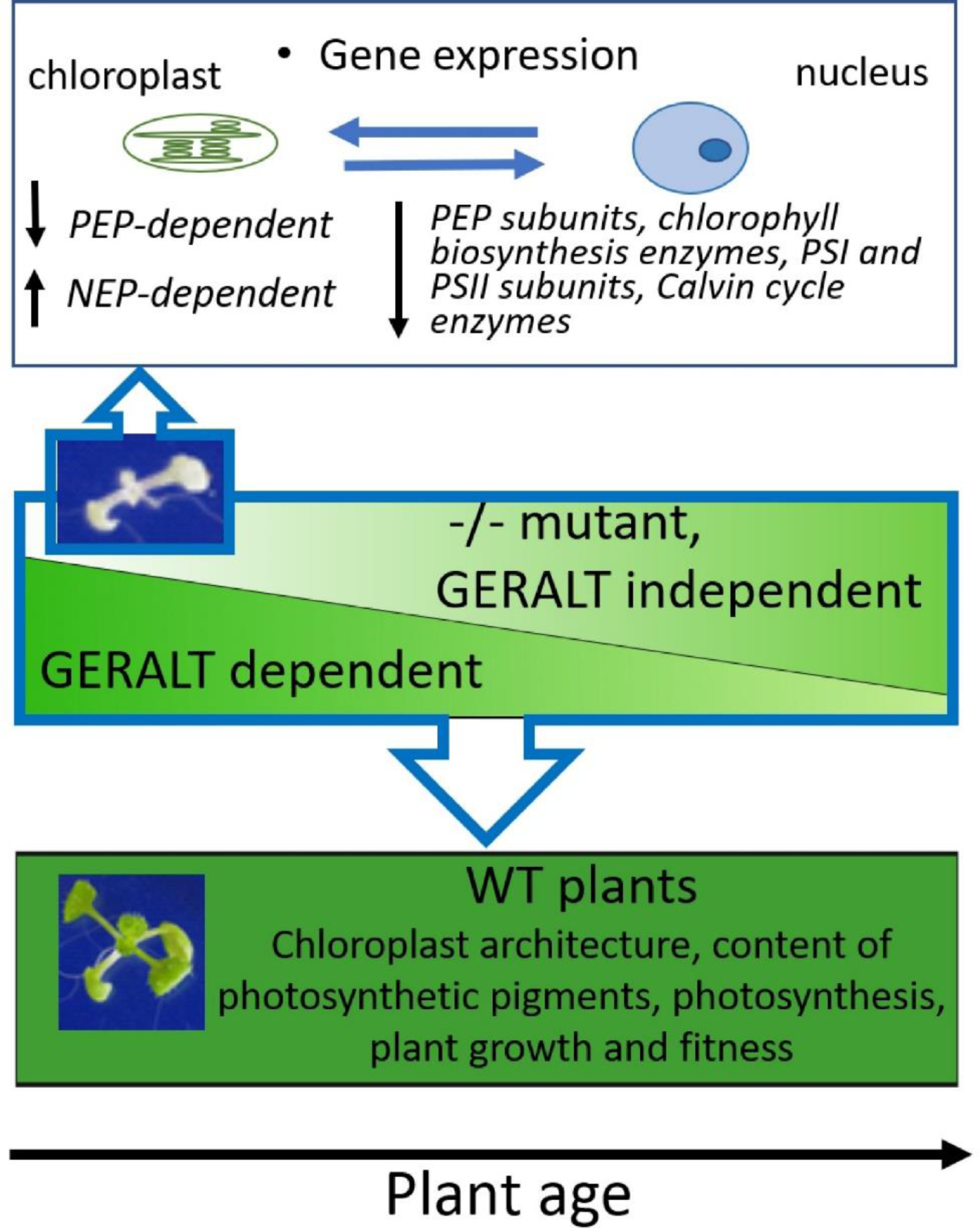
The working model for GERALT role in regulation of plant development. Chloroplast development and chlorophyll biogenesis are controlled by GERALT-dependent and GERALT-independent pathways. Albino cotyledons of mutant seedlings (see the inset) point to the crucial role of GERALT protein at the early stages of plant life. The expression of both nuclear- and chloroplast-encoded photosynthesis associated genes is disturbed in geralt seedlings. Activation of only GERALT-independent pathways results among others in a drop of the levels of PEP-transcribed mRNAs and nuclear transcripts involved in chlorophyll biosynthesis and being components of photosynthetic complexes. These changes lead to disturbance of signaling between chloroplasts and nucleus. As the plant ages, the role of GERALT-dependent pathways diminishes, but their co-operation with those GERALT-independent is essential for maintaining proper plant fitness. The intensity of the green color represents the importance of GERALT-dependent and -independent pathways as they add up to the level highlighted in dark green in WT plants.

### 3.7. The putative role of PPL/PHR2 proteins

PPL/PHR2 proteins are even more cryptic than their sister CRY-DASH clade. The only plant PPL/PHR2 protein characterized so far is CmPHR2 from *Chrysanthemum morifolium* (Zhao et al., 2023). In contrast to GERALT CmPHR2 localizes to nucleus and in complex with CIB (CRYPTOCHROME-INTERACTING bHLH1) regulates transcription. The localization of GERALT in chloroplasts, the disruption of Chl biosynthesis and chloroplast biogenesis in plants lacking this protein indicate that its function is different from that of CmPHR2. This raises the question of the physiological role of other proteins of the PPL/PHR2 subfamily. The shorter FAD binding domain and the lack of conserved residues present in other CPF members, including those involved in substrate recognition (Miles et al., 2020), suggest that the mechanism of action of PPL/PHR2 proteins is different from that described for cryptochromes and photolyases. The severe phenotype is observed in *geralt* mutants grown under standard conditions, without UV irradiation inducing pyrimidine dimer formation. This is another indication ruling out, on the one hand, that the effect observed in the mutant is due to a disruption of photoreactivation, and on the other, that photolyase activity is the primary function performed by the GERALT protein. Our experimental data suggest that GERALT acts as a modulator of chloroplast function, and its role decreases with plant age. They are only a starting point to test whether other members of the plant-specific PPL/PHR2 family also are crucial factors involved in regulation of early stages of seedling development.

## 4. Materials and methods

### 4.1. Plant material and culture conditions

The *Arabidopsis thaliana* WT Col (N6000), SALK_149834 and SALK_116579 seeds were purchased from the Nottingham Arabidopsis Stock Centre (NASC). The seedlings were cultured *in vitro* on B5 medium (Sigma-Aldrich, St Louis, MO, USA) with 1% sucrose (POCH SA, Poland), solidified with 0.7% agar (BTL, Poland). The seeds were surface sterilized (sterile water with a drop of Tween 20 for 5 min., 70% ethanol for 5 min., 50% commercial bleach for 5 min., rinsed three times in distilled, sterile water) and sown on the medium, stratified in a cold room for 2 days and cultured in a growth chamber (23°C, 10hL/14hD photoperiod, at 60–70 µmol·m^−2^·s^−1^ supplied by Sanyo FL40SS.W/37 lamps).

The mutant seeds were collected from plants grown in soil. For this, the mutants were grown *in vitro* for the first 4-5 weeks. The healthy, light green mutant seedlings were transfered to the Jiffy pots (Jiffy Products International AS, Norway). Soil grown plants were cultured in Sanyo MLR 350H chamber (23°C, 10hL/14hD photoperiod, at 60–70 µmol·m^−2^·s^−1^ supplied by Sanyo FL40SS.W/37 lamps, at 80% relative humidity).

SALK_116579 mutant, hereinafter referred to as *geralt*, was used for the research.

### 4.2. Preparation of plasmids

Gateway technology and restriction enzyme cloning method were used to obtain the plasmids with GERALT-GFP under the control of 35S promoter. The lists of the primers and the plasmids used are given in Supplemental Table S4 and S5.

### 4.3 Arabidopsis transformation and transformant selection

For floral dip transformation with *Agrobacterium* the C58 strain carrying pK7FWG2/pH7FWG2/pEarleyGate103:GERALT plasmids the soil grown WT or heterozygous *geralt*+/-plants were used. The transformed plants were selected on media supplemented with kanamycin (50 mg/L), hygromycin B (15 mg/L) or phosphinothricin (PPT; 15 mg/L), depending on the plasmid used. For the hygromycin B selection the protocol described by Harrison et al. (Harrison et al. 2006) was used. All reagents used for the selection were purchased from Sigma-Aldrich, St Louis, MO, USA. The homozygosity of *geralt* mutants both alone (Supplemental Fig. S1) and complemented with the CDS of the *GERALT* gene was confirmed using the primers given in Supplemental Table S4. The WT and homozygous *geralt* lines overexpressing GERALT-GFP fusion protein were marked: WT-OE and *geralt*-OE, respectively. For the experiments one WT-OE (i.e _3E) and 3 independent *geralt*-OE_2F, 3C and 9A lines were used (Supplemental Table S5).

### 4.4. Phenotypic analysis

Germination frequency of WT and *geralt* seeds was assessed at 14 day after sowing. The overall morphology of seedlings and plantlets was assessed at the age of 5, 12 (WT, *geralt* and *geralt*-OE lines) and 42 days (WT, *geralt*). At the same time points photographs were taken using E05 760D camera (Canon, Japan). At least 50 seedlings and 20 plantlets were used per each of three replicate experiments.

### 4.5. Treatment with blue and red light

*In vitro* grown, 6 day-old Arabidopsis seedlings were dark adapted overnight (for around 16 h). The next day, at a time corresponding to 2 h after turning on the light in the culture chamber, they were irradiated for 3 h with 50 µmol·m^−2^·s^−1^ of either blue (470 nm) or red (660 nm) LED light or stayed in darkness. Whole seedlings were then harvested and frozen in liquid nitrogen. The spectra of blue and red light used are given in Supplemental Fig. S11.

### 4.6 RNA isolation and real-time PCR

Spectrum™ Plant Total RNA Kit (Sigma-Aldrich) was used to isolate the RNA with on a column DNA digestion step according to the manufacturer’s protocol (Sigma-Aldrich). The RevertAid M-MuLV Reverse Transcriptase Kit (Thermo Scientific) with oligo(dT)_18_ primers was used to reverse transcribe the mRNA. The real-time PCR experiments were performed as given in Jedynak et al. 2022 (Jedynak et al., 2022) with all reactions run in triplicate. The reference genes: *UBC*, *SAND* and *PDF* were chosen based on Czechowski et al. (Czechowski et al., 2005). GeNorm v3.4 (Vandesompele et al., 2002) was used to calculate the normalization factor for all reference genes used in the experiment. *SAND* and *PDF* genes served as reference genes for the calculation of the relative expression of *GERALT* gene, for the other experiments *SAND* and *UBC* were used as reference genes.

The relative expression of each gene in a sample was determined using the mean value of *Ct* for all samples in the same PCR analysis as a reference. Each PCR analysis included the entire set of samples from one biological repetition (i.e. dark adapted and illuminated with either blue or red light; the whole set of organs tested). The sequences of the primers used in the experiments are given in Supplemental Table S4.

For *LPORA*, *LPORB* and *LPORC* genes, data were logarithmically transformed (log(x)) to ensure homogeneity of variance. One way Anova with a post-hoc TukeyHSD test was performed for the analysis of the influence of blue and red light on the levels of *GERALT* mRNA. Kruskal–Wallis test was used for analysis of *GERALT* expression in plant organs. The results of the expression of chloroplast transcripts and *LPOR* genes were analyzed with two way Anova with a post-hoc TukeyHSD test. The whole statistical analysis was performed using Statistica™ 13.3 software (StatSoft, Inc., Kraków, Poland).

### 4.7 Libraries preparation and RNA-Seq

The whole 5 day-old or 12 day-old grown *in vitro* seedlings were harvested approximately 4h after turn on the light in the culture chamber. The RNA was isolated as described above. Transcriptome libraries were prepared using mRNA-Seq Library Prep kit v2 (Lexogen) according to the manufacturer’s protocol. The libraries insert length was adjusted for 150 read paired-end sequencing. Libraries quality control was performed using 2100 Bioanalyzer (Agilent Technologies Inc.). Twelve libraries were sequenced by CeGaT GmbH, Tübingen, Germany. Six libraries with specific six nucleotides i7 indexes were pooled and used as a sample, two lanes were used. Sequencing was performed using NovaSeq 6000, 2 x 150 nt mode with the quality Q30 value: 90.02%. Adapters were trimmed using Skewer (version 0.2.2) software (Jiang et al., 2014).

### 4.8. RNA-Seq data analysis

Sequencing data analysis was performed using CLC Genomics Workbench 21.0.5 (QIAGEN Aarhus A/S). *Arabidopsis thaliana* reference genome TAIR 10.51 was used. The first nine nucleotides were removed from 5’ end of the both reads. In the next step additional trimming was performed using CLC Genomics Workbench software and following parameters: automatic removal of read-through adapter sequences, removal of low quality sequence (limit = 0.05), removal of 5’ and 3’ adapters, removal of sequences on length: minimum length 50 nt. RNA-Seq analyses for each library were calculated using TPM expression value. Differential expression in two group methods with global TMM normalization was used to calculate the differences in gene expression between *geralt* and WT Arabidopsis plants (control). Bonferroni and FDR p-value correction were applied in the analyses.

STRING database (https://string-db.org/, (Szklarczyk et al., 2023)) was used to visualize potential interactions between proteins encoded by down- and up-regulated genes in 5 or 12-day old *geralt* vs WT seedlings. Active interaction sources (textmining, experiments, databases, co expression and protein co-occurrence), an interaction score > 0.4, species limited to *Arabidopsis thaliana*, DBSCAN clustering with epsilon parameter = 3 were applied to obtain protein-protein interaction network.

### 4.9. Phylogenetic analysis

The NCBI non-redundant protein database was searched using the BLASTp method to find proteins with amino acid sequences similar to that of GERALT. Proteins described as “hypothetical” were removed; in the case of several proteins from a single species, only one was selected. A list of all protein sequences (including the above, plus selected proteins belonging to the CRY-DASH subfamily, CmPHR2 from *Chrysanthemum morifolium* and all Arabidopsis CPF proteins) used to create the phylogenetic tree is provided in Supplementary Data S1. Amino acid sequence of *Chrysanthemum morifolium CmPHR2* (OP889330) was obtained by translation of its nucleotide into amino acid sequence (Zhao et al., 2023). The phylogenetic tree was created using MEGA v. 11.0.13 software (Tamura et al., 2021) with unweighted pair-group method using arithmetic averages (UPGMA), bootstrap value = 100, model/method: Poisson model.

Multiple sequence alignment of GERALT (At2g47590) and selected CRY-DASH proteins was performed using ClustalX1.83 (http://www.clustal.org/) with default parameters and visualized with Jalview 2.07 (Troshin et al., 2011).

### 4.10. Electron microscopy

Leaf fragments (1-2 mm^2^) were fixed in 2.5% (v/v) glutaraldehyde and 2.5% (w/v) paraformaldehyde in 0.05 M cacodylate buffer pH 7.0 for at least 24 h at room temperature. Leaf samples were rinsed in the same buffer and post-fixed in 1% (w/v) osmium tetroxide in a cacodylate buffer at 4°C overnight. Then, material was processed with 1% (w/v) uranyl acetate in distilled water for 1 h, dehydrated in a graded acetone series and finally embedded in Spurr’s epoxy resin (Spurr 1969). Ultrathin, 50-80 nm sections were cut on a Leica EM UC7 ultramicrotome using a diamond knife. Next, the specimens were stained with a saturated solution of uranyl acetate in 50% (w/v) ethanol and with 0.04% (w/v) lead citrate and then viewed using a FEI Tecnai G2 Spirit TWIN transmission electron microscope at 120 kV (Bioimaging Laboratory, Faculty of Biology, University of Gdańsk).

### 4.11. Confocal microscopy

Arabidopsis seedlings were collected and directly placed in a fixing mixture, composed of chloroplast isoosmotic buffer with addition of 1.8% glutaraldehyde and 1% β-mercaptoethanol. Fixed seedlings were stored in a fixing solution in a fridge. For auxiliary DNA visualization, DAPI was added 15 min. prior to imaging to a final concentration of 1 μg/ml. For confocal laser scanning microscopy (CLSM) imaging fixed seedlings were mounted between microscopic glass slides. Imaging was performed using Stellaris confocal system (Leica, Germany) with a tunable pulsed white laser. HC PL Apo CS2 100x/1.4 oil objective was used. Emission ranges were set with a monochromator. Chl fluorescence was excited with a white pulse laser set at 470 nm. The emission was collected at 650-750 nm. GFP emission was excited with the same laser line and emission was collected at 490-540 nm. For an auxiliary DAPI (a dye used for DNA staining) excitation 405 nm laser diode was used, and emission was collected at 420-480 nm. The images were collected using LasX software (Leica, Germany). For better visualization image contrast was adjusted as needed, and - if necessary for clarity of the presentation - only a zoomed part of image was shown. Each time the zoom-in option was used, it is indicated in the figure legend. The original, unmodified versions of all images are available upon request. Some images in supplemental file (indicated in the figure legend) were obtained using Sp8 confocal system (Leica, Germany) with HC PL Apo CS2 63x/1.4 oil objective, with 405 nm/488 nm laser excitation and the same emission ranges as described above.

For high resolution imaging fixed seedling were mounted between microscopic glass slides and imaged, using the Elyra7 confocal system (Zeiss, Germany), equipped with HC PL Apo 100x/1.4 oil objective. Fluorescence of Chl was excited with a 488 nm laser diode, and emission was collected with a long pass 590 nm filter. For additional DAPI channel, 405 nm excitation and 420-480 nm emission band pass filter was applied. GFP emission was imaged with a 488 nm laser diode excitation and 490-550 nm emission band pass filter. SIM^2^ (structured illumination microscopy) processing, increasing resolution to 60 nm, was done in Zen Black (Zeiss, Germany) dedicated software. No additional processing was applied except contrast adjustment for easier reading. For particular figures the image contrast was adjusted, and - if necessary for clarity of presentation - only a part of the image was shown. If a zoom-in option was used to produce the figure, it is indicated in its legend. The original, unmodified images are available upon request.

For quantification of chloroplast size, their major axis were determined using Fiji (Schindelin et al., 2012) and AnalyseParticle plugin. Chloroplasts were identified manually in at least 6 separate images, collected from at least 3 independent seedlings. The total number of chloroplasts used for analysis is indicated in the figure legend. For studies of grana stacking thickness, at least 8 representative SIM^2^ chloroplast images were analyzed, using Fiji and LocalThickness plugin (Saito and Toriwaki, 1994; Hildebrand and Rüegsegger, 1997). 3D projections presented as Supplemental movies were constructed with 3D viewer plugin of Fiji software (Schmid et al., 2010).

For statistical analysis one way Anova with a post-hoc TukeyHSD test were performed using R (R Core Team).

### 4.12. Chloroplast isolation, fractionation and Western blot analysis

Chloroplasts were isolated from approximately 8 g of leaves from fully grown rosettes. The tissue was crushed in isolation buffer (330 mM sorbitol, 50 mM HEPES-KOH pH 8.0, 5 mM EGTA, 1 mM MgCl_2_, 10 mM NaHCO_3_, 5 mM sodium ascorbate) on ice and filtered through 2 layers of Miracloth (Merck Millipore). Intact chloroplasts were collected from a band at the interface between the 40 % (v/v) and 80 % (v/v) Percoll layers after centrifugation at 1800 × g for 20 min. at 4°C. Chloroplasts were washed with isolation buffer and collected for further analyses or lysed with osmotic lysis buffer (20 mM HEPES-KOH pH 8.0, 5 mM MgCl_2_, tablet of Roche Protease Inhibitor Cocktail). After centrifugation 15000 x g for 15 min. at 4°C the supernatant (stroma) and pellet (thylakoids) fractions were recovered. Stroma proteins were concentrated with Amicon Ultra Centrifugal Filters (Merck Millipore). The protein content was estimated using Bradford Assay Kit (Thermo Fisher Scientific, USA). Protein samples were solubilized in Laemmli’s buffer (Laemmli 1970) and heated at 95°C for 5 min. at 4°C (stroma fraction) or at 70°C for 5 min. at 4°C (chloroplasts and thylakoids). Chloroplast, thylakoid and stroma proteins were separated by SDS PAGE using 12 % polyacrylamide gels. The fractionated proteins were electroblotted on nitrocelulose membrane, stained with Pierce™ Reversible Protein Stain Kit (Thermo Fisher Scientific,) to ensure proper transfer and blocked with 5 % (w/v) non-fat milk powder. Immunodetections were performed using specific antibodies against GFP (sc-9996, Santa Cruz Biotechnology, USA) at dilution of 1:1000, against LHCII type chlorophyll a/b binding proteins (anti-Lhcb2, AS01 003, Agrisera, Sweden) at dilution of 1:5000 and against Rubisco large subunit (anti-RbcL, AS03 037, Agrisera, Sweden) at dilution of 1:10000. Immunodetection was performed by the chemiluminescence method.

### 4.13. Pigment composition

For pigment analysis 7 or 15 day old WT and *geralt* seedlings were harvested, frozen in liquid nitrogen and kept at -80°C until used. Extraction was performed under dim light. Frozen seedlings were ground in small portions of extraction solvent (solvent A). Extracts were pooled, centrifuged and filtered through 0.2 μm PTFE syringe filter (Thermo Scientific). Pigment composition was analyzed by HPLC using PU-2089 Plus system (JASCO) coupled with a UV-VIS detector (MD-2015 Plus, JASCO), using Agilent Zorbax 300SB-C18 column (4.6 x 250 mm, 5 µm particle size, Agilent). Samples were loaded onto a column equilibrated with Solvent A (acetonitrile:methanol:water, 72:8:1, v:v:v) and isocratic elution was performed for 22 min., then solvent was changed during steep 1-min. gradient to 100% solvent B (methanol : ethyl acetate, 34:16, v:v) for 4 min., followed by a 5 min. isocratic hold at 100% A. Flow rate was 0.8 ml min^−1^ for the first 22 min., then it increased to 1.7 ml min.^-1^ for 9 min., and decreased to 0.8·ml min.^−1^ for the last 2 min. The absorption spectra of the eluate (300–800 nm) were recorded every 0.6 s. Carotenoids were identified on the basis of absorption spectra and retention times. Amount of each xanthophyll was determined based on the maximum absorbance peak area. The concentrations of carotenoid pigments were calculated from Beer–Lambert law using specific extinction coefficients. Values are means ± SE calculated from at least three independent biological repetitions.

For statistical analysis one way Anova with a post-hoc TukeyHSD was performed using R (R Core Team).

### 4.14. Photosynthetic efficiency

Maximum PSII quantum yield (F_V_/F_M_)(Genty et al. 1989) was imaged using a pulse-modulated Open Fluor-Cam FC 800-O/1010 fluorimeter and FluorCam7 software (PSI, Drasov, Czech Republic). An additional lens (M-55 Canon) was placed in the front of the camera, allowing a 2-2.5 times magnification of the images of the examined seedlings. The intensity of the saturating pulse was set at 3700 μmol·m^-2^·s^-1^. Chl fluorescence measurements were performed on 5- and 12-day-old seedlings of WT or *geralt* grown on Ø 60 mm Petri dishes, with approximately 10-20 seedlings per dish. Each dish was pre-incubated in darkness for 10 min. immediately before measurement. Image and setting optimisation was performed before darkening. The dishes remained covered all the time to prevent the plants from drying out (the lid does not interfere with fluorescence measurements). Each experiment was repeated independently 3 times and the average values of F_V_/F_M_=(F_M_-F_0_)/F_M_ along with respective standard deviations, were calculated.

### 4.15. Gene Ontology (GO) analysis

Changes in gene expression were analyzed using Differential Expression in Two Groups method included in CLC Genomics Workbench 21.0.5 software (QIAGEN Aarhus A/S). One group was composed of SALK_116579 mutant, (*phr2*, *geralt* plants) and the second was represented by wild type *Arabidopsis thaliana* Col (N6000) plants. The RNA-Seq analyses were performed for two group of plants (each represented by three biological replication) at two time points (i) 5 day-old and (ii) 12 day-old seedlings. In total 27,655 genes were analyzed. Gene Ontology (GO) analyses were performed for genes with fold changes significantly with either p-value <= 0.05 or Bonferroni corrected p-value <= 0.05. There are 4 702 and 417 genes which expression was changed significantly with p-value and Bonferroni corrected p-value <= 0.05 for 5 day-old seedlings, respectively. There are 2 663 and 125 genes which expression was changed significantly with p-value and Bonferroni corrected p-value <= 0.05 for 12 day-old seedlings, respectively. GO was carried out using “AnnotationDbi”, “org.At.tair.db”, “biomaRT”, “clusterProfiler” packages in R. GO results are represented by two categories (a) denoted as GO Biological Process with first 10 gene categories and (b) denoted as GO Molecular Function with 10 activity categories.

## Abbreviations

Chl: chlorophyll Chlide - chlorophyllide
CLSM: confocal laser scanning microscopy
CPD: cyclobutane pyrimidine dimer
CPF: cryptochrome/photolyase family
CRY: cryptochrome
CRY-DASH: *Drosophila, Arabidopsis, Synechocystis*, human-type cryptochrome
DEG: differentially expressed gene
FAD: flavin adenine dinucleotide
GO: Gene Ontology
8-HDF: 8-hydroxy-7,8-didemethyl-5-deazariboflavin
LPOR: light dependent protochlorophyllide oxidoreductase
MTHF: methenyltetrahydrofolate
NEP: nuclear encoded RNA polymerase Pchlide - protochlorophyllide
PEP: plastid-encoded RNA polymerase
PPL: plant photolyase
PHR2: photolyase/blue-light receptor 2
SIM: structured illumination microscopy
TEM: transmission electron microscopy

## Supplementary data

**Supplementary Material 1: Supplementary Figures S1-S11, Supplementary Tables S1-S5,**

## Supplementary Figures

**Fig. S1.** GERALT gene structure and mutant genotyping.

**Fig. S2.** Phenotype of *geralt* mutants.

**Fig. S3**. Phenotypes of *geralt*-OE-2F/3C/9A lines at 5 and 12 days after sowing

**Fig. S4**. CLSM images of WT and *geralt* chloroplasts.

**Fig. S5.** Comparison of chloroplast population size in 5 day-old and 12 day-old seedlings of WT, *geralt*, WT-OE_3E_ and complementant geralt-OE_9A/2F/3C_ lines.

**Fig. S6.** The maximum PSII quantum yield for plants overexpressing *GERALT*

**Fig. S7.** Gene ontology (GO) analysis of 5 day-old and 12 day-old *geralt* and WT seedlings (p-value).

**Fig. S8.** Localization of GERALT-GFP protein in chloroplasts.

**Fig. S9.** Alignment of GERALT (At2g47590) and selected CRY-DASH proteins.

**Fig. S10.** Maps of interactions between proteins encoded by genes down-or down-regulated in *geralt* seedlings visualised using STRING database (https://string-db.org).

**Fig. S11**. Spectra of blue (470 nm) and red (660 nm) light used in the experiments.

## Supplementary Tables

**Table S1.** List of DEGs identified in both 5 and 12 day-old seedlings

**Table S2.** Selected photosynthesis-associated genes differentially expressed in *geralt* mutant

**Table S3.** The selected transcription factor binding sites identified in the *GERALT* promoter

**Table S4**. The list of primers used for cloning, genotyping and real-time PCR.

**Table S5.** The list of vectors used for *Arabidopsis* transformation

**Supplementary Excel file S1.** List of genes which expression differ over 2-fold (p-value Bonferroni correction ≤ 0.05) between 5 days old and 12 day-old *geralt* and WT seedlings.

## Supplementary Video

**Supplementary Video_S1. Supplementary file wt_3D_Z-stack - Collection of Z-stack images (combined as a movie) of mesophyll chloroplasts of WT plants**. Images obtained by CLSM technique, are overlay of chlorophyll (red) and DAPI (blue) emission. Scale bar is 20 μm.

**Supplementary Video_S2. Supplementary file Geralt_3D_Z-stack - Collection of Z-stack images (combined as a movie) of mesophyll chloroplasts of *geralt* plants.** Images obtained by CLSM technique, are overlay of chlorophyll (red) and DAPI (blue) emission. Scale bar is 20 μm.

**Supplementary Video_S3. WT_3D_Z-stack** - **Z-stack image (combined as a movie) of selected mesophyll chloroplasts of WT plants**. Images obtained by SIM^2^ technique (subfiles: 1_3D to 3_3D), are overlay of chlorophyll (red) and DAPI (blue) emissions.

**Supplementary Video_S4. geralt_3D_Z-stack** - Z-stack image (combined as a movie) of selected mesophyll chloroplasts of *geralt* plants. Images obtained by SIM^2^ technique (subfiles: 1_3D to 4_3D), are overlay of chlorophyll (red) and DAPI (blue) emissions.

**Supplementary Video_S5. Geralt_GFP_3D_Z-stack** - **Z-stack image (combined as a movie) of selected pavement chloroplast of WT-OE_3E_ plants**. Images obtained by SIM^2^ technique (subfile: Geralt_GF_Z-stack1), are overlay of chlorophyll (red), GFP (green) and DAPI (blue) emissions. Additionally, free hand rotations of 3D projections, based on that Z-stacks are presented for all three channels overlay (subfile: Geralt_GFP_Z-stack_movie1) and GFP channel only (subfile: Geralt_GFP_Z-stack_Gfp_movie). 3D projection was constructed with 3D viewer plugin of Fiji software (Schmid et al., 2010). No PSF correction was applied to the Z-stacks.

## Supplementary data

**Supplementary data S1**. Amino acid sequences of proteins used to construct a phylogenetic tree.

## Acknowledgements

We are grateful to Bo Hong (China Agriculture University, Beijing, China) for providing *CmPHR2* nucleotide sequence, Dawid Bielewicz (Center for Advanced Technologies, Adam Mickiewicz University, Poznań, Poland) for helping in RNA-Seq data deposition in the GEO database and Justyna Łabuz (Malopolska Centre of Biotechnology, Jagiellonian University, Krakow, Poland) for preparation of pDONR with *GERALT* CDS sequence.

## Author contributions

AKB – Conceptualization, Funding acquisition, Investigation, Project administration, Writing – original draft; JG – Conceptualisation, Investigation, Visualization, Resources, Writing – original draft; PZ – Investigation, Plant transformation, Selection of transgenic lines; AP – RNA-Seq libraries construction, RNA-Seq data analysis, GO data analysis, Visualization, Writing – original draft; BMK – Investigation, Methodology, Analysis, Visualization and Writing – original draft; KL – Investigation, Formal analysis, Visualization, Writing – review & editing; M.K-K. – TEM analysis of the ultrastructure of the chloroplasts, Visualization; KK– Investigation; RK – Investigation, Visualization, Writing – original draft; MP – Western blot analyses, Visualization; EN – Involvement in Conceptualization and in Writing of original draft, AB – Investigation; ŁSz – Gene data preparation, Writing R-scripts used for GO analysis, GO data analyses; EBK - Investigation, WS – Investigation.

## Data availability

All data supporting the findings of this study are available within the paper and within its supplementary data published online. Raw data can be obtained from the corresponding author under reasonable request.

## Accession numbers

Accession numbers (TAIR, www.arabidopsis.org) of analyzed genes: *GERALT/PHR2 (At2g47590); psbB (AtCg00680); psaB (AtCg00340); psbD (AtCg00270); RbcL (AtCg00490); accD (AtCg00500); rpoA (AtCg00740); rpoC1 (AtCg00180); rps18 (AtCg00650); atpB (AtCg00480); clpP (AtCg00670); LPORA (At5g54190); LPORB (At4g27440); LPORC (At1g03630).* NCBI accession numbers of proteins used for construction of phylogenetic tree are given in Supplementary data S1. The RNA-seq data discussed in this publication have been deposited in NCBI’s Gene Expression Omnibus, GEO (Edgar et al., 2002) and are accessible through GEO Series accession number GSE271033 (https://www.ncbi.nlm.nih.gov/geo/query/acc.cgi?acc= GSE271033).

## References

Bjőrkman O and Demmig B. Photon yield of 0 2 evolution and chlorophyll fluorescence characteristics at 77 K among vascular plants of diverse origins. Planta 1987:170:489– 504.

Brawley SH, Blouin NA, Ficko-Blean E, Wheeler GL, Lohr M, Goodson H V., Jenkins JW, Blaby-Haas CE, Helliwell KE, Chan CX, et al. Insights into the red algae and eukaryotic evolution from the genome of Porphyra umbilicalis (Bangiophyceae, Rhodophyta). Proc Natl Acad Sci U S A. 2017:114(31):E6361–E6370. 10.1073/pnas.1703088114

Charuvi D, Kiss V, Nevo R, Shimoni E, Adam Z, and Reich Z. Gain and loss of photosynthetic membranes during plastid differentiation in the shoot apex of arabidopsis. Plant Cell. 2012:24(3):1143–1157. 10.1105/tpc.111.094458

Chen M, Jensen M, and Rodermel S. The yellow variegated Mutant of Arabidopsis Is Plastid Autonomous and Delayed in Chloroplast Biogenesis. J Hered. 1999:90:207–214.

Czechowski T, Stitt M, Altmann T, Udvardi MK, and Scheible WR. Genome-wide identification and testing of superior reference genes for transcript normalization in arabidopsis. Plant Physiol. 2005:139(1). 10.1104/pp.105.063743

Davuluri R V, Sun H, Palaniswamy SK, Matthews N, Molina C, Kurtz M, and Grotewold E. AGRIS: Arabidopsis Gene Regulatory Information Server, an information resource of Arabidopsis cis-regulatory elements and transcription factors. BMC Bioinformatics. 2003:4:25.

Deppisch P, Helfrich-Förster C, and Senthilan PR. The Gain and Loss of Cryptochrome/Photolyase Family Members during Evolution. Genes (Basel). 2022:13(9). 10.3390/genes13091613

Edgar R, Domrachev M, and Lash AE. Gene Expression Omnibus: NCBI gene expression and hybridization array data repository. Nucleic Acids Res. 2002:30(1):207–210.

Gabruk M and Mysliwa-Kurdziel B. Light-Dependent Protochlorophyllide Oxidoreductase: Phylogeny, Regulation, and Catalytic Properties. Biochemistry. 2015:54(34):5255–5262. 10.1021/acs.biochem.5b00704

Genty B, Briantais J-M, and Baker NR. The relationship between the quantum yield of photosynthetic electron transport and quenching of chlorophyll fluorescence. Biochimica et Biophysica Acta (BBA) - General Subjects. 1989:990(1):87–92.

Hajdukiewicz PTJ and Allison LA. The two RNA polymerases encoded by the nuclear and the plastid compartments transcribe distinct groups of genes in tobacco plastids. EMBO J. 1997:16:4041–4048.

Harrison SJ, Mott EK, Parsley K, Aspinall S, Gray JC, and Cottage A. A rapid and robust method of identifying transformed Arabidopsis thaliana seedlings following floral dip transformation. Plant Methods. 2006:2(1). 10.1186/1746-4811-2-19

Hildebrand T and Rüegsegger P. A new method for the model-independent assessment of thickness in three-dimensional images. J Microsc. 1997:185(1):67–75. 10.1046/j.1365-2818.1997.1340694.x

Hooper CM, Castleden IR, Tanz SK, Aryamanesh N, and Millar AH. SUBA4: The interactive data analysis centre for Arabidopsis subcellular protein locations. Nucleic Acids Res. 2017:45(D1):D1064–D1074. 10.1093/nar/gkw1041

Hricová A, Quesada V, and Micol JL. The SCABRA3 nuclear gene encodes the plastid RpoTp RNA polymerase, which is required for chloroplast biogenesis and mesophyll cell proliferation in Arabidopsis. Plant Physiol. 2006:141(3):942–956. 10.1104/pp.106.080069

Iwai M, Roth MS, and Niyogi KK. Subdiffraction-resolution live-cell imaging for visualizing thylakoid membranes. Plant Journal. 2018:96(1):233–243. 10.1111/tpj.14021

Jedynak P, Trzebuniak KF, Chowaniec M, Zgłobicki P, Banaś AK, and Mysliwa-Kurdziel B. Dynamics of Etiolation Monitored by Seedling Morphology, Carotenoid Composition, Antioxidant Level, and Photoactivity of Protochlorophyllide in Arabidopsis thaliana. Front Plant Sci. 2022:12. 10.3389/fpls.2021.772727

Jiang CZ, Yee J, Mitchell DL, and Britt AB. Photorepair mutants of arabidopsis. Proc Natl Acad Sci U S A. 1997:94(14). 10.1073/pnas.94.14.7441

Jiang H, Lei R, Ding S-W, and Zhu S. Skewer: a fast and accurate adapter trimmer for next-vs sequencing paired-end reads. BMC Bioinformatics. 2014:15:182

Kiontke S, Göbel T, Brych A, and Batschauer A. DASH-type cryptochromes - Solved and open questions. Biol Chem. 2020:401(12):1487–1493. 10.1515/hsz-2020-0182

Kleine T, Lockhart P, and Batschauer A. An Arabidopsis protein closely related to Synechocystis cryptochrome is targeted to organelles. Plant Journal. 2003:35(1):93–103. 10.1046/j.1365-313X.2003.01787.x

Laemmli U. Cleavage of structural proteins during assembly of head of bacteriophage-T4. Nature. 1970:227:680–685.

Lescot M, Déhais P, Thijs G, Marchal K, Moreau Y, Van De Peer Y, Rouzé P, and Rombauts S. PlantCARE, a database of plant cis-acting regulatory elements and a portal to tools for in silico analysis of promoter sequences. Nucleic Acids Res. 2002:30:325–327.

Liang Z, Zhu N, Mai KK, Liu ZY, Tzeng D, Osteryoung KW, Zhong S, Staehelin LA, and Kang BH. Thylakoid-bound polysomes and a dynamin-related protein, FZL, mediate critical stages of the linear chloroplast biogenesis program in greening arabidopsis cotyledons. Plant Cell. 2018:30(7):1476–1495. 10.1105/tpc.17.00972

Liu D, Li W, and Cheng J. The novel protein DELAYED PALE-GREENING1 is required for early chloroplast biogenesis in Arabidopsis thaliana. Sci Rep. 2016:6. 10.1038/srep25742

Lucas-Lledó JI and Lynch M. Evolution of mutation rates: Phylogenomic analysis of the photolyase/cryptochrome family. Mol Biol Evol. 2009:26(5):1143–1153. 10.1093/molbev/msp029

Mei Q and Dvornyk V. Evolutionary history of the photolyase/cryptochrome superfamily in eukaryotes. PLoS One. 2015:10(9). 10.1371/journal.pone.0135940

Miles JA, Davies TA, Hayman RD, Lorenzen G, Taylor J, Anjarwalla M, Allen SJR, Graham JWD, and Taylor PC. A Case Study of Eukaryogenesis: The Evolution of Photoreception by Photolyase/Cryptochrome Proteins. J Mol Evol. 2020:88(8–9):662–673. 10.1007/s00239-020-09965-x

Oldemeyer S, Haddad AZ, and Fleming GR. Interconnection of the Antenna Pigment 8-HDF and Flavin Facilitates Red-Light Reception in a Bifunctional Animal-like Cryptochrome. Biochemistry. 2020:59(4):594–604. 10.1021/acs.biochem.9b00875

Paik I and Huq E. Plant photoreceptors: Multi-functional sensory proteins and their signaling networks. Semin Cell Dev Biol. 2019:92:114–121. 10.1016/j.semcdb.2019.03.007

Paysan-Lafosse T, Blum M, Chuguransky S, Grego T, Pinto BL, Salazar GA, Bileschi ML, Bork P, Bridge A, Colwell L, et al. InterPro in 2022. Nucleic Acids Res. 2023:51(D1):D418–D427. 10.1093/nar/gkac993

Petersen JL, Lang DW, and Small GD. Cloning and characterization of a class II DNA photolyase from Chlamydomonas.

Pfalz J and Pfannschmidt T. Essential nucleoid proteins in early chloroplast development. Trends Plant Sci. 2013:18(4):186–194. 10.1016/j.tplants.2012.11.003

Pogson BJ, Ganguly D, and Albrecht-Borth V. Insights into chloroplast biogenesis and development. Biochim Biophys Acta Bioenerg. 2015:1847(9):1017–1024. 10.1016/j.bbabio.2015.02.003

Pokorny R, Klar T, Hennecke U, Carell T, Batschauer A, and Essen L-O. Recognition and repair of UV lesions in loop structures of duplex DNA by DASH-type cryptochrome. Proc Natl Acad Sci U S A. 2008:105:21023–21027.

R Core Team. R: A Language and Environment for Statistical Computing. R Foundation for Statistical Computing, Vienna. 2021. https://www.R-project.org.

Rizzini L, Favory JJ, Cloix C, Faggionato D, O’Hara A, Kaiserli E, Baumeister R, Schäfer E, Nagy F, Jenkins GI, et al. Perception of UV-B by the arabidopsis UVR8 protein. Science. 2011:332(6025). 10.1126/science.1200660

Rodríguez-Alcocer E, Ruiz-Pérez E, Parreño R, Martínez-Guardiola C, Berna JM, Çakmak Pehlivanlı A, Jover-Gil S, and Candela H. Cloning of an Albino Mutation of Arabidopsis thaliana Using Mapping-by-Sequencing. Int J Mol Sci. 2023:24(4). 10.3390/ijms24044196

Ruppel NJ, Logsdon CA, Whippo CW, Inoue K, and Hangarter RP. A mutation in arabidopsis seedling plastid development1 affects plastid differentiation in embryo-derived tissues during seedling growth. Plant Physiol. 2011:155(1):342–353. 10.1104/pp.110.161414

Saito T and Toriwaki J-I. New algorithms for euclidean distance transformation of an n-dimensional digitized picture with applications. Pattern Recognition 1994:27:1551–1565.

Sancar A. Structure and function of DNA photolyase and cryptochrome blue-light photoreceptors. Chem Rev. 2003:103(6):2203–2237. 10.1021/cr0204348

Schindelin J, Arganda-Carreras I, Frise E, Kaynig V, Longair M, Pietzsch T, Preibisch S, Rueden C, Saalfeld S, Schmid B, et al. Fiji: An open-source platform for biological-image analysis. Nat Methods. 2012:9(7):676–682. 10.1038/nmeth.2019

Schmid B, Schindelin J, Cardona A, Longair M, and Heisenberg M. Open Access SOFTWARE A high-level 3D visualization API for Java and ImageJ. BMC Bioinformatics. 2010:11:274.

Shimada H, Mochizuki M, Ogura K, Froehlich JE, Osteryoung KW, Shirano Y, Shibata D, Masuda S, Mori K, and Takamiya KI. Arabidopsis cotyledon-specific chloroplast biogenesis factor CYO1 is a protein disulfide isomerase. Plant Cell. 2007:19(10):3157– 3169. 10.1105/tpc.107.051714

Shi Y, Chen J, and Hou X. Similarities and Differences of Photosynthesis Establishment Related mRNAs and Novel lncRNAs in Early Seedlings (Coleoptile/Cotyledon vs. True Leaf) of Rice and Arabidopsis. Front Genet. 2020:11. 10.3389/fgene.2020.565006

Sidorczuk K, Gagat P, Kała J, Nielsen H, Pietluch F, Mackiewicz P, and Burdukiewicz M. Prediction of protein subplastid localization and origin with PlastoGram. Sci Rep. 2023:13(1). 10.1038/s41598-023-35296-0

Spurr AR. A Low-Viscosity Epoxy Resin Embedding Medium for Electron Microscopy. J Ultra Mol Struct R. 1969:26:31–43.

Szklarczyk D, Kirsch R, Koutrouli M, Nastou K, Mehryary F, Hachilif R, Gable AL, Fang T, Doncheva NT, Pyysalo S, et al. The STRING database in 2023: protein-protein association networks and functional enrichment analyses for any sequenced genome of interest. Nucleic Acids Res. 2023:51(1 D):D638–D646. 10.1093/nar/gkac1000

Tadini L, Jeran N, Peracchio C, Masiero S, Colombo M, and Pesaresi P. The plastid transcription machinery and its coordination with the expression of nuclear genome: Plastid-encoded polymerase, nuclear-encoded polymerase and the genomes uncoupled 1-mediated retrograde communication. Philosophical Transactions of the Royal Society B: Biological Sciences. 2020:375(1801). 10.1098/rstb.2019.0399

Tamura K, Stecher G, and Kumar S. MEGA11: Molecular Evolutionary Genetics Analysis Version 11. Mol Biol Evol. 2021:38(7):3022–3027. 10.1093/molbev/msab120

Trotta A, Bajwa AA, Mancini I, Paakkarinen V, Pribil M, and Aro EM. The role of phosphorylation dynamics of curvature thylakoid 1B in plant thylakoid membranes1[open]. Plant Physiol. 2019:181(4):1615–1631. 10.1104/pp.19.00942

Vandesompele J, De Preter K, Pattyn F, Poppe B, Van Roy N, De Paepe A, and Speleman F. Accurate normalization of real-time quantitative RT-PCR data by geometric averaging of multiple internal control genes. Genome Biol. 2002:3(7). 10.1186/gb-2002-3-7-research0034

Wang Q and Lin C. Mechanisms of Cryptochrome-Mediated Photoresponses in Plants. Annu Rev Plant Biol. 2020:29(71):103–1292020. 10.1146/annurev-arplant-050718

Wang X, An Y, Qi Z, and Xiao J. PPR protein Early Chloroplast Development 2 is essential for chloroplast development at the early stage of Arabidopsis development. Plant Science. 2021:308. 10.1016/j.plantsci.2021.110908

Xu L, Chen S, Wen B, Shi H, Chi C, Liu C, Wang K, Tao X, Wang M, Lv J, et al. Identification of a Novel Class of Photolyases as Possible Ancestors of Their Family. Mol Biol Evol. 2021:38(10). 10.1093/molbev/msab191

Yamamoto YY, Puente P, and Deng X-W. An Arabidopsis Cotyledon-Specific Albino Locus: a Possible Role in 16S rRNA Maturation. Plant Cell Physiol. 2020:41:68–76.

Yoo CY, Pasoreck EK, Wang H, Cao J, Blaha GM, Weigel D, and Chen M. Phytochrome activates the plastid-encoded RNA polymerase for chloroplast biogenesis via nucleus-to-plastid signaling. Nat Commun. 2019:10(1). 10.1038/s41467-019-10518-0

Zhao X, Liu W, Aiwaili P, Zhang H, Xu Y, Gu Z, Gao J, and Hong B. PHOTOLYASE/BLUE LIGHT RECEPTOR2 regulates chrysanthemum flowering by compensating for gibberellin perception. Plant Physiol. 2023:193(4):2848–2864. 10.1093/plphys/kiad503

