## Supplementary Material 1: Supplementary Figures S1-S11, Supplementary Tables S1-S5, Supplementary data S1 for "Toss GERALT into chloroplast to make it green: an age-dependent regulator of chloroplast biogenesis and chlorophyll biosynthesis"

**Supplemental Figures**

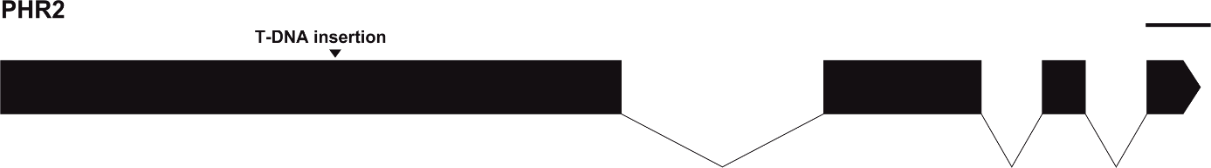

A)

B)

C)

100 bp

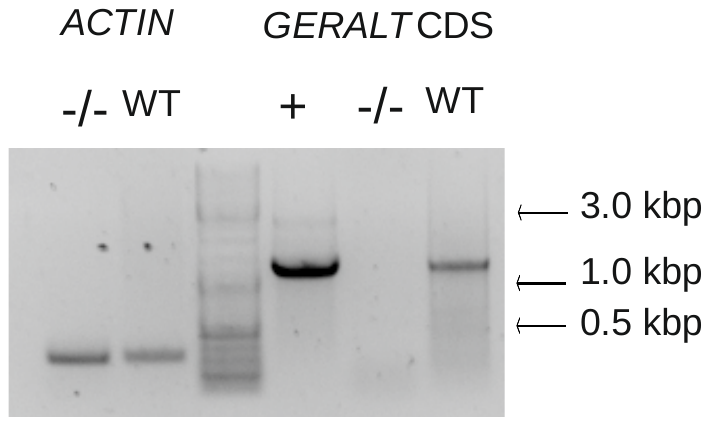

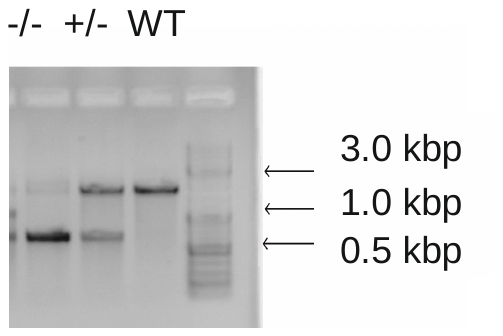

D)

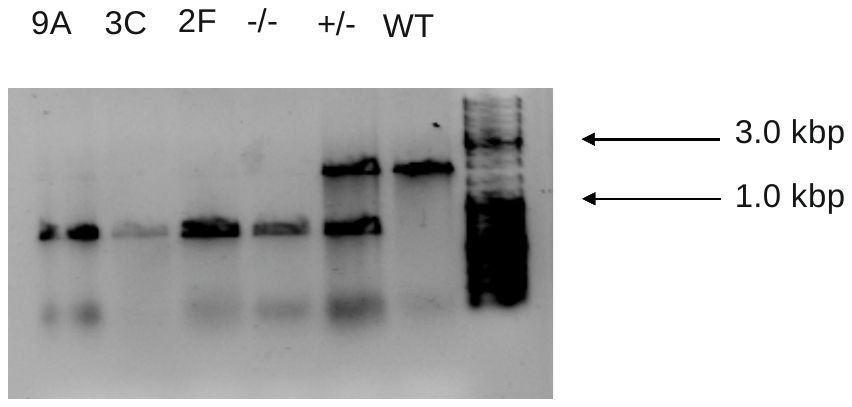

**Fig. S1.** GERALT gene structure and mutant genotyping. A) A scheme of GERALT gene (with indicated T-DNA insertion) prepared with webtool Exon-Intron Graphic Maker (wormweb.org). B) Genotyping of the T-DNA insertion line (*geralt*). B) Genotyping of *geralt* T-DNA and (D) complemented *geralt*-OE2F, 3C and 9A lines with geralt_genot_For/ geralt_genot_Rev/Lba1 primers. C) The expression of *GERALT* CDS amplified with Geralt_F/R and actin F/R primers. WT Arabidopsis, -/- – *geralt* mutant, +/- - heterozygous plant, + - positive control – plasmid PK7FWG2:GERALT. The primers used for the analyses are listed as in Supplemental **Table S1**

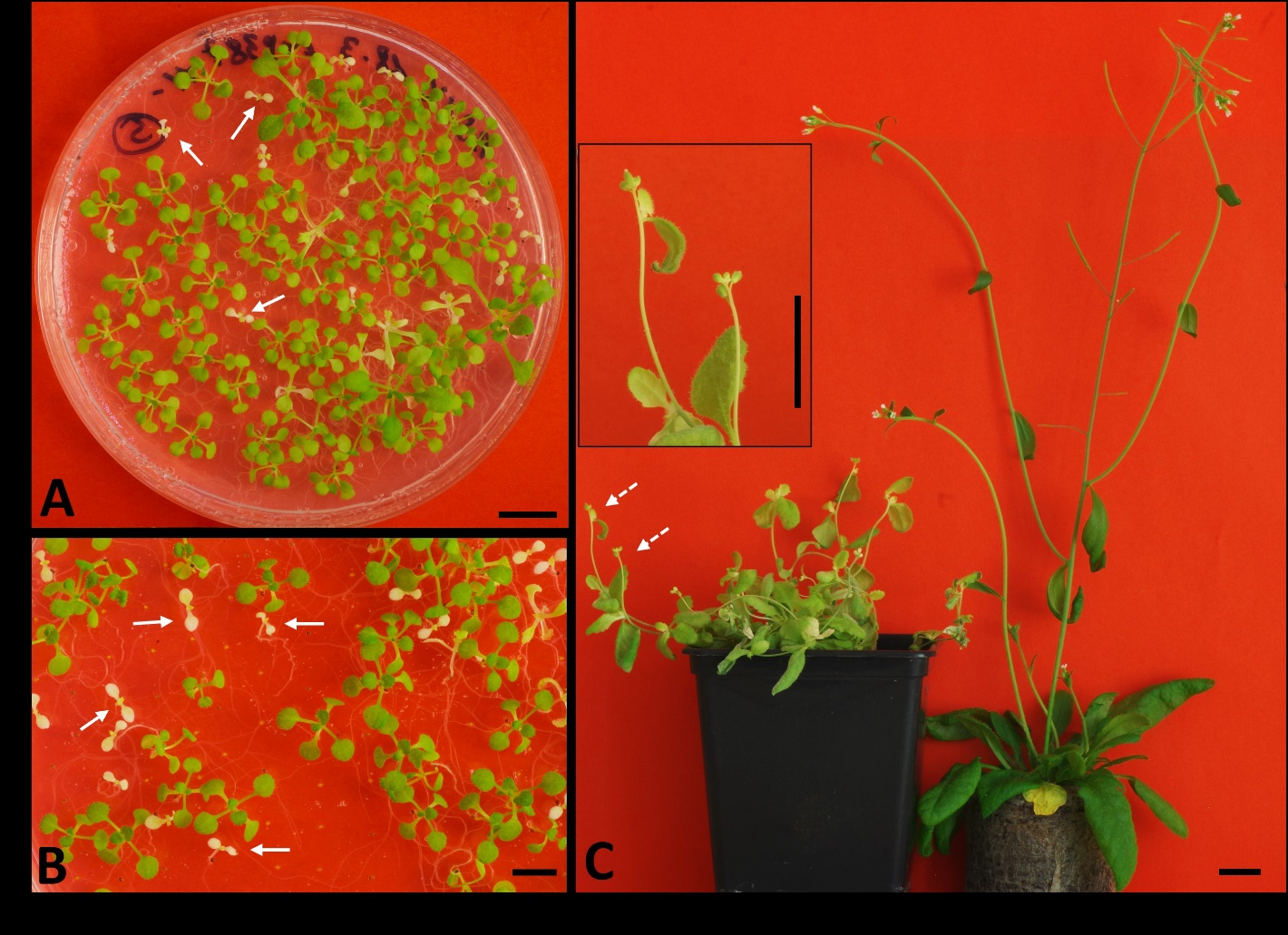

**Fig. S2**. Images of mutants in GERALT gene. The images of 18 day-old progenies of heterozygous (A) SALK_149834 and (B) *geralt* (SALK_116579) plants. The examples of homozygous *geralt* plants are marked with arrows. (C) The images of flowering *geralt* (left) and WT (right) plants. Dashed arrows indicate inflorescences on *geralt,* shown in higher magnification in the inset. Bar represents 10 mm.

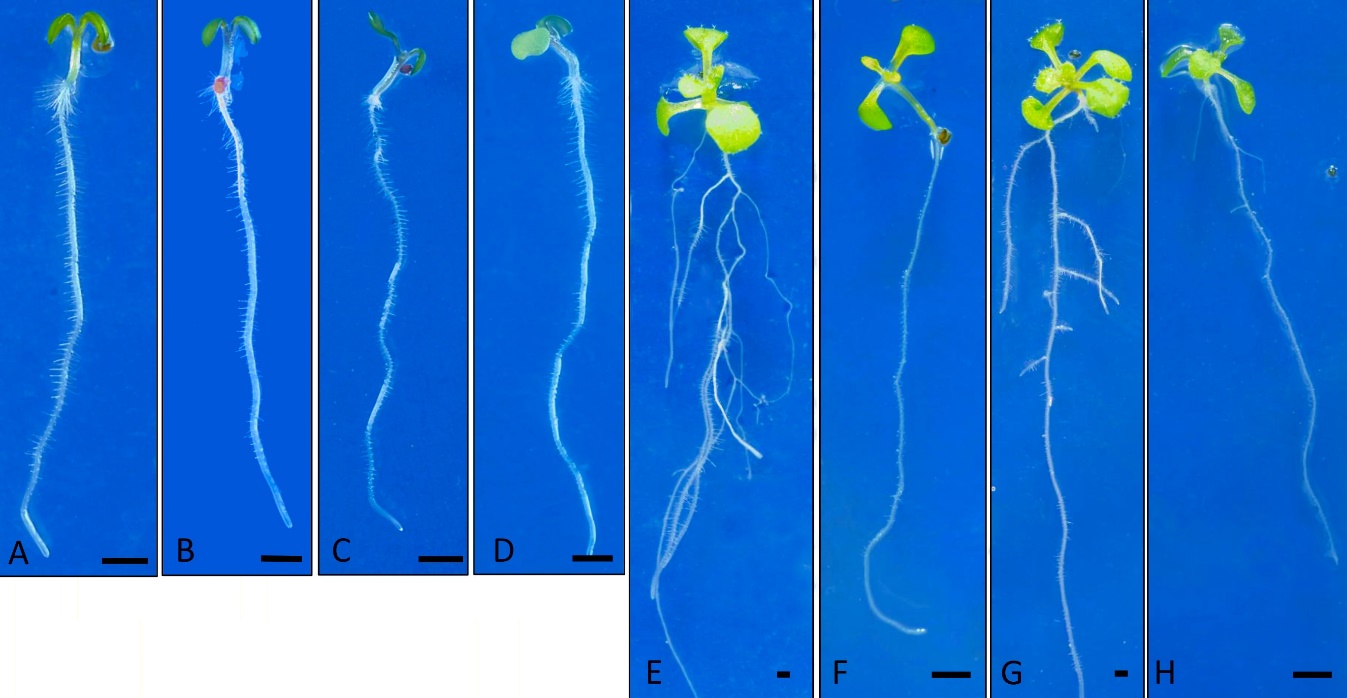

**Fig. S3.** Phenotypes of *geralt*-OE-2F/3C/9A lines at 5 (A-D) and 12 (E-H) days after sowing. The seedling lines: 2F (A, E), 3C (B, F), 9A (C, G) and 3E (D, H). Bar represents 1mm.

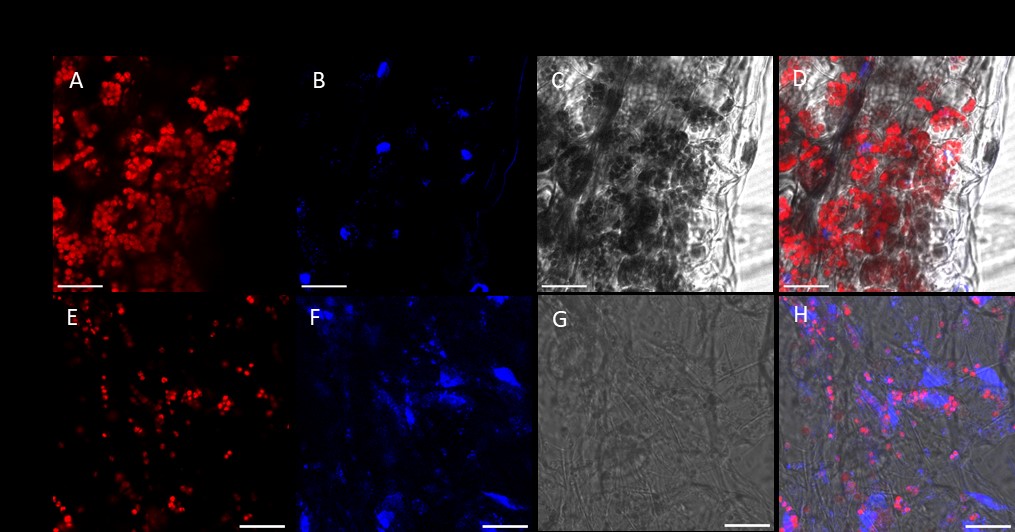

**Fig. S4**. CLSM images of WT and *geralt* chloroplasts**.** Example of CLSM images of WT (A-D) and *geralt* (E-H) chloroplasts from cotyledons of 5 day-old fixed seedlings. Chlorophyll emission (red - A, E), DAPI emission (blue - B, F), transmission channel (G, H) and overlay of all channels (D, H). Scale bar = 20 μm.

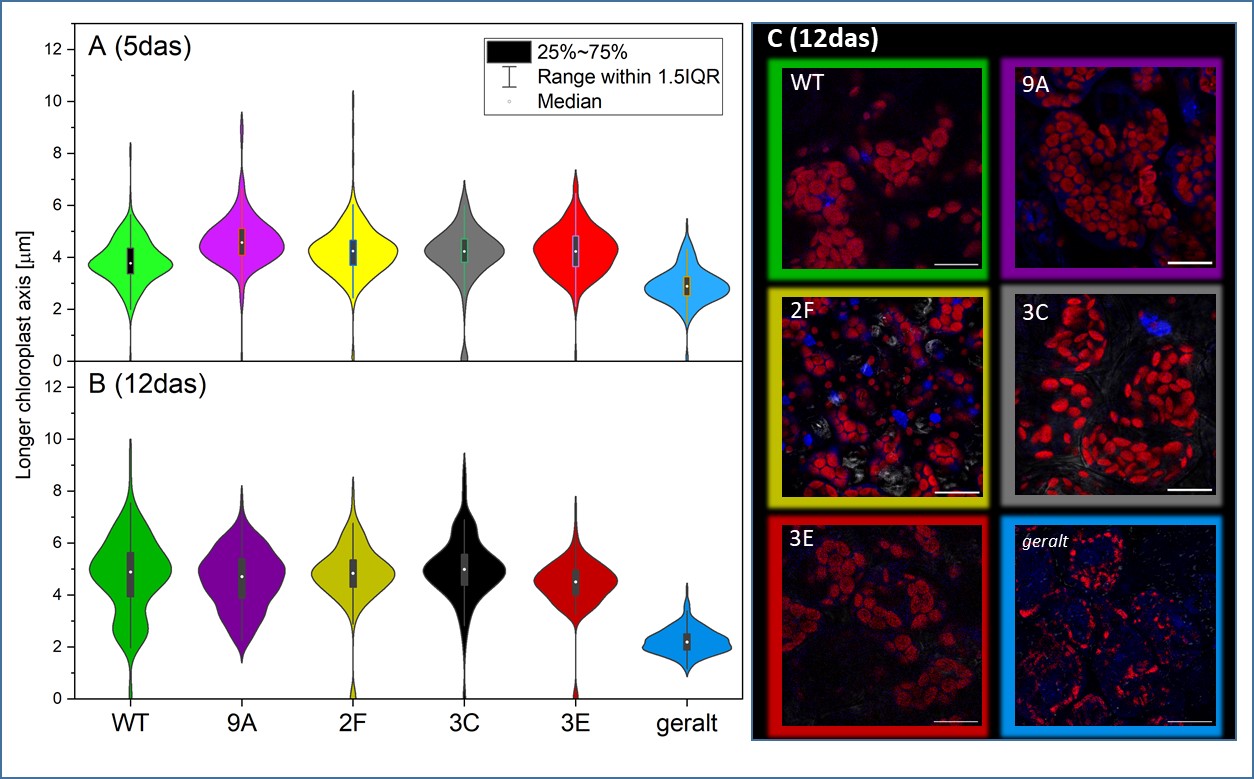

**Fig. S5. Comparison of chloroplast population size in 5 day-old and 12 day-old seedlings of WT, *geralt*, WT-OE_3E_ and complementant geralt-OE_9A/2F/3C_ lines.** Comparison of chloroplast population size (measured as length of chloroplast major axis, counted from SIM^2^ images) in (A) 5 day-old and (B) 12 day-old seedlings of WT, *geralt*, WT-OE_3E_ and complementant geralt-OE_9A/2F/3C_ lines. Violin plots (n = 170 to 220 individual chloroplasts) show the median and interquartile range (IQR) of the analyzed set. Right panel (C) shows examples of CLSM images (overlay of chlorophyll (red) and DAPI (blue) emission with transmitted light image) collected for 12 day-old seedlings of all mentioned plant lines. The color of the violin (A, B) corresponds to the color of frame (C).

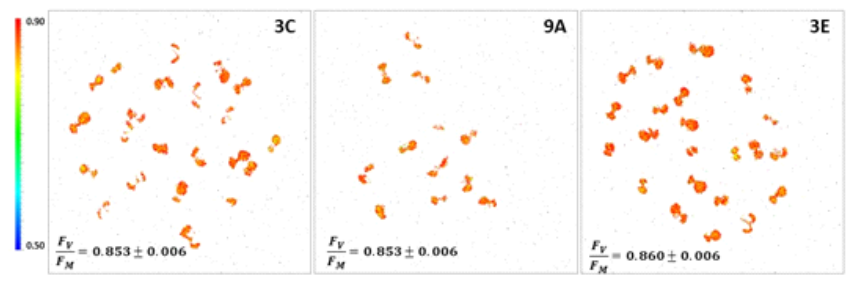

**Fig. S6. The maximum PSII quantum yield for plants overexpressing *GERALT*.** 5 day-old A) *geralt*_OE_3C; B) *geralt*_OE-9A, C) WT-OE_3E seedlings, were pre-adapted to darkness for 10 min prior to the fluorescence measurements. Mean F_V_/F_M_ values with standard deviations are respectively shown in the figures (calculated for 3 independent biological repetitions).

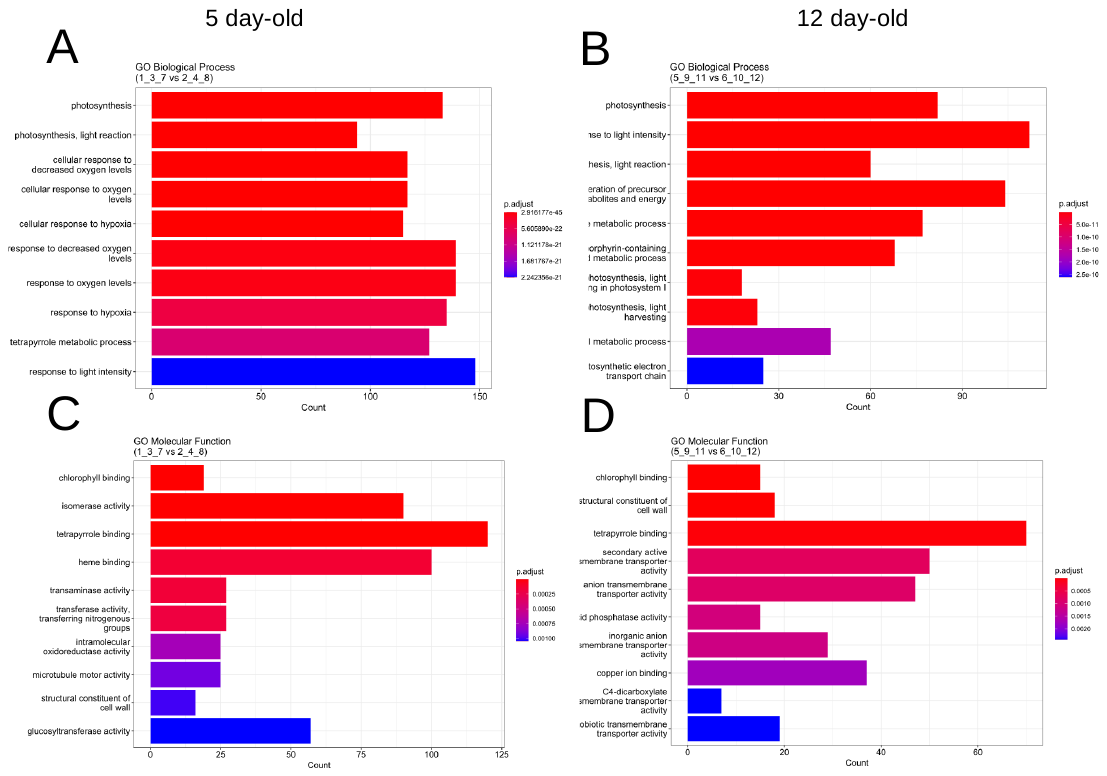

**Fig. S7**. GO analysis of 5 day-old (A, C) and 12 day-old (B, D) *geralt* and WT (control) seedlings. (A) GO Biological Process, ten gene categories and (C) Molecular function (MF), the list of ten activities present in proteins encoded by genes with significantly enrichment in GO molecular function categories out of 4 702 genes (p-value). (B) GO Biological Process, ten gene categories and (D) Molecular function (MF), the list of ten activities present in proteins encoded by genes with significantly enrichment in GO molecular function categories out of 2 663 genes (p-value).

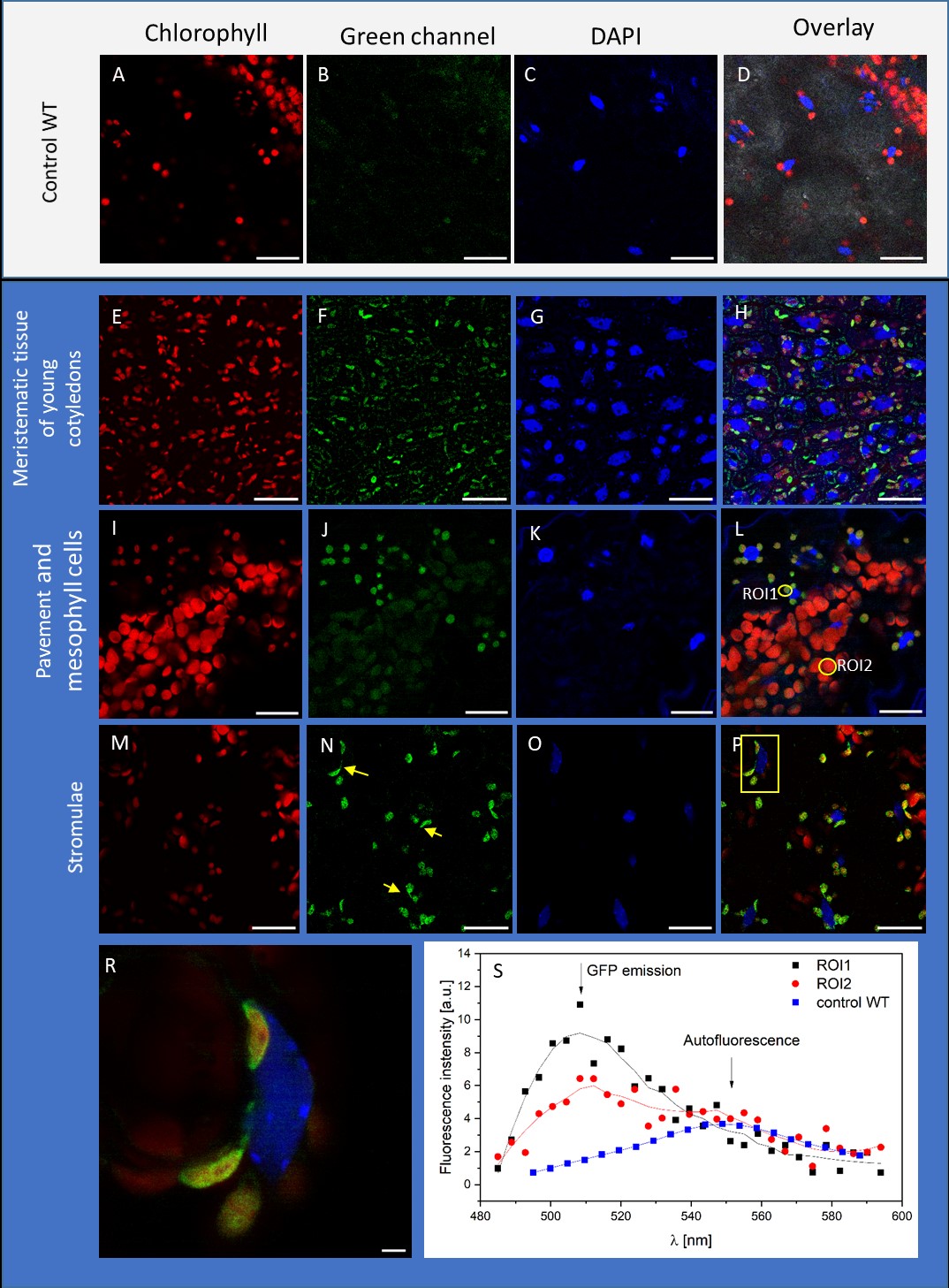

**Fig. S8.** **Localization of GERALT-GFP protein.** (A-D) Images of WT cotyledons of 5 day-old seedlings highlighted with white background. Images of WT-OE_3E_ plants highlighted in blue background. (E-H) Meristematic region of cotyledons of 3-day old WT-OE_3E_ seedlings. (I-L) – Typical localization of GERALT-GFP in pavement chloroplasts (see: ROI2 in L) with occasional detection over mesophyll chloroplasts (see: ROI1 in L) from 7 day-old seedlings. (M-P) – localization of GERALT-GFP in stromulae (marked with yellow arrow), formed between pavement cell chloroplasts and nuclei in 7 day-old seedlings. (R) Zoom-up at stromulae (all channels overlay, magnification of yellow box in P). (S) The graph of λ-stack based emission spectra recorded for ROIs marked on (L), in comparison with emission spectrum recorded for WT chloroplasts. Scale bar – 20 μm (A-P), inset R – 2 μm.

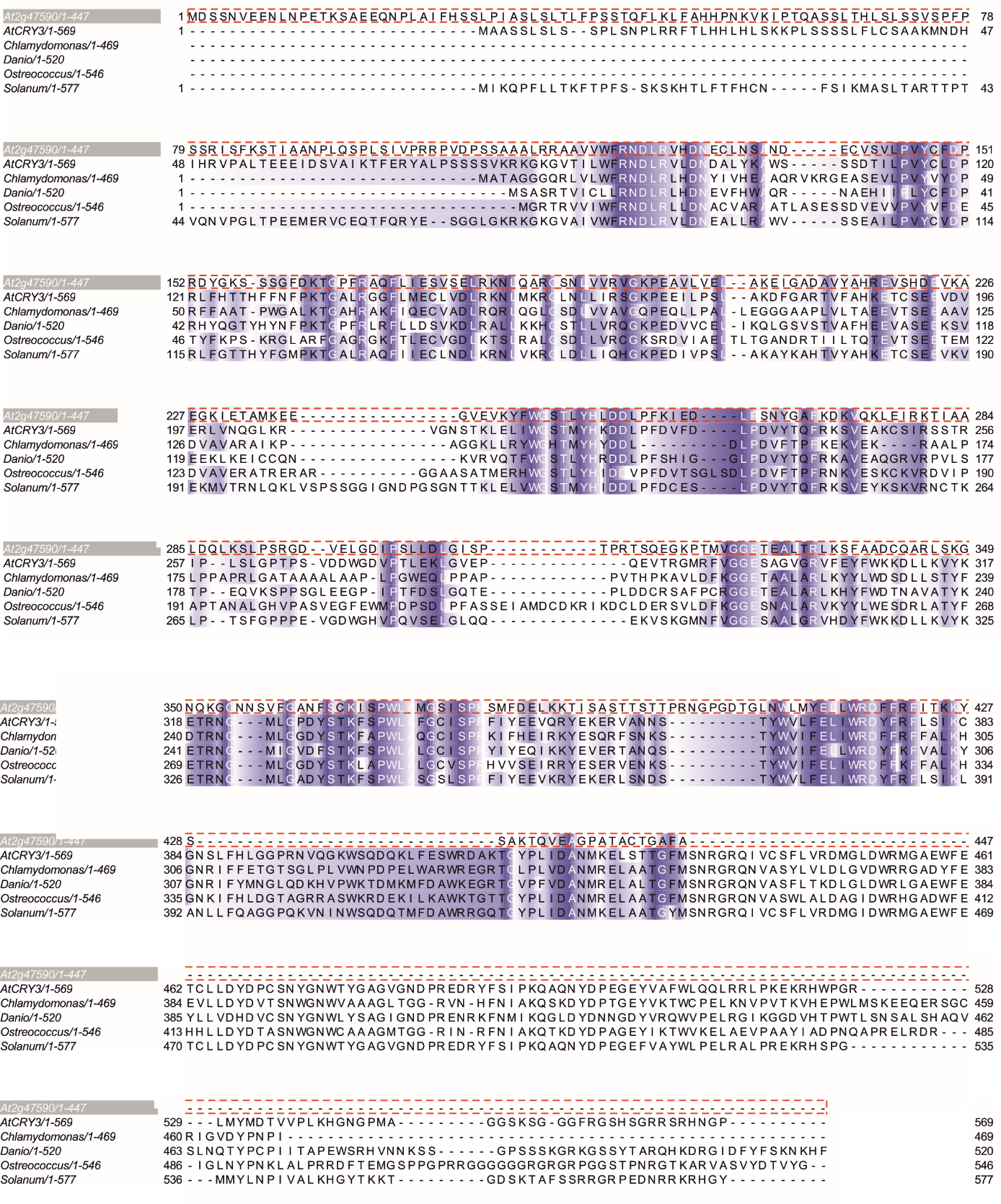

**Fig. S9.** **Alignment of GERALT (At2g47590) and selected CRY-DASH proteins**. Multiple sequence alignment was performed using ClustalX1.83 (http://www.clustal.org/) with default parameters and was visualized with Jalview (2.07).

Fig. S10.A – Proteins encoded by genes down-regulated in 5 day-old *geralt* seedlings

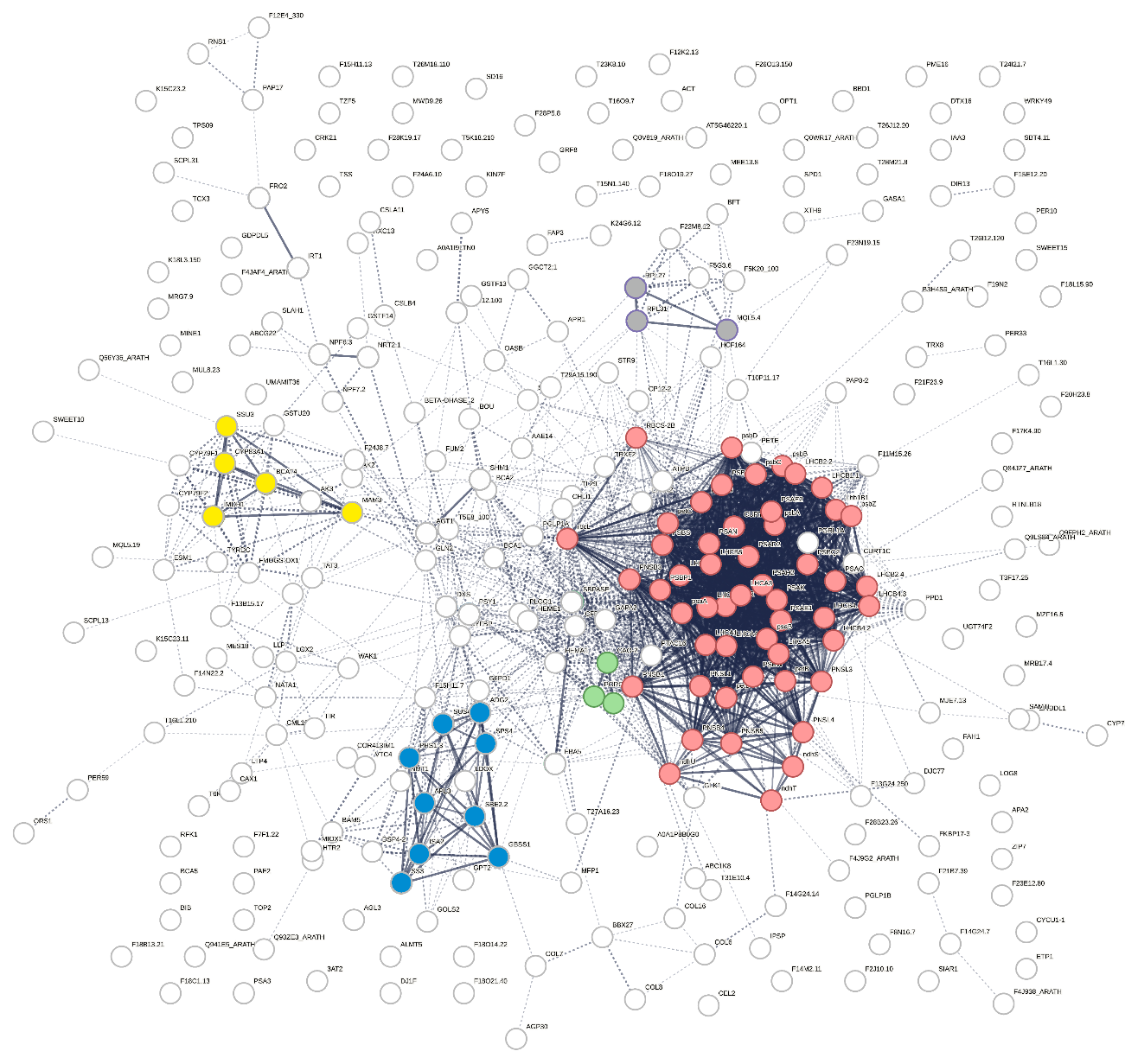

Clusters of proteins: 1) red – involved photosynthesis, 2) blue –involved starch and sucrose metabolism, 3) yellow – involved in glucosinolate biosynthesis, 3) green – involved in chlorophyll biosynthesis and 4) grey – being components of chloroplast ribosomes.

Fig. S10.B. Proteins encoded by genes up-regulated in 5 day-old *geralt* seedlings

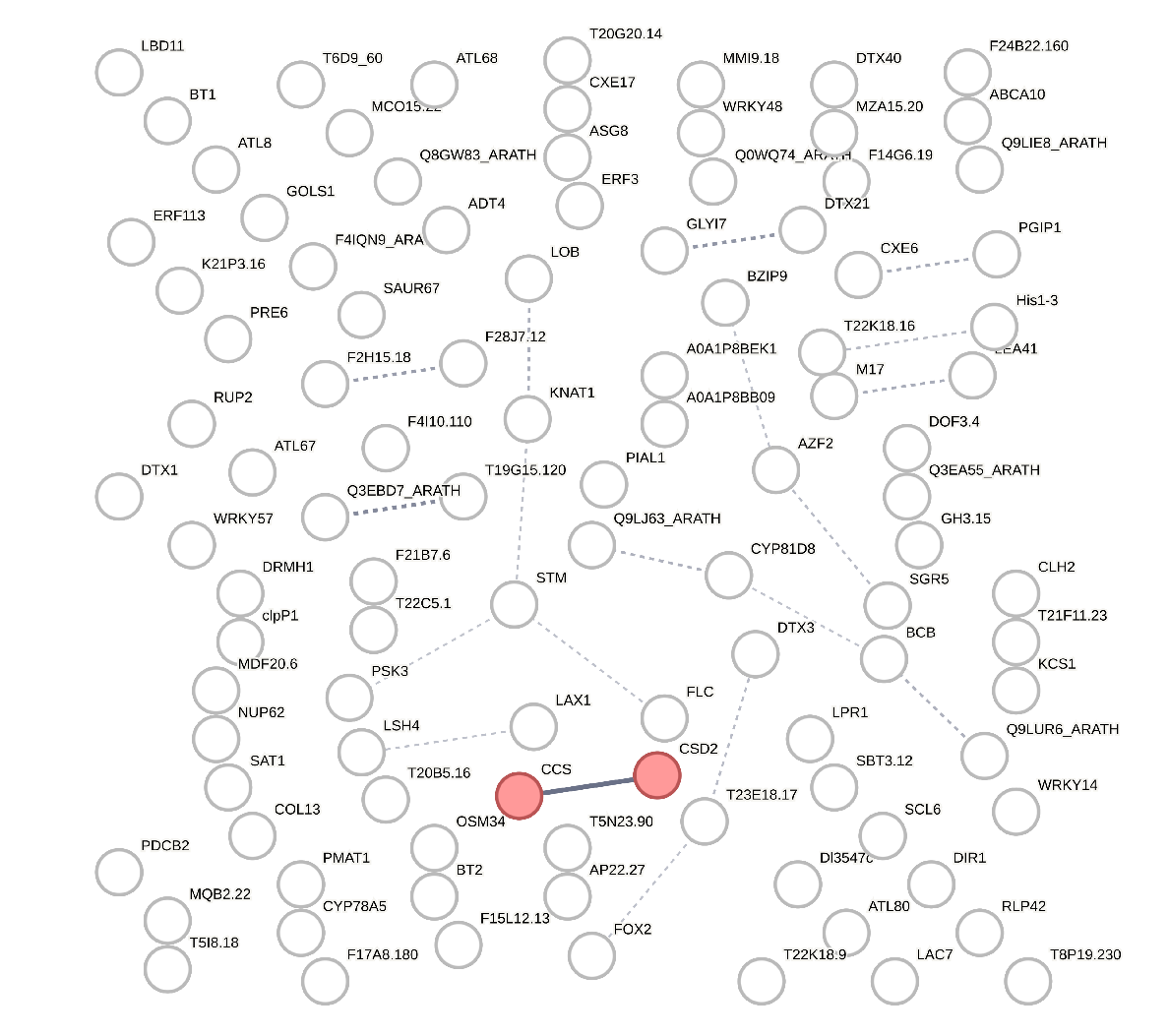

Fig. S10.C. Proteins encoded by genes down-regulated in 12 day-old *geralt* seedlings

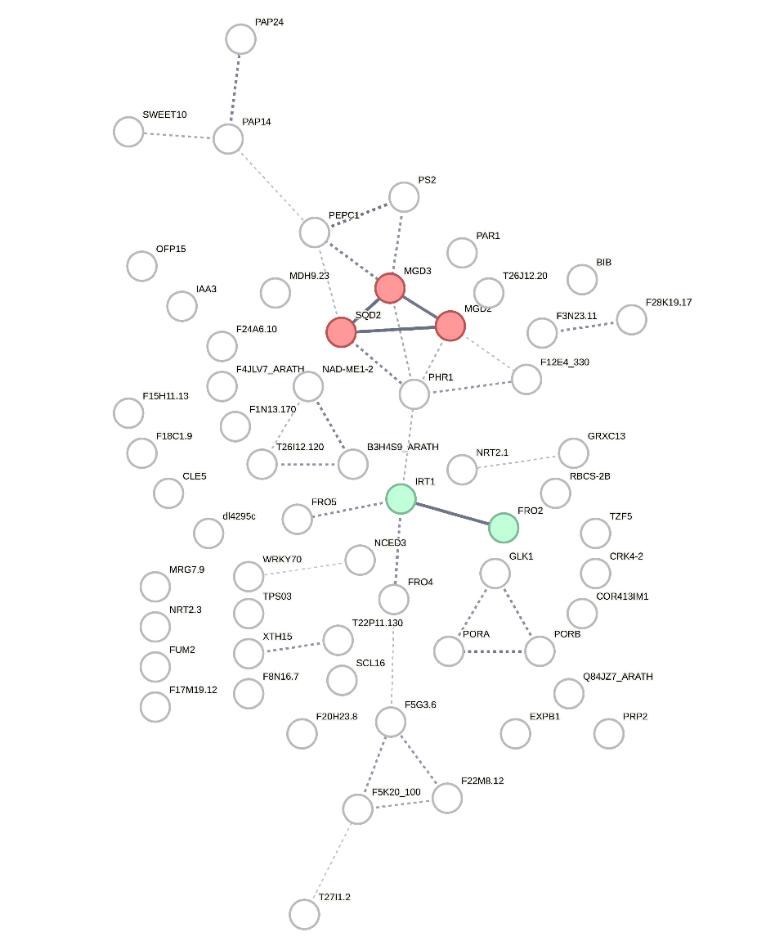

Red – lipid biosynthesis

Green – iron ion homeostasis

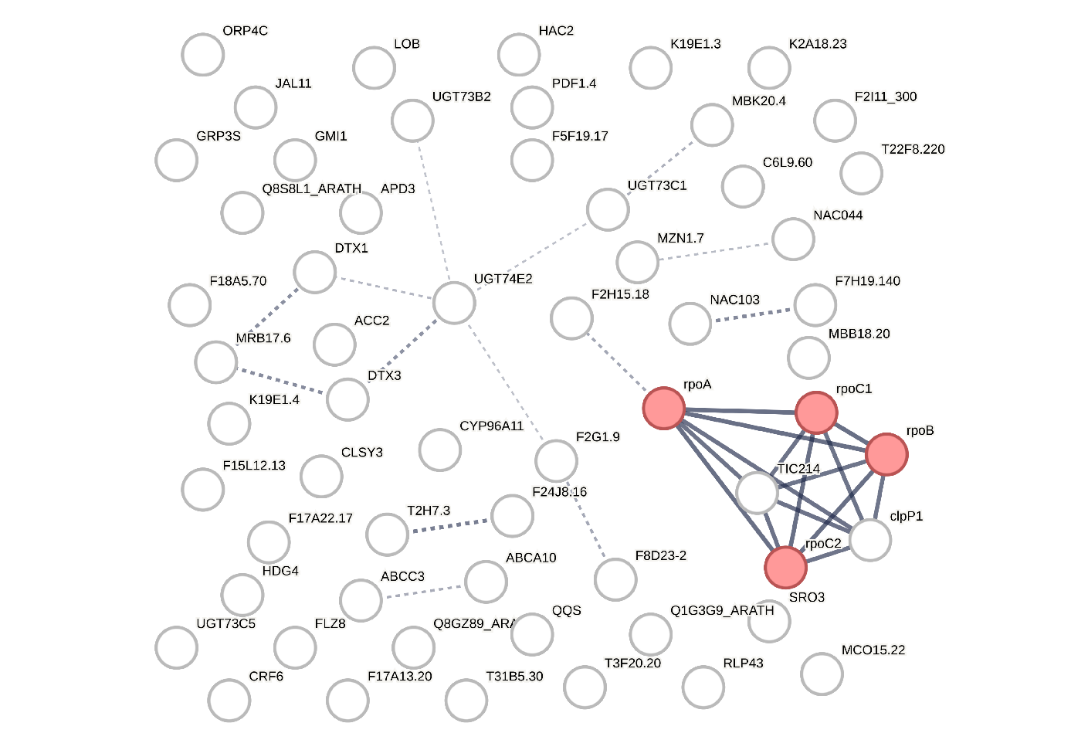
Fig. S10.D – 12 day–old up-regulated in *geralt*

Red - components of PEP polymerase (transcribed by NEP)

**Fig. S10**. **Maps of interactions between proteins encoded by genes down- or up-regulated in *geralt* seedlings visualised using STRING database (https://string-db.org)**. Line thickness indicates the strength of data suport (textmining, experiments, databases, co‑expression and protein co‑occurrence).

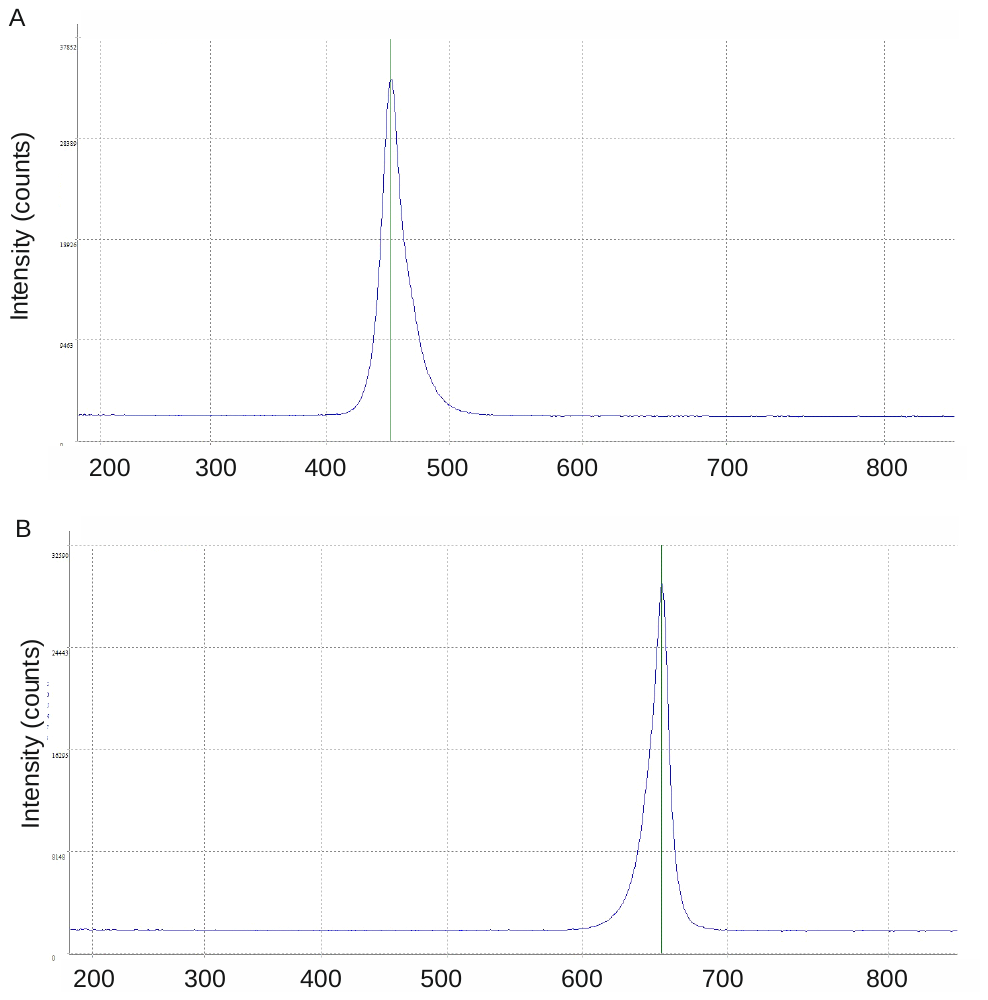

[nm]

**Fig. S11**. Spectra of (**A**) blue (470 nm) and (**B**) red (660 nm) light used in the experiments on gene expression.

**Supplemental Tables**

**Table S1.** List of DEGs identified in both 5 day-old and 12 day-old seedlings

| gene | fold change | | description |
| --- | --- | --- | --- |
| *Chloroplast encoded – up-regulated* | | | |
| *CLPP* | 3 | 5 | NEP dependent, chloroplastic ATP-dependent Clp protease |
| *Nuclear encoded – up-regulated* | | | |
| *AT5G55270* | 8 | 28 | putative mitochondrial, DUF 295 |
| *AT5G60250* | 5 | 19 | response to DNA damage, might act as an E3 ubiquitin-protein ligase, or as part of E3 complex |
| *AT1G17960* | 7 | 18 |  |
| *DTX3* | 4 | 10 |  |
| *ABCA10* | 4 | 7 | lipid metabolism |
| *Nuclear encoded – down-regulated* | | | |
| *PORB* | -8 | -6 | chlorophyll biosynthesis |
| *SHY2* | -4 | -6 |  |
| *AT5G18130* | -4 | -7 |  |
| *AT3G03870* | -3 | -7 |  |
| *TZF5* | -5 | -7 |  |
| *GPRI1* | -7 | -7 |  |
| *RbcS-2B* | -109 | -21 |  |
| *NRT2_1* | -20 | -23 | nitrate transporter |
| *IRT1* | -204 | -26 | iron transporter |
| *AT3G55240* | -348 | -50 | overexpression leads to PEL (Pseudo-Etiolation in Light) phenotype |
| *AT4G13575* | -920 | -461 |  |

**Table S2**. Selected photosynthesis-associated genes differentially expressed in *geralt* mutant

I. Chloroplast encoded genes

| **NEP dependent** | | | | | |
| --- | --- | --- | --- | --- | --- |
| gene | fold change | | gene | fold change | |
|  | 5 day-old | 12 day-old |  | 5 day-old | 12 day-old |
|  | *PEP subunits* | |  | *Translocon subunit* | |
| *RPOA** |  | 6 | *YCF1.2, TIC214_2* |  | 6 |
| *RPOC1** |  | 5 | *YCF1.1, TIC214_1* |  | 6 |
| *RPOC2** |  | 5 |  | *Proteasome subunit* | |
| *RPOB** |  | 5 | *CLPP1* | 3 | 5 |
| **PEP dependent** | | | | | |
| *PSI subunits* | | | *Cytochrome b6f complex* | | |
| *PSAB* | **-3** |  | *PETD* | **-10** |  |
| *PSAA* | **-3** |  | *PETB* | **-16** |  |
| *PSBD* | **-5** |  | *PSII subunits* | | |
| *PSBC* | **-6** |  | *PSBZ* | **-24** |  |
| *Calvin cycle enzyme* | | | *PSBA* | **-24** |  |
| *RBCL* | **-85** |  |  |  |  |

II. Nuclear encoded genes

| gene | fold change | | gene | fold change | |
| --- | --- | --- | --- | --- | --- |
|  | 5 day-old | 12 day-old |  | 5 day-old | 12 day-old |
| *PEP subunits* | | | *PSI assembly factors* | | |
| *SIGA (SIG1)** | **-3** |  | *Y3IP1* | **-3** |  |
| *PTAC16** | **-3** |  | *DEAP2* | **-2** |  |
| *PAP8_2** | **-3** |  | *PSA3* | **-3** |  |
| *PSI subunits* | | | *NADH-specific dehydrogenase* | | |
| *PNSB1* | **-2** |  | *ndhU* | **-3** |  |
| *LHCA5* | **-2** |  | *ndh5* | **-2** |  |
| *LFNR2* | **-3** |  | *ndhT* | **-3** |  |
| *PSAE2* | **-3** |  | *PSNB5* | **-3** |  |
| *PSAE1* | **-3** |  | *PNSL4* | **-4** |  |
| *PSAN* | **-3** |  | *PNSB4* | **-4** |  |
| *LHCA4* | **-4** |  | *PNSB3* | **-4** |  |
| *LHCA2* | **-4** |  | *PNSL3* | **-4** |  |
| *LHCA1* | **-4** |  | *PNSL1* | **-2** |  |
| *PsaK* | **-4** |  | *Cytochrome b6f complex* | | |
| *PSAO* | **-5** |  | *HCF164* | **-2** |  |
| *PSAD2* | **-5** |  | *PETC* | **-3** |  |
| *LHCB3* | **-5** |  | *Lipid synthesis* | | |
| *PSAH2* | **-6** |  | *MGD2* |  | **-6** |
|  |  |  | *AGAP1* | **-2** |  |
|  |  |  | *MGD3* |  | **-14** |

| gene | fold change | | gene | fold change | |
| --- | --- | --- | --- | --- | --- |
|  | 5 day-old | 12 day-old |  | 5 day-old | 12 day-old |
| *PSII subunits /maintanance* | | | *Chlorophyll biosynthesis* | | |
| *MPH2* | **-2** |  | *HEME1* | **-3** |  |
| *PSBR* | **-3** |  | *CHLI1* | **-2** |  |
| *PSBP1* | **-3** |  | *PORA* |  | **-37** |
| *LHCB4.1* | **-3** |  | *PORB* | **-8** | **-6** |
| *AT1G51400* | **-4** |  | *PORC* | **-5** |  |
| *PSBW* | **-4** |  | *CAO1* | **-4** |  |
| *PSBQ2* | **-4** |  | *CLA1* | **-3** |  |
| *LHCB6* | **-4** |  | *HEMA1* | **-3** |  |
| *PSBS* | **-4** |  | *Calvin cycle enzymes* | | |
| *LHCB5* | **-5** |  | *RBCS-2B* | **-109** | -**21** |
| *LHCB4.2* | **-5** |  | *SBPASE* | **-3** |  |
| *LHCB2.2* | **-6** |  | *Others* | | |
| *LHCB4.3* | **-6** |  | *ATPD* | **-3** |  |
| *LHCB1.1_1* | **-7** |  | *AT5G08050* | **-3** |  |
| *PSBP2* | **-8** |  | *AAE14* | **-4** |  |
| *PSBB* | **-19** |  | *PSY* | **-4** |  |
| *LHB1B1* | **-20** |  | *PETE1* | **-6** |  |
| *LHCB2.4* | **-33** |  |  |  |  |

**Table S3.** The selected transcription factors binding sites identified in the *GERALT* promoter based on PlantCare (https://bioinformatics.psb.ugent.be/webtools/plantcare/html/; Lescot et al. 2002) and AGRIS databases (https://agris-knowledgebase.org/AtcisDB/; Davuluri et al. 2003). The motifs which bind transcription factors involved in the light responses are marked with a yellow background.

| Binding site name | sequence | No | function |
| --- | --- | --- | --- |
| AtMYC2 BS in RD22 | cacatg | 5 | response to drought and abscicis acid |
| Bellringer/replumless/pennywise BS1 IN AG | aaattaaa | 2 | transcriptional repression of the floral homeotic gene AGAMOUS |
| ATB2/AtbZIP53/AtbZIP44/GBF5 BS in ProDH | actcat | 1 | response to hypoosmolarity |
| W-box promoter motif | ttgact | 3 | response to heat and salinity |
| ARF1 binding site motif | tgtctc | 1 | response to auxin |
| AuxRR-core | ggtccat | 1 | response to auxin |
| DPBF1&2 binding site motif | acacgag | 4 | response to abscisic acid |
| ABRE box | tacgtgtc | 1 | response to abscisic acid |
| MYB4 binding site motif | aac(t/a)aac | 3 | response to environmental stresses |
| RAV1-A binding site motif | caaca | 10 | growth regulation |
| LFY consensus binding site motif | cca(t/a)tg | 2 | floral merystem development |
| Hexamer promoter motif | ccgtcg | 1 | carbohydrate metabolism, plastid organization and translation |
| TGACG motif | tgacg | 4 | regulation of development and defence |
| GATA promoter motif [LRE] | (t/a)gataa/ agatag/ cacgtg | 10 | response to light |
| **Ibox promoter motif** | gataag | 2 | response to light |
| L1-box promoter motif | taaatg(c/t)a | 2 | L1-specific gene expression in the shoot system |
| T-box promoter motif | actttg | 3 | response to biotic and abiotic stress |
| SORLREP3 | tgtatatat | 1 | response to light |
| SORLIP2 | gggcc | 2 | response to light |
| SORLIP5 | gagtgag | 2 | response to light |
| 3-AF1 binding site | taagagaggaa | 1 | response to light, identified in *Solanum tuberosum* |
| AE-box | agaaacaa | 1 | response to light |
| Box 4 | attaat | 11 | response to light, identified in *Petroselinum crispum* |
| G Box | cacgtg | 4 | response to light, identified in *Pisum sativum* |
| I Box | tgataatgt | 1 | response to light, identified in *Solanum tuberosum* |
| TCT motif | tcttac | 3 | response to light |

**Table S4**. The list of primers used for cloning, genotyping and real-time PCR. Lower case - gateway sequences.

| Name | Primer sequence (5' - 3') | plasmid |
| --- | --- | --- |
| attB1Geralt_For | ggggacaagttgttacaaaaaagcaggcttcATGGATTCTTCGAATGTTGAAGAAAACC | pDONR221 |
| attB1Geralt_Rev_nostp | ggggaccactttgtacaagaaagctgggtcAGCAAAGGCACCGGTACAGG |  |
| Primers used for genotyping/testing the expression of *GERALT* CDS | | |
| Name | Primer sequence (5' - 3') | |
| Geralt_F | ATGGATTCTTCGAATGTTGAAGAAAACC | |
| Geralt_R | TTAAGCAAAGGCACCGGTAC | |
| geralt_genot_For | GAAATTGTGTGGGGAAAATCGACGAGAGC | |
| geralt_genot_Rev | GCTGAGCTGTATTTCTTGGTTATAAACCTGTTCC | |
| Lba1 | TGGTTCACGTAGTGGGCCATCG | |

| Primers used for real-time PCR | | |
| --- | --- | --- |
| Name | Primer sequence (5' - 3') | Tm [°C] |
| *qRT_GERALT_F1* | TTGGAGCTGACGCGGTTTAT | 56 |
| *qRT_GERALT_R1* | TGACTTCAACGCCTTCCTCC |  |
| *qRT_SAND For* | AACTCTATGCAGCATTTGATCCACT | 53 |
| *qRT_SAND Rev* | TGATTGCATATCTTTATCGCCATC |  |
| *qRT_PDF_For* | TAACGTGGCCAAAATGATGC | 53 |
| *qRT_PDF_For* | GTTCTCCACAACCGCTTGGT |  |
| *LPORA_For* | AGAGTCTAGTCTGTTCGGTGTTTCAC | 56 |
| *LPORA_Rev* | CTGATGGAGTTGAAGTCGCGATTGC |  |
| *LPORB_For* | CCGACCAAATCAAATCCGAACATGGA | 56 |
| *LPORB_Rev* | GTGGCTAGACCTAACCCAGACGAG |  |
| *LPORC_For* | AGATAAGCGTTGGAACCAACCATCTC | 56 |
| *LPORC_Rev* | AACTGTTTTGCCCATTCAATCCTGAC |  |
| *qRT_rpoC1_For* | TACATAGATTAGGCATACAGTCATTCCA | 50 |
| *qRT_rpoC1_Rev* | TGCGTCCTTCCACTAAAATAGGTT |  |
| *qRT_rpoA_For* | CAAGCCGACACAATAGGCAT | 50 |
| *qRT_rpoA_Rev* | AGCGCGTTGCGCGTTCCATA |  |
| *qRT_accD_For* | TTTATGGTTGGGATGAGCGT | 50 |
| *qRT_accD_Rev* | CCGATATGAAATTGCGAATGTCC |  |
| *qRT_RPS18_For* | GACGGGTGAATAGAGTGACTTT | 50 |
| *qRT_RPS18_Rev* | GGAGTCGACTCACTTCTTTCAA |  |
| *qRT_psbD_For* | GGATGACTGGTTACGGAGGG | 50 |
| *qRT_psbD_Rev* | GGTTGTACCTGTGAACCAACC |  |
| *qRT_psbB_For* | TCGTGCGACTTTGAAATCTGA | 50 |
| *qRT_psbB_Rev* | CAACCTCTTGGGCTGCTACG |  |
| *qRT_psaB_For* | GGACCCCACTACTCGTCGTA | 50 |
| *qRT_psaB_Rev* | ATTGCTAATTGCCCGAAATG |  |
| *qRT_rbcL_For* | TACCTGGTGTTCTGCCTGTG | 50 |
| *qRT_rbcL_Rev* | GCTACTCGGTTGGCTACGG |  |
| *qRT_atpB_For* | GAGCTCGTATGAGAGTTGGT | 50 |
| *qRT_atpB_Rev* | ACCCAATAAGGCGGATACCT |  |
| *qRT_clpP_For* | TGGGTTGACATATACAACCGACTTT | 50 |
| *qRT_clpP_Rev* | GCCTAAAAAAAATAATCTTTCTCGATAAA |  |
| *qRT_SAND_For* | AACTCTATGCAGCATTTGATCCACT | 53 |
| *qRT_SAND_Rev* | TGATTGCATATCTTTATCGCCATC |  |
| *qRT_UBC_For* | CTGCGACTCAGGGAATCTTCTAA | 53 |
| *qRT_UBC_Rev* | TTGTGCCATTGAATTGAACCC |  |
| *qRT_PDF_For* | TAACGTGGCCAAAATGATGC | 53 |
| *qRT_PDF_Rev* | GTTCTCCACAACCGCTTGGT |  |

**Table S5.** The list of vectors used for *Arabidopsis* transformation and the obtained transgenic lines

| Destination vector | Plant lines: name/description |
| --- | --- |
| pEarleyGate103 | *geralt*-OE_2F(3C)  *geralt*-/- complement lines expressing GERALT-GFP under 35S promoter obtained via transformation of *geralt* +/- plants, Basta resistance in plant |
| pH7FWG2 | *geralt*-OE_9A  *geralt*-/- complement line expressing GERALT-GFP under 35S promoter obtained via transformation of *geralt* +/- plants, hygromycine resistance in plant |
| pK7FWG2 | WT-OE_3E  WT plants overexpressing GERALT-GFP under 35S promoter, kanamycin restistance in plant |

Supplementary Video

**Supplementary Video_S1. Supplementary file wt_3D_Z-stack -** **Collection of Z-stack images (combined as a movie) of mesophyll chloroplasts of WT plants**. Images obtained by CLSM technique, are overlay of chlorophyll (red) and DAPI (blue) emission. Scale bar is 20 μm.

**Supplementary Video_S2. Supplementary file Geralt_3D_Z-stack - Collection of Z-stack images (combined as a movie) of mesophyll chloroplasts of *geralt* plants**. Images obtained by CLSM technique, are overlay of chlorophyll (red) and DAPI (blue) emission. Scale bar is 20 μm.

**Supplementary Video_S3. WT_3D_Z-stack** - **Z-stack image (combined as a movie) of selected mesophyll chloroplasts of WT plants**. Images obtained by SIM^2^ technique (subfiles: 1_3D to 3_3D), are overlay of chlorophyll (red) and DAPI (blue) emissions.

**Supplementary Video_S4. geralt_3D_Z-stack** - **Z-stack image (combined as a movie) of selected mesophyll chloroplasts of *geralt* plants**. Images obtained by SIM^2^ technique (subfiles: 1_3D to 4_3D), are overlay of chlorophyll (red) and DAPI (blue) emissions.

**Supplementary Video_S3. Geralt_GFP_3D_Z-stack** - **Z-stack image (combined as a movie) of selected pavement chloroplast of WT-OE_3E_ plants**. Images obtained by SIM^2^ technique (subfile: Geralt_GF_Z-stack1), are overlay of chlorophyll (red), GFP (green) and DAPI (blue) emissions. Additionally, free hand rotations of 3D projections, based on that Z-stacks are presented for all three channels overlay (subfile: Geralt_GFP_Z-stack_movie1) and GFP channel only (subfile: Geralt_GFP_Z-stack_Gfp_movie). 3D projection was constructed with 3D viewer plugin of Fiji software (Schmid *et al.*, 2010). No PSF correction was applied to the Z-stacks.

**Supplementary data S1**. Amino acid sequences of proteins used to construct a phylogenetic tree.

>NP_182281.1 photolyase/blue-light receptor 2 [Arabidopsis thaliana]

MDSSNVEENLNPETKSAEEQNPLAIFHSSLPIASLSLTLFPSSTQFLKLFAHHPNKVKIPTQASSLTHLSLSSVSPFPSSRISFKSTIAANPLQSPLSIVPRRPVDPSSAAALRRAAVVWFRNDLRVHDNECLNSANDECVSVLPVYCFDPRDYGKSSSGFDKTGPFRAQFLIESVSELRKNLQARGSNLVVRVGKPEAVLVELAKEIGADAVYAHREVSHDEVKAEGKIETAMKEEGVEVKYFWGSTLYHLDDLPFKIEDLPSNYGAFKDKVQKLEIRKTIAALDQLKSLPSRGDVELGDIPSLLDLGISPTPRTSQEGKPTMVGGETEALTRLKSFAADCQARLSKGNQKGGNNSVFGANFSCKISPWLAMGSISPRSMFDELKKTISASTTSTTPRNGPGDTGLNWLMYELLWRDFFRFITKKYSSAKTQVEAGPATACTGAFA

>XP_006397953.1 blue-light photoreceptor PHR2 [Eutrema salsugineum]

MDSNLVEENPNPETKSTEEPLAIIPSSLPIASLSLSFFPNSTTHFLRFSHLFSHHPNKVKIPTQASYLTHLSLSSVSPSPSTRISFKSTVSANPLQSPLSIVPRRPVDPSSAAALRRSAVVWFRNDLRVHDNECLNSANDECVSVLPVYCFDPRDYGKSSSGFDKTGPFRAQFLIDSVTELRKNLEARGSNLVVRVGKPESVLVELAKEIGADAIYAHREVSHDEVKAEGKIETAMKEEGVEVKYFWGSTLYHLDDLPFKIEDLPSNYGAFKDKVQKLEIRKTIAALDQLKSLPSRGDVQLGDIPSLLDLGISPSARTSQEGKPTMAGGETEALNRLKSFAADCQARLNKKNQKGDNNSVFGANFSCKISPWLAMGSISPRSMFDELKRSVSASTTPKTPRNGPGDTGLNWLMYELLWRDFFRFITKKYSSAKTQVEAGPATACTGAFA

>VVB03633.1 unnamed protein product [Arabis nemorensis]

MDSNLEENPNPETKSAEEQQNPLAIISSSLPIASLSLSFFPTCTQFLINPKFSRLFTHHPNKVKIPTQASYLTHLSLSSVSSSPLPSRISFKSTIAANPLQSPLSIVPRRPVDPSSAAALRRAAVVWFRNDLRVHDNECLNSANDECVSVLPVYCFDPRDYGKSSSGFDKTGPFRAQFLIESVSELRKNLQARGSNLVVRVGKPESVLVELAKEIGADAVYAHREVSHDEVKAEGKIEKAMKEEGVEVKYFWGSTLYHLDDLPFKIEELPSNYGAFKDKVQKLEIRKTIAALDQLKSLPSRGDVELGDIPSLLDLGVSPSARTSQEGKPTMVGGETEALNRLKSFAADCQARLNKGNQKGGNNSVFGANFSCKISPWLAMGSISPRSMFDELKKTVSASTTPKGPRNGPGDTGLNWLMYELLWRDFFRFITKKYSSAKTQVEAGPATACTGAFA

>XP_023639395.1 blue-light photoreceptor PHR2 [Capsella rubella]

MDSKNVEEKLNEETKSAEEHNPVAAIIPSSLPIASLSLITFFPTSTQFLRLFSHHPNKVRIPTQASSLTHLSLSSVSPPPPSSISFKSTIAANPLQSPLSIVPRRPVDPSSAAALRRAAVVWFRNDLRVHDNECLNSANDECVSVLPVYCFDPRDYGKSSSGFDKTGPFRAQFLIESVSELRKNLQARGSNLVVRVGKPEAVLLELAKEIGADAVYAHREVSHDEVKAEGKIETALKEEGVEVSFFWGSTLYHLDDLPFNIEDLPSNYGAFKDKVQKLEIRKTIAALDQLKSLPSRGDVELGDIPSLLDLGVCPTARTSQEGKPTMVGGETEALTRLKSFAAECQARLSKGNQKGGNNSVFGANFSCKISPWLAMGSISPRSMFDELKKTISASTTTTTPRNGPGDTGLNWLMYELLWRDFFRFITKKYSSVKTQVEAGPATACTGAFA

>XP_013747552.2 blue-light photoreceptor PHR2-like [Brassica napus]

MESKVVVDAETKSTEEEEAHAIVTCPCLPIASLSLSFSPLSTTTTQFRFSHLFAHHPNKVKIPTQASYLTHLSLSSLSSPSPPLPSRISFKSTFSANPLHSPLSIVPRRPLDPSSAAALRRSAVVWFRNDLRVHDNECLTSANDECLSVLPVYCFDPRDYGKSSSGFDKTGPFRAQFLIESVSELRKNLEARGSNLVVRVGKPEAVLVELAKEIGADAVYAHREVSHDEVKAEGKIESAMKEEGVEVKFFWGSTLYHLDDLPFKVEDLPSNYGAFKDKVQKLEIRKTIAALDQLKSLPSRGDVQLGDIPSLLDLGINPSARTSQEGKPTMVGGETEALTRLKSFAADCQARLTKGGNQKGGNNSVFGANFSCKISPWLAMGSISPRSMFDELKKTISASTTPSTPRNGPGDTGLNWLMYELLWRDFFRFITKKYSSAKTQVEEAGPATACTGALA

>XP_018440307.2 blue-light photoreceptor PHR2 [Raphanus sativus]

MESNVVVVENQNAERKSTEEEEAVATIVTSSLPIASLSLSLSTQFRFSQLFAHHPNKVKIPTQASYLTHLSLSSLSPSPPRITSSFSANPLHSPLSIVPRRPFDPSSAAALRRSALVWFRNDLRVHDNECLNSANDESLSLLPVYCFDPRDYGKSSSGFDKTGPFRAQFLIESVSELRKNLEARGSNLVVRVGKPETVLVELAKEIGADAIYAHREVSHDEVKAEGKIESAMKEEGVEVKYFWGSTLYHLDDLPFKIQDLPSNYGAFKDKVHKLEIRKTIAALDQLKSFPSRGDVQLGDIPSLLDLGINPTPRTSQEGKPTMVGGETEALTRLKSFAADCQARLTKGGNHKGGNNSVFGANFSCKISPWLAMGSISPRSMFDELKKTISAASTPRNGPGDTGLNWLMYELLWRDFFRFITKKYSSAKTQVEAGPATACTGALA

>XP_009142517.1 blue-light photoreceptor PHR2 [Brassica rapa]

MESKVVVDGETKSTEEEEAHAIVTCPCLPIASLSLSFSPLSTTTTQFRFAHLFAHHPNKVKIPTQASYLTHLSLSSLSSPSPPLPSRISFKSTFSANPLHSPLSIVPRRPLDPSSAAALRRSAVVWFRNDLRVHDNECLNSANDECLSVLPVYCFDPRDYGKSSSGFDKTGPFRAQFLIESVSELRKNLEARGSNLVVRVGKPEAVLVELAKEIGADAVYAHREVSHDEVKAEGKIESAMKEEGVEVKFFWGSTLYHLDDLPFKVEDLPSNYGAFKDKVEKLEIRKTIAALDQLKSLPSRGDVQLGDIPSLLDLGINPSARTSQEGKPTMVGGETEALTRLKSFAADCQARLAKGGNQKGGNNSVFGANFSCKISPWLAMGSISPRSMFDELKKTISASTTPSAPRNGPGDTGLNWLMYELLWRDFFRFITKKYSSAKTQVEEPGPATACTGALA

>KAJ0258099.1 Blue-light photoreceptor PHR2 [Hirschfeldia incana]

MESNVVVKNQNAETKSTEEEEESLAIAIVTSSLPIASLSLSFSPLTTTQFRFSDLFAHHPNKVKIPTQASYLTHLSLSSLPPSPRRIFKSTFKSANPLHSPLSIVPRRPVDPSSSAALRRSAVVWFRNDLRVHDNECLNSANDECLSLLPVYCFDPRDYGKSSSGFDKTGPFRAQFLIESVSELRKNLEARGSNLVVRVGKPESVLVELAKEIGADAVYAHREVSHDEVKAEGKIEKAMKEEGVEVNYFWGSTLYHLDDLPFKIEELPSNYGAFKEKVQKLEIRKTIAALDQLKSLPSRGDVQLGDIPSLLDLGINPSARTSQEGKPTMVGGETEALTRLKSFAADCQARLSKGGNQKGGNNSVFGANFSCKISPWLAMGSISPRSMFDELKKTISTPSTPRNGPGDTGLNWLMYELLWRDFFRFITKKYSSAKTQVEAGPATACTGALA

>XP_023880694.1 blue-light photoreceptor PHR2 [Quercus suber]

MDPNTQALENPETKSSEEQNPHAIVASKVTLPFATLSLSISLPKILQTPSLETKIQSLFSRHPSKVKVPTQASSLAHLSLSATAPVPSKLSFKSTISANPLQNPLSLGPRRPLDPNNGAGIRRAAIVWFRNDLRVNDNECLNTANNEAMSVLPVYCFDPRDYGKSSSGFDKTGPYRASFLIESVSDLRKNLQAKGSDLVVRIGQPETVLLELAKAIGADAVYAHREVSHDEVKGEDKIESAMKEENVEVKYFWGSTLYHLDDLPFKLEDMPSNYGGFKEKVQGLEVRKTIEALDQLKNLPSRGDVEAGDIPSLADLGLNPSATMSQANASMVGGETEALQRLQKFAAECQAQPHKGTKDGSNNSIYGANFSCKVSPWLAMGCISPRSMFDELKKTAARTISASSNRNDGGSPDTGMNWLMFELMWRDFFRFITKKYSSAKRQIEAPVTTCTGALA

>XP_021906987.1 blue-light photoreceptor PHR2 [Carica papaya]

MVSSLQITENPEPKSADEENPVAIIASPPPFLPVASISLSLSTILSNPTAFSFLQPKISALFSHHPNKVKVPTQASSLSHLSLSSSSSNFSPTRISFKSTIAANPLQSPLSLGPRRPLDPSNGAALRRASIVWFRNDLRVHDNECLHTANEESMSVLPVYCFDPREYGKSSSGFDKTGPFRAQFLIESVSDLRKNLQARGSDLVVRFGKPETVLVELAKAIGADAVYSQREVSHDEVKAEEKIEAAMKEEGMEVKYLWGSTLYHIDDLPFKLEDMPTNYGGFREKVKGLEVRKTIDALDQLKGLPSRGDVEPGEIPSLMDLGLNPSATMTQGGKPVTNASMVGGETEALQRLKKFAAECQAQPNKGNRDGGNDSIYGANFSCKISPWLAMGCISPRSMFDELKKTVTRNVYAESKKNSGGNGSGDTGTNWLMFELLWRDFFRFITKKYSCAKRQTEAAPATACTGALA

>XP_042992757.1 blue-light photoreceptor PHR2 [Carya illinoinensis]

MDPNAQIHENPEMKSSEEQNPLAIVPSKDQSNSAPFATASLSLSLNTILPTPFSLQPKIYSLFSHQPSKVKVPTQASSLSHLSLTATINSPSPSRLSFKSTVSANPLQNPLSLGPRRPLDPSNGAGLRRAAIVWFRNDLRVHDNECLNTANNESMSVLPVYCFDPRDYGKSSSGFDKTGPYRATFLIESVSDLRKNLQAKGSDLVVRIGKPETVLVELAKEVGADAVYAHREVSHDEVKAEGRIEAAMKEENVEVKYFWGSTLYHLDDLPFKLEDMPANYGGFREKVQGLEVRKTIEALDQVKGLPARGDVEPGDIPSLVDLGLNPSATMAQDGKPAANASMVGGETEALQRLKKFAAECQAQPHKGKSDGSSNSIYGANFSCKISPWLAMGCLSPRSMFDGLKKTATRTISASSNRNDSGSPDTGNNWLMFELLWRDFFRFITKKYSSKQLEAAPVTACTGALA

>XP_030946281.1 blue-light photoreceptor PHR2 [Quercus lobata]

MDPNTQALENPETKSSEEQNQHAIVASKDALPFATLSLSISLPKILQTPSLETKIQSLFSRHPSKVKVPTQASSLAHLSLSTTAPVPSKLSFKSTISANPLQNPLSLGPRRPLDPNNGAGIRRAAIVWFRNDLRVNDNECLNTANNEAMSVLPVYCFDPRDYGKSSSGFDKTGPYRASFLIESVSDLRKNLQAKGSDLVVRIGQPETVLLELAKAIGADAVYAHREVSHDEVKGEDKIESAMKEENVEVKYFWGSTLYHLDDLPFKLEDMPSNYGGFKEKVQGLEVRKTIEALDQLKNLPSRGDVEAGDIPSLADLGLNPSATMSQANASMVGGETEALQRLQKFAAECQAQPHKGTKDGSNDSIYGANFSCKISPWLAMGCVSPRSMFDELKKTAARTISASSNRNDGGSPDTGMNWLMFELMWRDFFRFITKKYSSANRQNEAPVTACTGALA

>XP_018858305.2 blue-light photoreceptor PHR2-like [Juglans regia]

MDPNSQIHENPEMKSSEEQNPLAIVPSKDQTTSAPFATASLSFSLNTILPTPFSLQHKIYSLFSHQPSKVKVPTQASSLSHLSLTATTNSPSPSRLSFKSTVSANPLQNPLSLGPRRPFDPSNGAGLRRAAIVWFRNDLRVHDNECLNTANNESMSVLPVYCFDPRDYGKSSSGFDKTGPYRATFLIESVSDLRKNLQAKGSDLMVRIGKPETVLVELAKEVGADAVYAHREVSHDEVKAEGRIEAAMKEENVEVKYFWGSTLYHLDDLPFKLEDMPANYGGFREKVQGLEVRKTIEALDKVKGLPARGDVEPGDIPSLVDLGLNPSATMAQDGKPAANASMVGGETEALQRLNKFAVECQAQPHKGKSDGSSDSIYGANFSCKISPWLAMGCLSPRSMFDELKKTATRTISASSNRNDGGSPDTGNNWLMFELLWRDFFRFITKKYSSKQLEAAPVTACTGALA

>XP_059438818.1 blue-light photoreceptor PHR2 [Corylus avellana]

MDPNTQNLENPEAKSSEEQNPLAIVPCKDQTTALPFATASLSLSLSTILPTPFTLQPKIQSLFSHQPSKVKVPTQASSLAHLSLSATSLSPTPSKLSFKSTISANPLQNPLSLGPRRPLDPSNGAGIRRAAIVWFRNDLRVHDNECLNTANNEAMSVLPVYCFDPRDYGKSSSGFDKTGPYRASFLIESVSDLRKNLQAKGSDLVVRVGKPETVLAELAKAVGADAVYAHREVSHDEVKAEEKIEAAMKDESVEIKYFWGSTLYHLDDLPFKLEDMPANYGGFREKVKSLDVRKTIEALDQVKGMPARGDVEPGDIPSLVDLGLNPSATMGQDGKPAANASMVGGETEALQRLQKFAAECQAQPPKGTKDGSNDSIYGANFSCKISPWLAMGCLSPRSMFDELKKTATRTISASSNRDDGGSPDTGMNWLMFELMWRDFFRFITKKYSSKQLEAAPATACTGALA

>XP_021805119.1 blue-light photoreceptor PHR2 [Prunus avium]

MDPNLQVSENPESKSSEEQNALAIVPSSSLPPFATASLSLSLSTILPTHFFQQPKVSTLFSSQPNKVKVPTQASSLAHLSLSTANVTPPKLSFKSTISANPLQNPLTLGPRRPLDPSNGAAIRRASIVWFRNDLRVHDNECLNSASNESMSVLPVYCFDPRDYGKSSSGFDKTGPYRASFLVESVSDLRKNLQARGSNLVVRIGKPETVLVELAKAIGADAIYAHREVSHDEVKAEEKIEAAMKEENVEVKYFWGSTLYHMDDLPFKLEEMPTNYGGFREKVKGLEVRKTIEALEQMKGLPSRGDVEPGDIPSLMDLGLNPSATMSQDGRPAANASMVGGEAEALERLKKFAAECQAQPPKGGKDGSHDSIYGANFSCKISPWLAMGCLSPRSMFDELKKTANRTISASSNRDDGGSGMNWLMFELLWRDFFRFVTKKYSSAKKQLDAAPATACTGALA

>XP_006479977.1 blue-light photoreceptor PHR2 isoform X2 [Citrus sinensis]

MDPNKQNLENPENHSNEEQNPLATIPSQSPFATLSLSFSLPQVLPANTFFIQPKISTLFSHQPNKVKVPTQASTLTHISLSASSTLSPSKISFKSTLSANPLQSPLSLGPHRPLDPNNGAAIRRASIVWFRNDLRVHDNESLNTANNESVSVLPVYCFDPRDYGKSSSGFDKTGPYRASFLIESVSDLRKNLQARGSDLVVRVGKPETVLVELAKAIGADAVYAHREVSHDEVKSEEKIEAAMKDEGIEVKYFWGSTLYHLDDLPFKLGEMPTNYGGFREKVKGVEIRKTIEALDQLKGLPSRGDVEPGDIPSLLDLGLSQSAAMSQGGKPAANSMKGGETEALQRLKKFAAEYQAQPPKGNKDGNHDSIYGANFSCKISPWLAMGCLSPRSMFDELKKTATSISAASKRNDGESGSSGAGSNWLMFELLWRDFFRFITKKYSSAKKVVEAVPATACTGALA

>XP_007199831.1 blue-light photoreceptor PHR2 [Prunus persica]

MDPNLQVSENPESKSSEEPNSLAIVPSSSLPPFATASLSLSLSTILPTHFFQQPKVSTLFSSQPNKVKVPTQASSLAHLSLSTANVTPPKLSFKSTISANPLQNPLSLGPRRPLDPSNGAAIRRASIVWFRNDLRVHDNECLNSANNESMSVLPVYCFDPRDYGKSSSGFDKTGPYRASFLVESVSDLRKNLQARGSDLVVRIGKPETVLVELAKAIGADAIYAHREVSHDEVKAEDKIEAAMKEENVEVKYFWGSTLYHMDDLPFKLEEMPTNYGGFREKVKGLEVRKTIEALEQMKGLPSRGDVEPGDIPSLMDLGLNPSATMSQDGRPAANASMVGGEAEALERLKKFAAECQAQPPKSGKDGSHDSIYGANFSCKISPWLAMGCLSPRSMFDELKKTANRTISASSKRDDGGSGMNWLMFELLWRDFFRFVTKKYSSAKKQLDAAPATACTGALA

>XP_034228312.1 blue-light photoreceptor PHR2 [Prunus dulcis]

MAPNLQVSENPESKSSEEPNSLAIVPSSSLPPFATASLSLSLSTILPTHFFQQPKVSTLFSSQPNKVKVPTQASSLAHLSLSTANVTPPKLSFKSTISANPLQNPLSLGPRRPLDPSNGAAIRRASIVWFRNDLRVHDNECLNSANNESMSVLPVYCFDPRDYGKSSSGFDKTGPYRASFLVESVSDLRKNLQARGSDLVVRIGKPETVLVELAKAIGADAIYAHREVSHDEVKAEEKIEAAMKEENVEFKYFWGSTLYHMDDLPFKLEEMPTNYGGFREKVKGLEVRKTIEALEQMKGLPSRGDVEPGDIPSLMDLGLNPSATMSQDGRPAANASMVGGEAEALERLKKFAAECQAQPPKSGKDGSHDSIYGANFSCKISPWLAMGCLSPRSMFDELKKTANRTISASSKRDDGGSGMNWLMFELLWRDFFRFVTKKYSSAKKQLDAAPATACTGALA

>KAJ4727695.1 Blue-light photoreceptor PHR2 [Melia azedarach]

MDPNHQNLENPEDSSNEEQNPLAIIPSQSPFATLSLSFSLSKVLPTNTFFVQPKISTLFSHQPNKVKVPSQASSLSHLSLSASSATIAPSKLSFKSTISANPLQSPLDLGPHRPLDPNNGAAIRRASIVWFRNDLRVHDNECLNTASNESMSVLPVYCFDPRDYGKSSSGFDKTGPYRASFLIESVSDLRKNLQARGSDLVVRVGKPETVLVEMAKAIGADAVYAHREVSHDEVKSEEKIEAAMKEEGVEVKYFWGSTLYHIDDLPFKLEEMPTNYGGFREKVQGLEIRKTIEALDQLKGMPSRGDVEPGDIPTLLDLGLSSSATMSQDGKPATNTMVGGETEALQRLKKFAAECQAQPHKGTKDGSSDNIYGANFSCKISPWLAMGCLSPRSMFDELKKTATSISAAAKRNDGGSDSSATGSNWLMFELLWRDFFRFITKKYSSAKKLVEAAPATACTGALA

>PON55309.1 DNA photolyase class 1, 8-HDF type [Trema orientale]

MDPDSKMEKNPETQPLEEQNSLDIIVASSPFATASLSLSLSTIIPTHFFQQPKISTLFSQPNKVKVPTQASSLTHLSLSSASVTPPPKLSFKSTISANPLQNPLSLGPRRPMDPSNGAAVRRASIVWFRNDLRVHDNECLNSAHNESMSVLPVYCFDPRDYGKSSSGFDKTGPYRATFLVESVTDLRKNLQARGSDLVVRVGKPETVLAELAKAIGADAVYAHREVSHDEVKAEGRIEAAMKEENIEVKYFWGSTLYHVDDLPFKLEDMPSNYGGFREKVQGLDVRKTIEALDQMKGLPSRGDVEPGDIPSLMDLGINPSATMAQDGKPAANVSLVGGETEALERLKKFAAECQAQPHKGSKDGSHDSIYGANFSCKISPWLAMGCLSPRSMFDELKKTTTRTISASSSQNDGGSGMNWLLFELLWRDFFRFITKKYSSAKKQLDSSPATACTGALA

>XP_028771214.1 blue-light photoreceptor PHR2 [Prosopis alba]

MDSNLPVAENQDLKVEEEQNPHAVVPSKETAVPPFATASLSLSLSTILPSHFFAQPKTSALFSAHPNKVKVPTQASSLTHLSLSTTSSPPPSKISFKSTISANPLQNPLTLGPHRPLDPSNGAAIRRTSIVWFRNDLRVHDNECLNSANNESMSVLPVYCFDPRDYGKSSSGFDKTGPYRATFLIESVSNLRKNLQARGSDLVVRVGKPETVLVEIAKVIGADAVYAHREVSHDEVKAEERIEAALKEENVEVKYFWGSTLFHVDDLPFKLEDMPTNYGGFREKLQKLEIRKTIEALDQLKGLPSRGDVEPGDIPSLVDLGLNPSAAMAQNGKSAANTSMVGGETEALQKLRKFAAECEAQPQKGLADGKQGSIYGANFSCKISPWLAMGCLSPRTMFDELKKTAGRTISASSSQNNGGGSISNNGTNWLMFELLWRDFFRFITKKYSSAKQQREGAPATACTGALA

>KAB1202325.1 Blue-light photoreceptor PHR2 [Morella rubra]

MDSHGQILENPETKSLEEQNSLAIVTAKDQTTSPAFATASLSLSLSTILPTPFSIQPKIYSLFSHQPSKVKVPTQASSLAHLSLSATSPSPSKLSFKSTVSANPLQNPLSLGPRRPSDPSNGAGIRRAAIVWFRNDLRVSDNECLNTANNEAMSVLPVYCFDPRDYGKSTSGFDKTGPYRASFLIESVSDLRKNLQSRGSDLVVRIGKPETVLVELAKAVGADAVYAHREVSHDEVKSEGKIEAAMKEDNVEVKYSWGSTLYHLDDLPFKLEDMPANYGGFREKVQGLEVRKTIEALDQIKGLPARGDVEPGDIPSLMDLGLNPSATISQALPAAAKWYCVGSTWDGKPAANASMVGGETEALQRLKKFAAECQAQPPKGNNDSIYGANFSCKISPWLAMGCLSPRSMFDELKKTATRTISATSNQNDGGSPDSGNNWLMFELLWRDFFRFITKKYSSKQLEAAPATACTGALA

>XP_030483036.1 blue-light photoreceptor PHR2 [Cannabis sativa]

MDPDSKMENPETQSSEEQNSMAVVVPSSSGETRLPPFATASLSLSLSTILPTHFFQQSKISTLFSQPNKAKVPTQASSLTHLSLSAASVSQQPKLSFKSTISANPLQNPLSLGPRRPMDPSNGAAVRRASIVWFRNDLRVHDNECLNSAHNESMSVLPVYCFDPRDYGKSSSGFDKTGPYRATFLVESVKDLRKNLQARGSDLVVRIGKPETVLVDLAKAIGADAIYAHREVSHDEVKTEERIEAAMKDENFEVKYFWGSTLYHADDLPFKLEDMPSNYGGFREKVQGLEVRQTIEALEQMKGLPSRGDVEPGDIPSLMDLGVNQSATLTQDGKPAANASLVGGETEALERLKKFAAECQAQPPKGGKDGSHDSIYGANFSCKISPWLAMGCLSPRSMFDELKKTTSRTISASSNKNDDGSGMNWLLFELLWRDFFRFITKKYSSAKKQLDNAPATACTGAFA

>XP_050111085.1 blue-light photoreceptor PHR2-like [Malus sylvestris]

MDANRQIPENPESKSPEEQNPLAIVPSAAGLSPFATASLSLSLSTILPTHFFQQPKVSTLFSSQPTKVKVPTQASSLTHLSLSTANVTPPRLSFKSTIAANPLQSPLSLGPRRPLDPSNGAAIRRASIVWFRNDLRVHDNECLNSANNESVSVLPVYCFDPRDYGKSSSGFDKTGPYRATFLVESVADLRKNLQARGSDLVVRIGKPETVLVELAKAIGADAIYAHREVSRDEVKEEEKIEAAMKEENVEVKYFWGSTLYHAEDLPFKLEDMPTKYGDFREKVKGLEVRKTIEALDQMKGLPSRGDVEPGDVPTLMDLGLNPSATTSQDGRPAAIASVVGGETEALERLKKYAAECQAQPPKASKDGKHDSIYGANFSCKVSPWLVLGCLSPRSMFDELKKTSSRTISASSNRNDDGGSGMNWLMFELLWRDFFRFVTCSSTKKQLNAAPATACTGALA

>PON65891.1 DNA photolyase class 1, 8-HDF type [Parasponia andersonii]

MDPDSKMEKNTETQSPEEQNSLAIIVASSPFATASLSLSLSTVIPTHFFQQPKISTLFSQPNKVKVPTQASSLSHLSLSSASVTPPPKLSFKSTISANPLQNPLSLGPRRPMDPSNGAAVRRASIVWFRNDLRVHDNECLNSAHNESMSVLPVYCFDPRDYGKSSSGFDKTGPYRATFLVESVTDLRKNLQARGSDLVVRVGKPETVLAELAKAIGADAVYAHREVSHDEVKAEERIEAAMKEENIEVKYFWGSTLYHVDDLSFKLEDMPSNYGGFREKVQGLDVRKTIEALDQMKGLPSRGDVEPGDIPSLMDLGINPSATMAQDGKPAANASLVGGETEALERLKKFAAECQAQPHKGSKDGSHDSIYGANFSCKISPWLAMGCLSPRSMFDELKKTTTRTISASSNINDGGSGMNWLLFELLWRDFFRFITKKYSSAKKQLDSSPATACTGALA

>XP_008359419.2 blue-light photoreceptor PHR2-like [Malus domestica]

MDANRQIPENPESKSPEEQNPLAIVPSAAGLSPFATASLSLSLSTILPTHFFQQPKVSTLFSSQPTKVKVPTQASSLAHLSLSTTANVTPPKLSFKSTIAANPLQSPLSLGPRRPLDPSNGAAIRRASIVWFRNDLRVHDNECLNSANNESVSVLPVYCFDPRDYGKSSSGFDKTGPYRATFLVESVADLRKNLQARGSDLVVRIGKPETVLVELAKAIGADAIYAHREVSRDEVKEEEKIEAAMKEENVEVKYFWGSTLYHAEDLPFKLEDMPTKYGDFREKVKGLEVRKTIEALDQMKGLPSRGDVEPGDVPTLMDLGLNPSATTSQDGRPAAIASVVGGETEALERLKKYAAECQAQPPKASKDGKHDSIYGANFSCKVSPWLVLGCLSPRSMFDELKKTSSRTISASSNRNDDGCSGMNWLMFELLWRDFFRFVTCSSTKKQLNAAPATACTGALA

>XP_018815193.2 blue-light photoreceptor PHR2-like [Juglans regia]

MDPNTQILENPEAKSLEEQNPLAIVPSKEQITSPPFATASLSLSLNTILPTPFSLQPKIYSLFSHQPSRVKVPTQASSLAHLSLTATSPSPSKLSFKSTVSANPLQNPLSLGPRRPSDPSNGAGIRRAAIVWFRNDLRVHDNECLNTASNEAMSVLPVYCFDPRDYGKSSSGFDKTGSNRASFLIESVSDLRRNLQAKGSDLVVRIGKPETVLVELAKAVGADAVYAHREVSHDEVKAEGRIEAAMKEESVEVKYFWGSTLYHLDDLPFKLEEMPANYSGFREKVQGLEVRKTIEALDQVKGLPARGDVEPGDIPSLVDLGLNPSATMAQDGKLAANASMVGGETGALQRIKRIAAECQAQPHKETKDGNDIYGANFSCKISPWLAMGCLSPRSMFDELRKTATRTISASTNRNDGGSPDTGNNWLMLELMWRDFFRFVTKKYSSKRLEAAPATASTGALA

>XP_027363396.1 blue-light photoreceptor PHR2 [Abrus precatorius]

MDSNHQIAAETEKDEQNQNPTHSTEPPFAVASLSLTLSTVLPFVQPKIPSFFSRTQPNKLKFPPTQASSLTHLSLSTPAPTKTSFKSNLSANPLHTPHSLGPHRPLDPSNAAALRRASIVWFRNDLRLHDNECLNAANNESLSVLPLYCFDPSDYGKSSSGFDKTGPYRASFLIQSVSDLRRNLQARGSDLVVRVGKPETVLVELAKAVGADAVYAHREVSHDEVKTEEKIEAAMKEENVEVKYFWGSTLYHVDDLPFKLEDMPSNYGGFRDRVQKLEIRNTIEALDQLKGLPSRGDVEPGDIPSLMDLGLNPSPTMPQDGKPAANASMVGGETEALQRLKRFAAECEAQPHKGSKDGTQSIYGANFSCKISPWLAMGCLSPRTMYDELKKTVSRSISASSNRNDGGSGSSKTGTNWLMFELLWRDFFRFITKKYSSAKKQLEATPATACTGALA

>XP_044502252.1 blue-light photoreceptor PHR2-like [Mangifera indica]

MDPNLKTLENSENEEQNPLAIVRSQSPFATLSLSFSLPKVLPTNSFFLQPKISSLFSHQPCKVKVPTQASSLSHLSLSSSASLAPTKISFKSTISANPLQNPLTLGPLRPLDPNNGAGIRRAAICWFRNDLRVHDNECLNTANNESMSVLPVYCFDPRDYAKSSSGFDKTGPYRASFLIESVSDLRKNLQARGSDLVVRIGKPEDVLVELAKAIGADAVYAHREVSHDDVKSEDKIEAAMKEEGVEVKCFWGSTLYHIDDLPFKLEEMPTTYGGFREKAQGLEVRKTIEALDQMKGLPSRGDVEPGDIPSLMDLGLNPSAAMSQDGKPAANSLVGGETEALQRLKKFAAECQAQPPQGSKGGSHDSIYGANFSCKISPWLTVGCISPRSMFDELKKTAASISASSNRNGGASGSSDTGGNWLMFELLWRDFFRFITKKYSSAKKVVEAAPATACMGALA

>GMI66595.1 photolyase/blue-light receptor 2 [Hibiscus trionum]

MDSNTQSRENPEIESTEQQSQTPNSVSQSPFATASLSLSSLPTTLPTQFFIQPKILSLFSAQSPTKVKVPTQASSLSNLSLSSTSPSPSKFSFKSTFANNPLQSPLSLGPRRPLDPSNGAALRRASIVWFRNDLRVHDNECLNTASNESMSVLPVYCFDPRDYGKSSSGFDKTGPYRATFLIESVSDLRKNLQARGSDLVVRIGKPESVLVELAKAIGADAIYAHREVSHDEVKAEEKIESAMKEEGVEVKYFWGSTLFHVDDLPFKLEDMPSNYGGFKEKVKGLEIRKTIEALDQMKGMPSRGDVETGDIPSLTDLGLNPTATMAQDGRQSVSATMAGGENEAMQRLKKFAAECQAQPYKGSKDGSQGSIYGANFSCKISPWLAMGCISPRFMFDELKKTVNRTISATSKKNDGGSGSPDTQMNWLMYELLWRDFFRFITKKYSCAKVGSAAPATACTGALA

>XP_024164953.1 blue-light photoreceptor PHR2 [Rosa chinensis]

MDPNSQISENPESSEEQQQQQQNPLALLLPSSSSPFATASLSLSLASILPTHFFQQPKVATLFSSQPTKAKIPSQASSLAHLSLSASANVATPSKLSFKSTISANPLQNPLTLGPRRPLDPNNGAGIRRASIVWFRNDLRVHDNECLNSANNESMSVLPVYCFDPRDYGKSSSGFDKTGPFRAQFLVESVSDLRKNLQARGSDLVVRIGKPETVLAELAKTIGADAIYAHREVSHDEVKSEERIESAMKDENVEVKYFWGSTLYHMEDLPFKLEDMPTNYGGFREKVKGLEVRKTIEALEQLKGMPSRGDVEPGDVPSLMDLGLNPSASMAQDGKPGAASMVGGEAEALERLKKFAAECQAQPPKGSKEGSQNSIYGANFSCKISPWLAMGCLSPRSMFDELKKTASRSVSASSNPDDGGSGTNWLMFELLWRDFFRFVTKKYSAGKKQLDASPATACTGALA

>XP_044492929.1 blue-light photoreceptor PHR2 [Mangifera indica]

MDPNFKTLENSANDEQNPLAIAPSQSPFATLSLSFSLSKVLPNNSFFLQPKISSLFSHQPSKVKVPTQASSLSHLSLSSSSASLTPTKISFKSTISANPLQNPLTLGPHRPLDPNNGAGLRRAAICWFRNDLRVHDNECLNTANNESMSVLPVYCFDPRDYGKSSSGFDKTGPYRASFLIESVSDLRKNLQARGSDLVVRLGKPENVLVELAKAIGADAVYAHGEVSHDEVKCEEKIEAAMKEEGVEVKYFWGSTLYHVDDLPFKLEEMPTNYGGFREKVQGLEVRKTIEALDQMKGLPSRGDVEPGDIPSLMDLGLNPSAAMSQGGKPAANSMVGGETEALQRLKKFAAECQAQPHKGTKDGSHDSVYGANFSCKISPWLAMGCISPRSMFDELKKTTTSISASSNWNGNGSGSSDSGGNWLMFELLWRDFFRFITKKYSSAKKAVEAAPATACTGALA

>KAF7818196.1 blue-light photoreceptor PHR2 [Senna tora]

MDVNHQIVESEDPKSGEETNVPPPFATASLSLSLSTILPSHFFVQPKTPPSFSSLPKVKIPTQASSLAHLSLSTASSPSPSKLSFKSTISANPLHNPLSLGPHRPLDPSNGAVIRRASIVWFRNDLRVHDNECLNSAHNESMSVLPVYCFDPRDYGKSSSGFDKTGPYRATFLIESVSNLRKNLQARGSDLVVRIGKPETVLVELAKTIGADAVYAHREVSHDEVKAEERIEAAMKEENVELKYFWGSTLYHVDDLPFKLEDMPTNYGGFREKVHKLEIRKTIEALDQLKGIPSRGDVEPGDIPSLMDLGLNPSATMSQDGKSAANSSMVGGETEALQRLRKFAAECEAQPHKGFKDGKQDSIYGANFSCKISPWLAMGCLSPRTMFDELKKTASRMVSASSSQNNGGSGVSNNGTNWLMFELLWRDFFRFITKKYSSAKKQLEGAPATACTGALA

>XP_031261651.1 blue-light photoreceptor PHR2 [Pistacia vera]

MDPNLQTLENSENEEQNLLAIVPSQSPFATLSLSFSLSKVLPNNSFFLQPKISSLFSHQPSKVKVPTQASSLSHLSLSSSSASLAPTKISFKSTISANPLQNPLTLGPHRPLDPNNGAGIRRAAICWFRNDLRVHDNECLNTANNESMSVLPVYCFDPRDYGKSSSGFDKTGPYRASFLVESVSDLRKNLQARGSDLVVRVGKPENVLVELAKAIGADAVYAHREVSHDEVKSEDKIEAAMKEEGFEVKYFWGSTLYHVDDLPFKLEEMPTNYGGFREKVQGLEVRKTIEALDQMKGLPSRGDVEPGDIPSLMDLGLNPSAAISQDGKPAANSMVGGETEALQRLKRFAAECQAQPHKGSKDGSHDSIYGANFSCKISPWLAMGCISPRSMFDELKKTATSISASSNRNGGGSGSSDSGGNWLMFELLWRDFFRFVTKKYSSAKKVVEAAPATACTGALA

>XP_057963902.1 blue-light photoreceptor PHR2 isoform X1 [Malania oleifera]

MDPNPKTLENPEGKTPEDQEQLAVVPSNPSIAPPLATASVSLSLSAILPNNLLLSPKISSIFAPQPTKVKIPSQVSTLSQLSLFAALSPPKPFFKSTPISANPLQNPLSLGPRRTSDPSNGAGIRRASIVWFRNDLRVHDNECLNSANNESVSVLPVYCFDPRDYGKSSSGFDKTGPYRAAFLIESVSNLRRNLREKGSNLVVRIGKPETVLVELAKTVGADAVYAHREVSHDEVKAEEKIEAAMKEEGVEVKYFWGSTLYHVDDLPFKLEDMPSNYGGFREKVQGLEVRKTMAALDQLKRLPSRGDVEPGEIPSLLDLGLNQSATMPQDGKQAANASMVGGETEALQRLKKFAAECQAQPHKGTNSSENSIYGANFSCKISPWLAMGCLSPRSMFDELKKTACSRTISARKDNGGPSDPGMNWLMYELLWRDFFRFITKKYSSAKRQLDVAPATACTGALA

>XP_057963903.1 blue-light photoreceptor PHR2 isoform X2 [Malania oleifera]

MDPNPKTLENPEGKTPEDQEQLAVVPSNPSIAPPLATASVSLSLSAILPNNLLLSPKISSIFAPQPTKVKIPSQVSTLSQLSLFAALSPPKPFFKSTPISANPLQNPLSLGPRRTSDPSNGAGIRRASIVWFRNDLRVHDNECLNSANNESVSVLPVYCFDPRDYGKSSSGFDKTGPYRAAFLIESVSNLRRNLREKGSNLVVRIGKPETVLVELAKTVGADAVYAHREVSHDEVKAEEKIEAAMKEEGVEVKYFWGSTLYHVDDLPFKLEDMPSNYGGFREKVQGLEVRKTMAALDQLKRLPSRGDVEPGEIPSLLDLGLNQSATMPQDGKQAANASMVGGETEALQRLKKFAAECQAQPHKGTNSSENSIYGANFSCKISPWLAMGCLSPRSMFDELKKTACRTISARKDNGGPSDPGMNWLMYELLWRDFFRFITKKYSSAKRQLDVAPATACTGALA

>RDX90692.1 Blue-light photoreceptor PHR2, partial [Mucuna pruriens]

MDSTHQTEAEKKEEQKAPLPTEAPFAVASLSLSLSTIFPHIQPKIPSAPAPSKVKVPTQASSLTHLSLSTASPPPSKTSFKSTLSATPLHAPLSLGPHRPRDPSNTAALRRAAVVWFRNDLRLHDNECLAAANNESLSVLPVYCFDPADYGKSASGFDKTGPYRATFLIESVSDLRRSLQSRGSDLVVRVGKPETVLVELAKAVGADAVYAHREVSHDEVKAEEKVEAAMKEENVEVKYFWGSTLYHMDDLPFKLEDMPSNYGGFRDRVQKLEIRKTIEALDQLKGMPSRGDVEPGDIPSLMDLGLNPSPTMPQDGKFAANACMVGGETEALQRLKRFAAECEAQPNKGSKDGTQSIYGANFSCKISPWLAMGCLSPRTMYDELKKTASSVISASSNRNDGGNGSTKTGSNWLMFELLWRDFFRFITKKYSSAKKQEQLEAAPATACAGCGLLPSADMAVGCWMNKDEFRDCYFSYEE

>XP_004495945.1 blue-light photoreceptor PHR2 [Cicer arietinum]

MEPTNQTPPQEKENPFPTPPSEPQLPPSIASLSLSLSSILPLFQPKIPSLLSPPQPNKLKIPTQASSLTHLSLSTKTQSPPSKNKSSFKSTLSANPLKSPLSLGPHRPVDPSAAAAFRRTTIVWFRNDLRVHDNECLNTANNESISVLPVYCFDPADYGKSSSGFDKTGPFRATFLIESVSDLRKNLKARGSDLVVRVGKPETVLVELAKEVGAESVFAHREVSHDEVKMEEKIESKMKEENVEVKYFWGSTLYHVDDLPFKLEDMPSNYGGFRDRVQKLEVRKSIEALDQLKGLPSRGDVQPGDIPTLMDLGLNPSATMSQDGKPGPNASMAGGETEALQKLKRFAAECEAQPHKGSKDGAQDSIYGANFSCKISPWLAMGCLSPRAMYDELKKTASRAVSASSSRNDGGSGSSKTGTNWLMFELLWRDFFRFITKKYSSTKKQLEAAPATACTGALA

>NP_567341.1 CRY1 [Arabidopsis thaliana]

MSGSVSGCGSGGCSIVWFRRDLRVEDNPALAAAVRAGPVIALFVWAPEEEGHYHPGRVSRWWLKNSLAQLDSSLRSLGTCLITKRSTDSVASLLDVVKSTGASQIFFNHLYDPLSLVRDHRAKDVLTAQGIAVRSFNADLLYEPWEVTDELGRPFSMFAAFWERCLSMPYDPESPLLPPKKIISGDVSKCVADPLVFEDDSEKGSNALLARAWSPGWSNGDKALTTFINGPLLEYSKNRRKADSATTSFLSPHLHFGEVSVRKVFHLVRIKQVAWANEGNEAGEESVNLFLKSIGLREYSRYISFNHPYSHERPLLGHLKFFPWAVDENYFKAWRQGRTGYPLVDAGMRELWATGWLHDRIRVVVSSFFVKVLQLPWRWGMKYFWDTLLDADLESDALGWQYITGTLPDSREFDRIDNPQFEGYKFDPNGEYVRRWLPELSRLPTDWIHHPWNAPESVLQAAGIELGSNYPLPIVGLDEAKARLHEALSQMWQLEAASRAAIENGSEEGLGDSAEVEEAPIEFPRDITMEETEPTRLNPNRRYEDQMVPSITSSLIRPEEDEESSLNLRNSVGDSRAEVPRNMVNTNQAQQRRAEPASNQVTAMIPEFNIRIVAESTEDSTAESSSSGRRERSGGIVPEWSPGYSEQFPSEENGIGGGSTTSSYLQNHHEILNWRRLSQTG

>NP_171935.1 CRY2 [Arabidopsis thaliana]

MKMDKKTIVWFRRDLRIEDNPALAAAAHEGSVFPVFIWCPEEEGQFYPGRASRWWMKQSLAHLSQSLKALGSDLTLIKTHNTISAILDCIRVTGATKVVFNHLYDPVSLVRDHTVKEKLVERGISVQSYNGDLLYEPWEIYCEKGKPFTSFNSYWKKCLDMSIESVMLPPPWRLMPITAAAEAIWACSIEELGLENEAEKPSNALLTRAWSPGWSNADKLLNEFIEKQLIDYAKNSKKVVGNSTSLLSPYLHFGEISVRHVFQCARMKQIIWARDKNSEGEESADLFLRGIGLREYSRYICFNFPFTHEQSLLSHLRFFPWDADVDKFKAWRQGRTGYPLVDAGMRELWATGWMHNRIRVIVSSFAVKFLLLPWKWGMKYFWDTLLDADLECDILGWQYISGSIPDGHELDRLDNPALQGAKYDPEGEYIRQWLPELARLPTEWIHHPWDAPLTVLKASGVELGTNYAKPIVDIDTARELLAKAISRTREAQIMIGAAPDEIVADSFEALGANTIKEPGLCPSVSSNDQQVPSAVRYNGSKRVKPEEEEERDMKKSRGFDERELFSTAESSSSSSVFFVSQSCSLASEGKNLEGIQDSSDQITTSLGKNGCK

>NP_568461.3 CRY3 [Arabidopsis thaliana]

MAASSLSLSSPLSNPLRRFTLHHLHLSKKPLSSSSLFLCSAAKMNDHIHRVPALTEEEIDSVAIKTFERYALPSSSSVKRKGKGVTILWFRNDLRVLDNDALYKAWSSSDTILPVYCLDPRLFHTTHFFNFPKTGALRGGFLMECLVDLRKNLMKRGLNLLIRSGKPEEILPSLAKDFGARTVFAHKETCSEEVDVERLVNQGLKRVGNSTKLELIWGSTMYHKDDLPFDVFDLPDVYTQFRKSVEAKCSIRSSTRIPLSLGPTPSVDDWGDVPTLEKLGVEPQEVTRGMRFVGGESAGVGRVFEYFWKKDLLKVYKETRNGMLGPDYSTKFSPWLAFGCISPRFIYEEVQRYEKERVANNSTYWVLFELIWRDYFRFLSIKCGNSLFHLGGPRNVQGKWSQDQKLFESWRDAKTGYPLIDANMKELSTTGFMSNRGRQIVCSFLVRDMGLDWRMGAEWFETCLLDYDPCSNYGNWTYGAGVGNDPREDRYFSIPKQAQNYDPEGEYVAFWLQQLRRLPKEKRHWPGRLMYMDTVVPLKHGNGPMAGGSKSGGGFRGSHSGRRSRHNGP

>NP_849651.1 PHR1/UVR2 [Arabidopsis thaliana]

MASTVSVQPGRIRILKKGSWQPLDQTVGPVVYWMFRDQRLKDNWALIHAVDLANRTNAPVAVVFNLFDQFLDAKARQLGFMLKGLRQLHHQIDSLQIPFFLLQGDAKETIPNFLTECGASHLVTDFSPLREIRRCKDEVVKRTSDSLAIHEVDAHNVVPMWAASSKLEYSARTIRGKINKLLPDYLIEFPKLEPPKKKWTGMMDKKLVDWDSLIDKVVREGAEVPEIEWCVPGEDAGIEVLMGNKDGFLTKRLKNYSTDRNNPIKPKALSGLSPYLHFGQVSAQRCALEARKVRSTSPQAVDTFLEELIVRRELSDNFCYYQPHYDSLKGAWEWARKSLMDHASDKREHIYSLEQLEKGLTADPLWNASQLEMVYQGKMHGFMRMYWAKKILEWTKGPEEALSISIYLNNKYEIDGRDPSGYVGCMWSICGVHDQGWKERPVFGKIRYMNYAGCKRKFNVDSYISYVKSLVSVTKKKRKAEEQLTRDSVDPKITIV

>NP_566520.1 UVR3 [Arabidopsis thaliana]

MQRFCVCSPSSYRLNPITSMATGSGSLIWFRKGLRVHDNPALEYASKGSEFMYPVFVIDPHYMESDPSAFSPGSSRAGVNRIRFLLESLKDLDSSLKKLGSRLLVFKGEPGEVLVRCLQEWKVKRLCFEYDTDPYYQALDVKVKDYASSTGVEVFSPVSHTLFNPAHIIEKNGGKPPLSYQSFLKVAGEPSCAKSELVMSYSSLPPIGDIGNLGISEVPSLEELGYKDDEQADWTPFRGGESEALKRLTKSISDKAWVANFEKPKGDPSAFLKPATTVMSPYLKFGCLSSRYFYQCLQNIYKDVKKHTSPPVSLLGQLLWREFFYTTAFGTPNFDKMKGNRICKQIPWNEDHAMLAAWRDGKTGYPWIDAIMVQLLKWGWMHHLARHCVACFLTRGDLFIHWEQGRDVFERLLIDSDWAINNGNWMWLSCSSFFYQFNRIYSPISFGKKYDPDGKYIRHFLPVLKDMPKQYIYEPWTAPLSVQTKANCIVGKDYPKPMVLHDSASKECKRKMGEAYALNKKMDGKVDEENLRDLRRKLQKDEHEESKIRNQRPKLK

>XP_001701871.1_cry_DASH1 [Chlamydomonas_rei]

MATAGGGQRLVLWFRNDLRLHDNYIVHEAAQRVKRGEASEVLPVYVYDPRFFAATPWGALKTGAHRAKFIQECVADLRQRLQGLGSDLVVAVGQPEQLLPALLEGGGAAPLVLTAEEVTSEEAAVDVAVARAIKPAGGKLLRYWGHTMYHYDDLDLPDVFTPFKEKVEKRAALPLPPAPRLGATAAAALAAPLPGWEQLPPAPPVTHPKAVLDFKGGETAALARLKYYLWDSDLLSTYFDTRNGMLGGDYSTKFAPWLAQGCISPRKIFHEIRKYESQRFSNKSTYWVIFELIWRDFFRFFALKHGNRIFFETGTSGLPLVWNPDPELWARWREGRTGLPLVDANMRELAATGFMSNRGRQNVASYLVLDLGVDWRRGADYFEEVLLDYDVTSNWGNWVAAAGLTGGRVNHFNIAKQSKDYDPTGEYVKTWCPELKNVPVTKVHEPWLMSKEEQERSGCRIGVDYPNPI

>OUS42345.1_cry_DASH [Ostreococcus_tau]

MGRTRVVIWFRNDLRLLDNACVARAATLASESSDVEVVPVYVFDETYFKPSKRGLARFGAGRGKFTLECVGDLKTSLRALGSDLLVRCGKSRDVIAELTLTGANDRTIILTQTEVTSEETEMDVAVERATRERARGGAASATMERHWGSTLYHIDDVPFDVTSGLSDLPDVFTPFRNKVESKCKVRDVIPAPTANALGHVPASVEGFEWMPDPSDLPFASSEIAMDCDKRIKDCLDERSVLDFKGGESNALARVKYYLWESDRLATYFETRNGMLGGDYSTKLAPWLALGCVSPRHVVSEIRRYESERVENKSTYWVIFELIWRDFFKFFALKHGNKIFHLDGTAGRRASWKRDEKILKAWKTGTTGYPLIDANMRELAATGFMSNRGRQNVASWLALDAGIDWRHGADWFEHHLLDYDTASNWGNWCAAAGMTGGRINRFNIAKQTKDYDPAGEYIKTWVKELAEVPAAYIADPNQAPRELRDRIGLNYPNKLALPRRDFTEMGSPPGPR RGGGGGGRGRGRPGGSTPNRGTKARVASVYDTVYG

>NP_194259 At4g25290 [Arabidopsis thaliana]

MAFLALPHFLHLRLRRNDRKNRCKCCLSSATNEGSTAVVWFKHDLRVDDHPGLLAASKHRAVIPLYVLDRRILSRYTTDTLELAIIALEDLRKTLKKQGSNLMLRYGNAENVIEDLVKEVRAPFVFVEEEVEYHLCEVLDAVKNKLEGVSLSGESPRIVAWRTPFYESQNLTDLPQSWEEFKKLKLPLTLPVPAAKFSSPGSELQWGSVPTLDDLKDYLKESLWEIENSWREMAQASAERVLMERLGNLKESSMEPIVDGSLGKKVDNSVFVTSKRDTVGGGNEVVLNALAGYLRYLEGTSRDDWQEVHARLRDAETRPGASFFKLFGPVLCLGIVSRRSVHYEAIEYEKERNAGFISPFGYSAATVSAATDAVCSMEWYYLLALSRERIDEKRHAIRIWRWKGYLIQYTVVGNEGPAVLLVHGFGAFLEHYRDNVDNIVNSKNRVWTITVLGFGKSEKPNIIYTELLWAELLRDFMAEVVGEPAHCVGNSIGGYFVALMAFLWPALVKSVVLVNSAGNVVPGYSPLPISRERRVPFGAQFGSRLLLFFLQLNVKKLLKDCYPVKPERADDFLVTEMLRASRDPGVVMVLESIFGFDLSLPLNYLLKGFEEKTLVIQGMEDPISDPQKKVALLKELCPAMVIKKVKAGHCPHDEISEEVNPIICEWIVKVTNDDRELKASSSQQLYHSNKQN

>NP_001296304.1 cryptochrome DASH, chloroplastic/mitochondrial [Solanum lycopersicum]

MIKQPFLLTKFTPFSSKSKHTLFTFHCNFSIKMASLTARTTPTVQNVPGLTPEEMERVCEQTFQRYESGGLGKRKGKGVAIVWFRNDLRVLDNEALLRAWVSSEAILPVYCVDPRLFGTTHYFGMPKTGALRAQFIIECLNDLKRNLVKRGLDLLIQHGKPEDIVPSLAKAYKAHTVYAHKETCSEEVKVEKMVTRNLQKLVSPSSGGIGNDPGSGNTTKLELVWGSTMYHIDDLPFDCESLPDVYTQFRKSVEYKSKVRNCTKLPTSFGPPPEVGDWGHVPQVSELGLQQEKVSKGMNFVGGESAALGRVHDYFWKKDLLKVYKETRNGMLGADYSTKFSPWLASGSLSPRFIYEEVKRYEKERLSNDSTYWVLFELIWRDYFRFLSIKLANLLFQAGGPQKVNINWSQDQTMFDAWRRGQTGYPLIDANMKELAATGYMSNRGRQIVCSFLVRDMGIDWRMGAEWFETCLLDYDPCSNYGNWTYGAGVGNDPREDRYFSIPKQAQNYDPEGEFVAYWLPELRALPREKRHSPGMMYLNPIVALKHGYTKKTGDSKTAFSSRRGRPEDNRRKRHGY

>XP_002178889.1 cry-dash from the cryptochrome/photolyase family [Phaeodactylum tricornutum CCAP 1055/1]

MSSSRSKQSFWFLIFVHTVLILTFAAGAATAMRGASVSTVTKATSRASSQVVLHWFRHGDLRLLDNPALIHSSKTAESCVPVFCFDDSVYGNDNRTPDTRAPHSNDRGQLKCGPRRAQFVLDSVQDLRRSLQSRGSALYVAHGKPAQVFQRLVDAWPAVPAADTAAPNGSLLTIVCQREVVREENDAVRAVQSVLRRRFPQAKVQQIWGSTMYELDDLPFATDLANMPDTFTPFRNKVEKNCQIGTPLPVPKQLSLPENFPSALKQGLEYLPTLKELGYTDAQIQQVETHDERGVLHFMVAKQPDSHDCLKDYFETRNGMLGPNYSTKFSPWLAHGNVSPRYVAAQCRKYEEERVENKSTYWVVFELLWRDFCKFFATKHGDAIFYPYGTTERTDRHKPWSTFGRNLQAWQEGRTGYPLVDANMRELVATGFMSNRGRQNVASFLAINLNHDWRCGGDFFESHLLDYDVYSNWVNWCAAAGMTGGRLNRFNISKQSKDYDQHGDYVRHWLPELAKVPNEFVHEPWKMTSFQQMEYECKLGVDYPNPIVPPSRPNPHTDRNNRGRGGHQSKGNNRHGPPDKSRKANASGSNRHQKYEMKSLQPGSFRVKES

>NP_991249.1 cryptochrome DASH [Danio rerio]

MSASRTVICLLRNDLRLHDNEVFHWAQRNAEHIIPLYCFDPRHYQGTYHYNFPKTGPFRLRFLLDSVKDLRALLKKHGSTLLVRQGKPEDVVCELIKQLGSVSTVAFHEEVASEEKSVEEKLKEICCQNKVRVQTFWGSTLYHRDDLPFSHIGGLPDVYTQFRKAVEAQGRVRPVLSTPEQVKSPPSGLEEGPIPTFDSLGQTEPLDDCRSAFPCRGGETEALARLKHYFWDTNAVATYKETRNGMIGVDFSTKFSPWLALGCISPRYIYEQIKKYEVERTANQSTYWVIFELLWRDYFKFVALKYGNRIFYMNGLQDKHVPWKTDMKMFDAWKEGRTGVPFVDANMRELALTGFMSNRGRQNVASFLTKDLGLDWRLGAEWFEYLLVDHDVCSNYGNWLYSAGIGNDPRENRKFNMIKQGLDYDNNGDYVRQWVPELRGIKGGDVHTPWTLSNSALSHAQVSLNQTYPCPIITAPEWSRHVNNKSSGPSSSKGRKGSSYTARQHKDRGIDFYFSKNKHF

>OP889330 Blue-light_photoreceptor_PHR2 [Chrysanthemum morifolium]

MESSSTKKPQDTTTESNQDSDQTQNQQEDQLTINPIPIFTSITLSFPSFIPTKHLTPTPKHTPKPLFKLPNTTLSSLSHNLTLISSSIPTKPLIKPTITLNPLQNPLSLNPRRPSDPFNSAAARRTTIVLFRNDLRCHDNEALVSANNESTSVLPVYCFDPRDYGKSKSGFDKTGPYRASFLIESVSNLRKNLQARGSDLVVRIGKVESVLSELVKAVGAEVVYAHREVSHDEVKCEENIEARLKDEGVEVKYFWGSTLYHIDDLPFKLDEIPTNYGGFREKVKGLKIRKVVEVVDQFRGLPVAGDVEVGEIPSLSDLGLTPTPTMNQAKATANASLVGGETEALERLKKFAAECQAKPHKDGSNDTNLYGANFSCKISPWLAMGCLSPRSMFDELKKSASRTISAASNQKDGGDTGMNWLIFITKKYSSAKQNNAAPVTACTGAAA
